# Supercentenarian and remarkable age records exhibit patterns indicative of clerical errors and pension fraud

**DOI:** 10.1101/704080

**Authors:** Saul Justin Newman

**Affiliations:** University College, University of Oxford, Oxford, UK; Social Research Institute, University College London, London, UK; Oxford Institute of Population Ageing, University of Oxford, Oxford, UK, City: London; Oxford Country: United Kingdom

## Abstract

The observation of individuals attaining remarkable ages, and their concentration into geographic sub-regions or ‘blue zones’, has generated considerable scientific interest. Proposed drivers of remarkable longevity include high vegetable intake, strong social connections, and genetic markers. Here, we reveal new predictors of remarkable longevity and ‘supercentenarian’ status. In the United States, supercentenarian status is predicted by the absence of vital registration. The state-specific introduction of birth certificates is associated with a 69-82% fall in the number of supercentenarian records. In Italy, England, and France, which have more uniform vital registration, remarkable longevity is instead predicted by poverty, low per capita incomes, shorter life expectancy, higher crime rates, worse health, higher deprivation, fewer 90+ year olds, and residence in remote, overseas, and colonial territories. In England and France, higher old-age poverty rates alone predict more than half of the regional variation in attaining a remarkable age. Only 18% of ‘exhaustively’ validated supercentenarians have a birth certificate, falling to zero percent in the USA, and supercentenarian birthdates are concentrated on days divisible by five: a pattern indicative of widespread fraud and error. Finally, the designated ‘blue zones’ of Sardinia, Okinawa, and Ikaria corresponded to regions with low incomes, low literacy, high crime rate and short life expectancy relative to their national average. As such, relative poverty and short lifespan constitute unexpected predictors of centenarian and supercentenarian status and support a primary role of fraud and error in generating remarkable human age records.

## Introduction

The concentration of remarkable-aged individuals within geographic regions or ‘blue zones’^1^ has stimulated diverse efforts to understand factors driving survival patterns in these populations^2,3^. Both the overall population residing within these regions, and the individuals exceeding remarkable age cut-offs, have been subject to extensive analysis of lifestyle patterns^2,4–6^, social connections^3,7^, biomarkers^8,9^, and genomic variants^10^, under the assumption that these represent potential drivers behind the attainment of remarkable age.

However, alternative explanations for the distribution of remarkable age records appear to have been overlooked. Previous work has noted the potential of population illiteracy^11^ or heterogeneity^12^ to explain remarkable age patterns. Other investigations have revealed the potential role of errors^13–16^, bad data^17^, and operator biases^18^ in generating old-age data and survival patterns. In turn, these findings prompted a response with potentially disruptive implications: that, under such models, the majority if not all remarkable age records may be errors^19^.

There is a theoretical reason to expect that most or all old-age records are errors^13,15^. Consider, for example, a population of fifty-year-olds into which we introduce a vanishingly low rate of random, undetectable age-coding errors. These rare errors involve taking a forty-year-old, and making their documents state they are now fifty years old.

The 40-year-old ‘young liar’ errors will have over double the annual survival rate of the (error-free) 50-year-old population. They are, after all, biologically 10 years younger. As the ‘young liar’ errors are more likely to survive than the error-free data, age-coding errors constitute an exponentially larger fraction of the population over time. Eventually, this exponential growth overtakes the population and, even from a vanishingly low baseline error rates of (say) 0.001%^15^, the entire population becomes ‘young liar’ errors at advanced ages^15^. Of course, the baseline error rate in most populations is not vanishingly low. The USA, which has more supercentenarian records than any other country, exhibits historical error rates of 17%-66%^20–23^, even in the general population^20^.

Here, we explore the possibility that extreme old-age data are dominated by age-coding errors and/or fraud by linking civil registration rates and indicators of poverty to per-capita estimates of remarkable age attainment, obtained from central population registries and validated supercentenarian databases, across the USA, France, Japan, England, and Italy.

These data reveal that remarkable age attainment is predicted by indicators of error and fraud, including illiteracy, poverty, high crime rates, short average lifespans, and the absence of birth certificates. As a result, these findings raise serious questions about the validity of an extensive body of research based on the remarkable reported ages of populations and individuals.

## Methods

The number and birthplace of all validated supercentenarians (individuals attaining 110 years of age) and semisupercentenarians (SSCs; individuals attaining 105 years of age) were downloaded from the Gerontology Research Group or GRG supercentenarian table^24^ (updated 2017) and the International Database on Longevity or IDL ^25^. These data were aggregated by subnational units for birth locations, which were provided for the IDL data, and obtained through biographical research for the GRG data. Populations were excluded due to incomplete subnational birthplace records (<25% complete) or countries with an insufficient number of provinces to fit spatial regressions (<15 total provinces), leaving population data on SSCs and supercentenarians in the USA, France, and England (Fig 1). These countries contained a substantial majority of global supercentenarians and were the best placed to evaluate population patterns.

**Figure 1.**
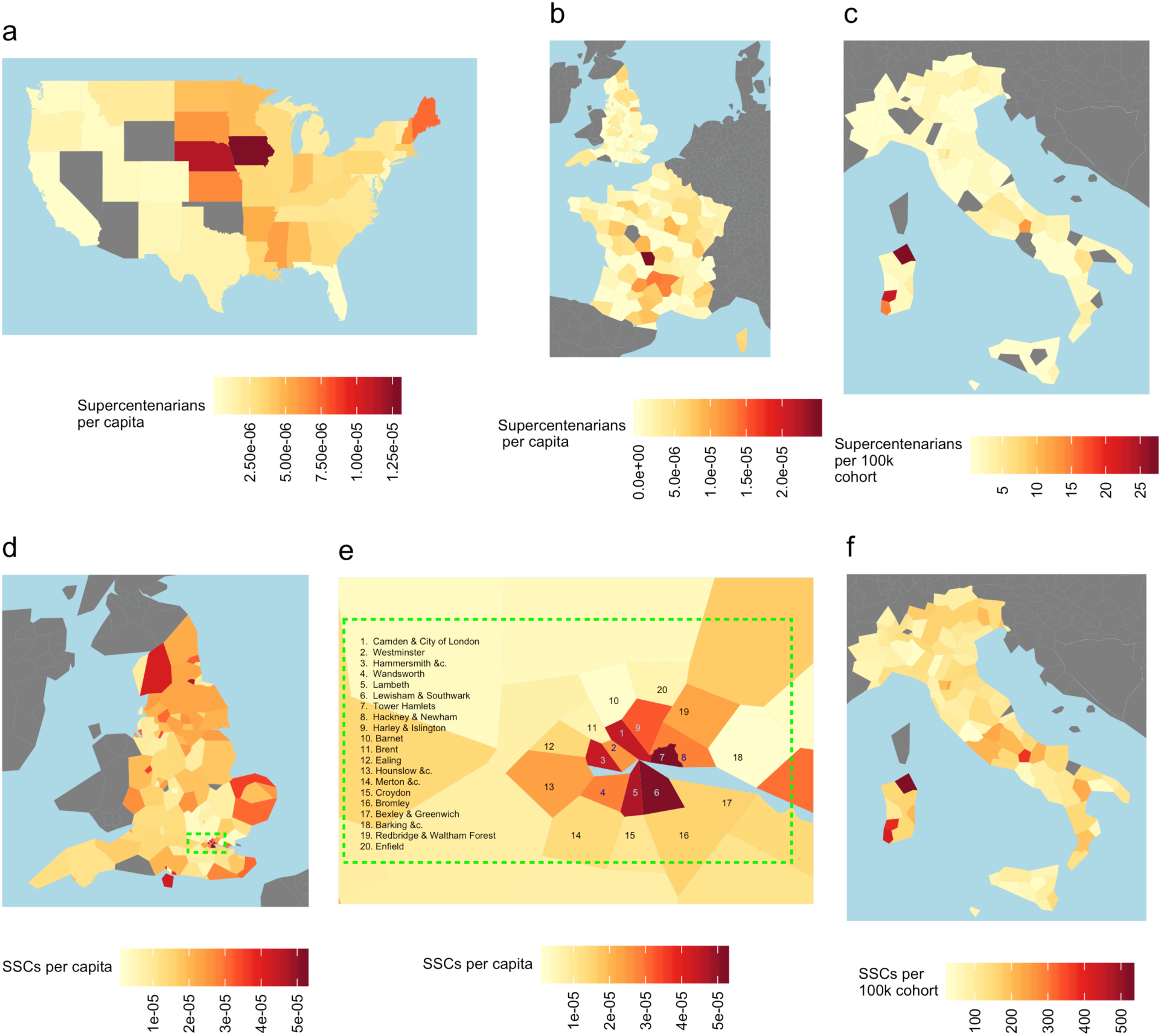
Regional distributions for the density of the oldest-old. A substantial majority of the oldest-old people are concentrated in a few countries exhibiting large regional variation in density of the oldest-old. Known supercentenarians are concentrated in the USA (a), with large numbers in France(b), England (b), and Italy (c; life table estimated rates). Marked variation exists in estimated SSC densities across England (d) – including (e) the 19-fold differences in SSC abundance across London (green box in d-e) between Tower Hamlets (region 7; most) and Barnet (region 10; least) – and across Italian provinces (f).

To quantify the distribution of remarkable-aged individuals in Italy, province-specific quinquennial life tables were downloaded from the Italian Istituto Nazionale di Statistica Elders.Stat database ^26^ to obtain age-specific (longitudinal) cohort survivorship data (Fig 1c,f; S1 Code). Using contemporary data across Italian provinces, probabilities of survival (l_x_) to ages 90-115, and life expectancy at age 100 were fit as dependent variables, and survival rates at age 55 and life expectancy at age 55 within the same cohort as independent variables, using a simple linear mixed model regression (S1 Code).

Japanese centenarian data were downloaded from the Japanese Ministry of Health, Labour, and Welfare^27^ through the Statistics Japan portal^28^ for all 47 prefectures (Fig S1), alongside data on prefectural income per capita (in 2011 yen), employment rates, age structure, survivorship, and a financial strength index, for 2010: the most complete recent year available for these data (S1 Code; Fig S1). These data were also linked to the most recent available prefecture-specific poverty rates^29^.

Supercentenarians recorded in the GRG database and born in the USA were matched to the 1900 census counts for state and territory populations^30^, and linked to the National Center for Health Statistics estimates for the timing of complete birth and death certificate coverage in each US state and territory^31^. Both the number of supercentenarian births overall, and estimates of supercentenarians per capita, approximated by dividing supercentenarian number by state population size in the 1900 US census^30^, were averaged across the USA and represented as discontinuity time series relative to the onset of complete-area birth registration (S1 Code).

To capture the geographic distribution of French supercentenarians, all 175 supercentenarians in the GRG database who were either born or deceased in France were linked to the smallest discoverable region of birth using biographical searches^24^. In addition, de-identified records in the IDL were already linked to birth locations encoded by the Nomenclature for Territorial Units level 3 codes (NUTS-3), which divide France into 101 regions ^25^. These modern regions were linked manually to their corresponding Savoyard-era department to obtain historic region-specific estimates of life expectancy at birth (denoted as e0)^32^ for the birth year and location of all supercentenarians in metropolitan France. For each supercentenarian, life expectancy at birth e0 was then measured relative to the contemporary average e0 of metropolitan France (S1 Code).

The number of total supercentenarians and SSCs born into Eurostat NUTS-3 coded regions, either documented for French and English regions in the IDL or estimated for Italian regional cohorts by ISTAT, were linked to modern socioeconomic indicators available at this administrative level: total regional gross domestic product (GDP), GDP per capita, GDP per capita adjusted for purchase power scores (PPS), murder and employment rates per capita, and the number of 90+ year-olds, using the EUROSTAT regional database^33^ (S1 Code).

Across England, additional data were obtained for the Index of Multiple Deprivation or IMD: a national metric used to indicate relative levels of deprivation, including income deprivation in people aged 60+, by the UK Office of National Statistics^34^. The IMD data are measured in 317 local authority districts, each of which is a subset of a single Eurostat NUTS-3 encoded region. To capture the relative degree of deprivation within England, the IMD and its component scores were averaged within each of the 175 NUTS-3 regions (S1 Code).

Similar estimates of deprivation were obtained for French NUTS-3 regions, by downloading the regional poverty rates and poverty rates in the oldest available age group, ages 75 and over from the French National Institute of Statistics and Economic Studies INSEE^35^.

To overcome the three orders of magnitude differences in population size across subnational geographic units, the number of centenarians, SSCs and supercentenarians were adjusted to per capita rates. However, the ‘correct’ adjustment for per capita rates of remarkable longevity is dependent on the *a priori* assumptions of their cause. For example, if the null hypothesis was that all supercentenarians are ‘real’, adjustment for birth cohort size 110+ years previously would be a more correct method for best predicting the population density of supercentenarians. However, if the null hypothesis is that supercentenarians are more frequently modern-era pension frauds or clerical mistakes, per capita correction for a birth cohort 110 years in the past is of uncertain value for predicting modern events. In this latter case, the occurrence of supercentenarians would be better and more accurately predicted by correcting for modern population sizes. This concern is mitigated by the non-migration of most SSCs and supercentenarians with age – where data is available, two thirds died in their region of birth – and the spatial stability of poverty patterns over time^36^. That is, most supercentenarians never moved, and national poverty and income inequality maps barely changed.

Historical rates were used whenever possible, favoring the ‘real supercentenarians’ assumption. Per capita rates of remarkable age attainment, calculated relative to the size of historical birth cohorts, were downloaded from the respective government statistical bureaus of Japan and Italy ^26,27^. Due to the absence of birth certificates, USA supercentenarian data from the GRG were corrected to historical per capita rates based on population data in the 1900 US census^30^.

However, individuals in France and England were located into geographic units that have only existed since 2003. As a result, there were no data on historical population sizes available for these geographic units. It was therefore necessary to estimate per capita rates using modern population sizes surveyed at the NUTS-3 geographic level within France and England.

To address this unavoidable difference in per capita rate calculations the number of the centenarians, SSCs, and supercentenarians were also corrected relative to the number of old-age residents in each modern geographic unit of Japan, England, and France (S1 Code). This adjustment was less susceptible to migration or large longitudinal shifts in population size, and better reflected the density of older people in modern geographic units after survival and migration processes. However, the insufficient granularity of birth cohorts within England and the considerable rearrangement of geographic units within France remains an important constraint on the upper accuracy of these models.

Collective socioeconomic indicators obtained for each country were used to develop linear mixed models across all regions for centenarian in Japan, SSCs in Italy and England and supercentenarians in England and France (S1 Code), to predict the regional per capita and per 90+ year old density of the oldest available populations in each country. Linear mixed models were fit using either the population poverty rate (England, France, and Japan) or estimates of old-age poverty rates (percent in poverty over 75 in France, the IDOP index in England) as the single predictor variable, and the number of centenarians, SSCs and supercentenarians both per capita and per 90+ year old. These models were then extended by fitting, as interactive effects, basic socioeconomic indicators used as global indicators of health and deprivation available at a sufficient geographic level (S2 Code). Such models focused on capturing fundamental indicators, representing crime rates, health, and income, available at the NUTS-3 regional level in the EU and the prefectural level in Japan.

Where available, French supercentenarians were linked to regional estimates of e0, calculated quinquennially for each of the Savoyard-era departmental boundaries of France into which they were born^32^. These local rates were then corrected relative to the contemporary French national average e0 to yield the relative life expectancy at birth, in years^32^. For example, Jeanne Calment was born in the Alpes-Maritime department in 1875, when average e0 was just 33.4 years and the contemporary French national average e0 was 37.8 years: a relative regional e0 of −4.4 years below the national average. These rates were used to measure if regional e0 was significantly higher or lower than the French national average, using a one sample t-test, for each supercentenarian.

To explore the potential for age manufacture amongst remarkable age records, birthdate data were aggregated within the GRG and IDL databases. Enrichment for specific birth days is usually indicative of nonrandom age selection due to fraud, error, and clerical uncertainty. This check, however, is limited in that it cannot detect diverse sources of error, such as identity fraud or failed death registrations, which retain a representative distribution of birth days.

As population representative birthdates were unavailable within the target populations, the distribution of births was tabulated by days of the month to remove the often poorly-categorized or undocumented effects of birth seasonality. This distribution was compared to both modern birthdate distributions from seventy million births in the US, which suffer distortion from the heaping of elective births and caesarean sections on certain dates, and to the distribution of birthdays under a uniform distribution of births.

To facilitate reproduction of these findings, all shareable data and code are available in a single structured file, with instructions and links for the non-shareable data, in S1 Data.

## Results

Between the 1880 and 1900 census, a period covering 79% of US supercentenarian births, the US population increased by 150% and average life expectancy at birth by 20%^30,37^. The introduction of complete vital registration in the USA coincided with this rapid increase in lifespan and population size and was expected to result in a large increase in the number of supercentenarian records per capita.

Instead, the introduction of state-wide birth certification coincides with a sharp reduction in the number of supercentenarians. In total, 82% of supercentenarian records from the USA predate state-wide birth certification. Forty-two states achieved complete birth certificate coverage during the survey period. When these states transition to state-wide birth registration, the number of supercentenarians falls by 80% per year (Fig 2a) and 69% per capita (Fig 2b).

**Figure 2.**
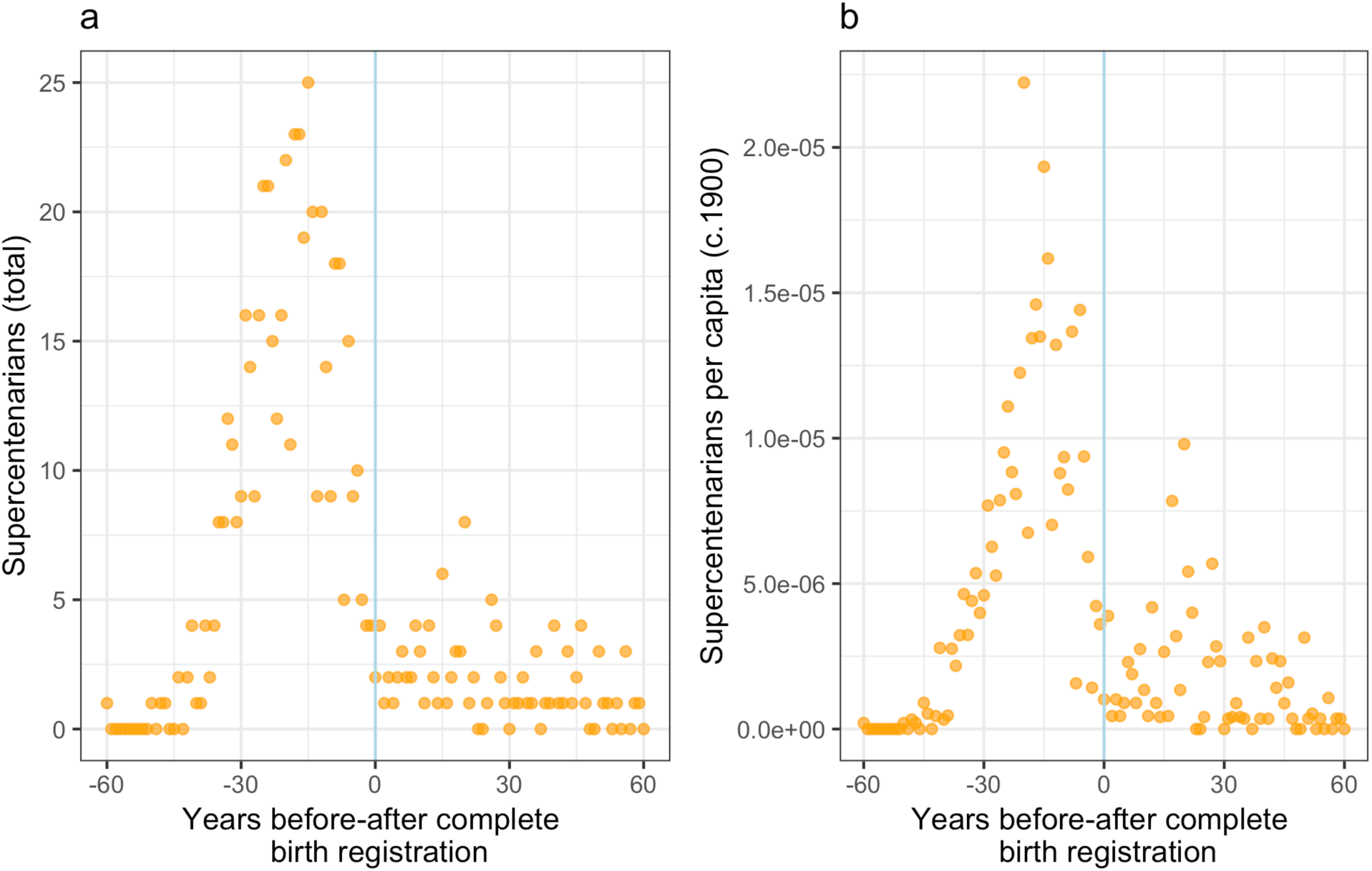
Number and per capita rate of attaining supercentenarian status across US states, relative to the introduction of complete-area birth registration. Despite the combined effects of rapid population growth and increasing life expectancy at all ages during this period, the total number of US supercentenarians (a) falls dramatically after birth certificates achieve state-wide coverage (vertical blue line). This trend remains after adjusting for total population size c.1900 (b) within each state.

The introduction of birth certificates in Italy largely predates the onset of supercentenarian records. Instead, the attainment of remarkable age in Italy is predicted by a short average lifespan. Within Italian cohorts, life expectancy at age 55 is positively correlated with life expectancy at all ages at all ages from 60 to 95. However, better early- and mid-life survival is predictive of worse mortality rates after age 95 (r = 0.15; p = 0.1; Fig S2a). Even within longitudinal cohorts, survival to age 55 – selected as a marker for mid-life survival – is increasingly negatively correlated with survival to ages 100 (r = −0.17; p = 0.06; Fig S2b), 105 (r = −0.38; p = 0.00003; Fig S2c) and 110 years (r = −0.45; p < 0.000001; Fig S2d), and with life expectancy at age 100 (r = −0.4; p=0.00001). That is, the better the mid-life survival of an Italian cohort, the worse their late-life survival: not cross-sectionally, but longitudinally within the same cohort. Furthermore, individuals across Italy are more likely to attain supercentenarian ages if their province has higher unemployment rates (Fig S3a), fewer people above age 90 (Fig S3b), and a worse economy (Fig S3c).

Findings from the Italian data support the hypothesis that centenarians, SSCs, and supercentenarians largely constitute a collection of errors and pension fraud^19^. For example, of Italy’s 114 provinces Medio Campidano ranked second for supercentenarians, third for SSCs and fourth for centenarians per capita. However, the province also had the second-lowest employment rate of all Italian provinces and the single lowest GDP per capita (both unadjusted and PPS-adjusted; Table S1). Individuals born in Medio Campidano were the second-least likely to survive from birth to age 55, after the Blue Zone province of Ogliastra. By this metric, the province of Olbia-Tempio was the seventh-worst province for survival to age 55, and according to Eurostat had the eighth-fewest individuals alive over the age of 90, yet somehow also ranked as the best province for survival to ages 100,105 and 110 (Table S1).

French supercentenarians are over-represented in the overseas departments, former colonial holdings, and Corsica (which is included in metropolitan France; Table S2): regions that historically constitute some of the most neglected, least well-documented, and shortest-lived administrative regions of France. At the first reliable estimate of population size in 1950 these departments contained around 1.7% of French citizens. However, at least 11% (N=16) of the French supercentenarians in the GRG database originate there: a 6.5-fold over-representation. This number increases when integrating the deidentified IDL data, which only includes regions monitored by Eurostat (Réunion, Guadeloupe, Martinique and Corsica), to establish the minimum numbers of supercentenarians born in each region. Once these data are included, at a minimum the overseas and former colonial regions of France contain twenty-four (15.5%) of a total 155 supercentenarians: Guadeloupe and Martinique each contain eight, French Algeria four, and one each in French Guiana, Saint Barthélemy, Réunion, and New Caledonia.

When IDL and GRG data are combined, the top six (NUTS-2 level) regions of France by supercentenarians per capita are, respectively, Saint Barthélemy, Martinique, Guadeloupe, Corsica, New Caledonia, and French Guiana. These six regions contain eight times as many supercentenarians per capita (1.3 per 100,000), and more supercentenarians overall (N=21), than the region of Île-de-France (0.16 per 100,000; N=19). This is despite Île-de-France receiving more than double the per capita income, being the longest-lived region in mainland France, and containing seven times as many citizens including huge numbers of (non-supercentenarian) Guadeloupe- and Martinique-born internal migrants^38^.

Within these regions, Martinique has the second-highest poverty rate both overall (29%) and for people aged 75 and over (31%) in all the 101 NUTS-3 coded provinces. Of the ranked regions only Réunion has higher poverty rates, with 39% below the poverty line. Amongst the unranked regions, French Guyana has a 44% unemployment rate, with 44% of households are listed as ‘poor’, and Guadeloupe a 24% unemployment rate^39^. The two departments of Corsica have the highest and second-murder rate, and the third and fifth highest poverty rates, of the 101 French departments.

When compared across the 101 modern NUTS-3 coded regions of France, Guadeloupe and Martinique are both equal second for total supercentenarians after Paris (Table S2), with at least eight supercentenarians each. However, these two provinces contained only 190,000 French citizens each c.1900: by comparison the Paris department contained 2.7 million people in 1901. The two small, remote departments of Guadeloupe and Martinique produced the same number of supercentenarians as the Nord department, despite the latter containing ten times as many citizens in 1901.

When measuring supercentenarians per capita using modern population sizes, due to incomplete data, Martinique and Guadeloupe ranked second and third for supercentenarians per capita out of the 101 NUTS-3 regions of France. Creuse ranked as having the highest percentage of oldest-old individuals in France. Creuse also ranks as having the fourth-highest poverty rate in older people, the 16^th^-worst poverty rate overall, and the fourth-lowest GDP per capita (both raw numbers and PPS adjusted; Table S2) of the 101 regions of France. Per capita rates in Creuse may have also been impacted by massive out-migration, with the Creuse population falling by 60% since 1901.

Region-specific estimates of e0 were complicated by large-scale historical changes in province number, boundaries, and even nations. From 1900 the number of French regions fell from 88 to eighteen, with just twelve mainland regions. Outside the overseas regions and Corsica, whose boundaries are largely unchanged, few of the GRG supercentenarians could be located within the Savoyard-era department boundaries required to gain an accurate estimate of e0 values. Most records could only be resolved to the 12 modern administrative regions, which contain up to thirteen Napoleonic/Savoyard era provinces each, resulting in a wide range of possible socioeconomic data and e0 values for each individual. Combined with the initially small number of supercentenarians in France, this coarse resolution excluded a more detailed analysis of the GRG data.

There were notable disparities in the abundance of the oldest-old across England (Table S3). For example, the region with the most 90+ year-olds per capita^34^, West Sussex, had only five SSCs in a population of 460,000. In contrast, in Tower Hamlets, the region with the least 90+ year olds in the country^34^, records fifteen SSCs amongst 289,000 residents.

As a result, Tower Hamlets has the most SSCs per capita in England. Tower Hamlets also has the highest poverty rate^40^, highest child poverty rate^40^, highest income inequality^40^, the shortest disability-free life expectancy by ten to 15 years^40^, and the worst index of multiple deprivation^34^ of the 32 London boroughs. According to the Income Deprivation Affecting Older People Index (or IDOP), which captures the financial stress and deprivation of older people relative to the national average, Tower Hamlets is the single most income-deprived population of older people across all 317 local authority districts^34^. Tower Hamlets has the smallest percentage of people aged 90 and over^34^: a notable discrepancy for a region that, according to the IDL figures, has the largest percentage of SSCs.

Of the 131 regions included in analysis, the top 20 regions for SSCs per capita contained the 1^st^ and 3^rd^-8^th^ most income-deprived regions for older people in England (Table S3). As such, high rankings for income deprivation in older people predicts higher rates of achieving ages 105 and over. Out of 175 regions in England, for example, Southwark and Lewisham ranked as the eighth most income-deprived district for older people^34^, second for SSCs per capita, and first for SSCs overall (Table S3). Central Manchester produced 18 SSCs overall (equal sixth) and ranks 14^th^ for SSCs per capita yet is the third most income-deprived district for older people, and has the highest crime index, third-worst population health index, fourth-worst index of multiple deprivation, and sixth smallest percentage of 90+ year old people of any region (Table S3). Such rankings are not recent, with inner-city Central Manchester the second most persistently deprived of all 317 local-authority districts^34^.These specific cases reflect broader patterns across England, where regions with the highest number and per capita rate of SSCs also have lower incomes, worse health, more deprivation, and higher crime rates (Fig 3; Table S4).

**Figure 3.**
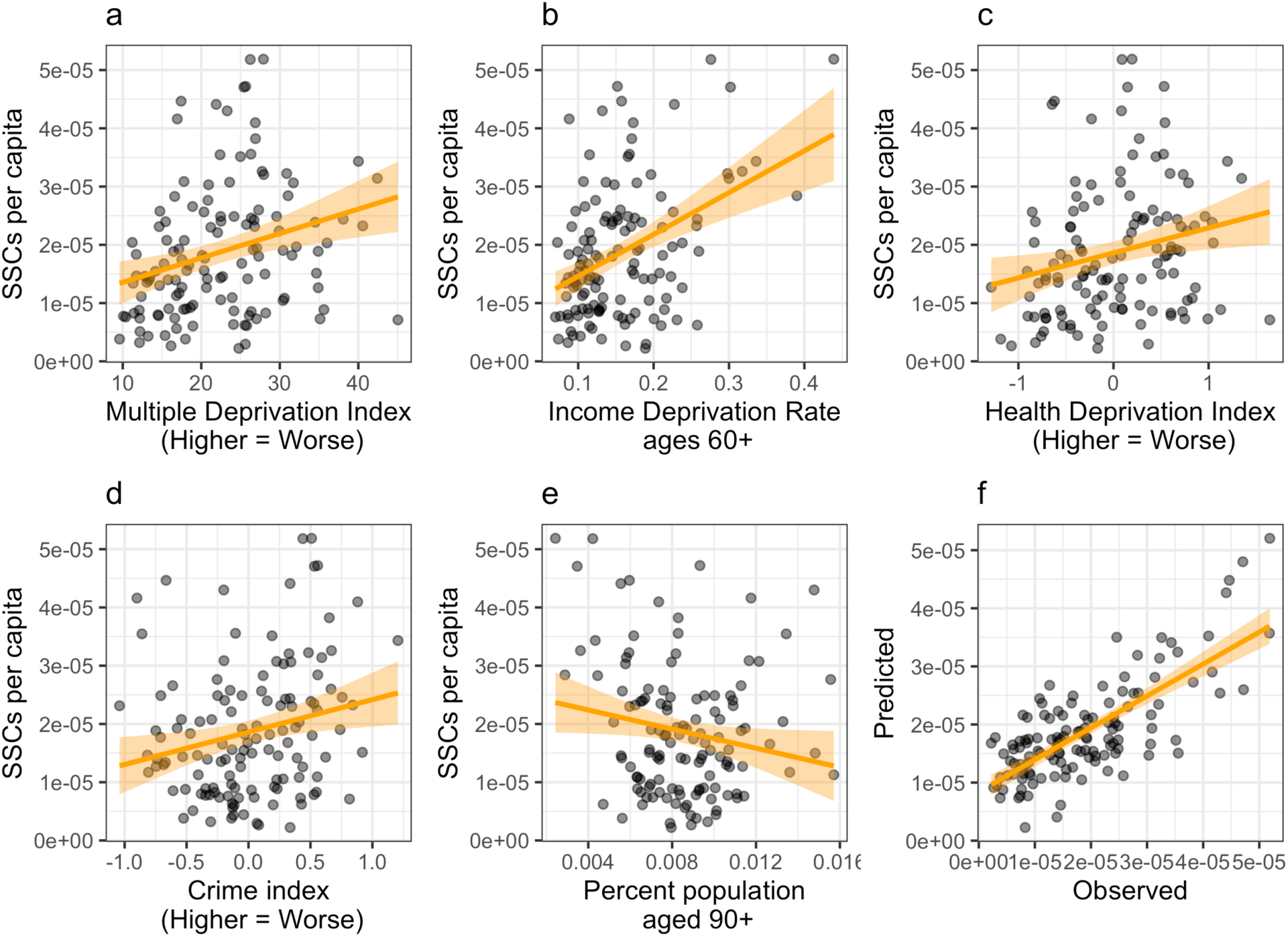
Social hardship and the distribution of the oldest-old in England. Across 128 regions containing at least one semisupercentenarian, the number of SSCs per capita is highest in regions where: (a) people are more deprived overall (r = 0.28; p = 0.001; 2019 data), (b) people over age 60 are more income-deprived (r = 0.42; p < 0.000001; 2019 data), (c) multidimensional health indices are worse (r = 0.23; p =0.01; 2019 data), and (d) crime rates are higher (r = 0.22; p= 0.01; 2019 data). In contrast, regions with a higher fraction of people surviving past 90 years of age (e), have significantly fewer SSCs per capita (r = −0.18, p = 0.04; 2015 data). Integrated into a simple linear mixed model (f), these five variables plus PPS-adjusted GDP and employment rates, collectively account for half of the regional differences in SSC abundance (most recent data available used; multiple R^2^ = 0.54; adjusted R^2^ = 0.39; p = 1e-06; Table S4).

Higher poverty is a particularly good predictor of higher survival to extreme ages. For example, a single-variable linear model using only (general or old-age) poverty as a predictor captures a substantial amount of the variation in centenarian, SSC, and supercentenarian abundance. Higher poverty and deprivation are positively correlated with all age categories of the oldest-old in every country measured (Fig 3; Fig 4a-f).

**Figure 4.**
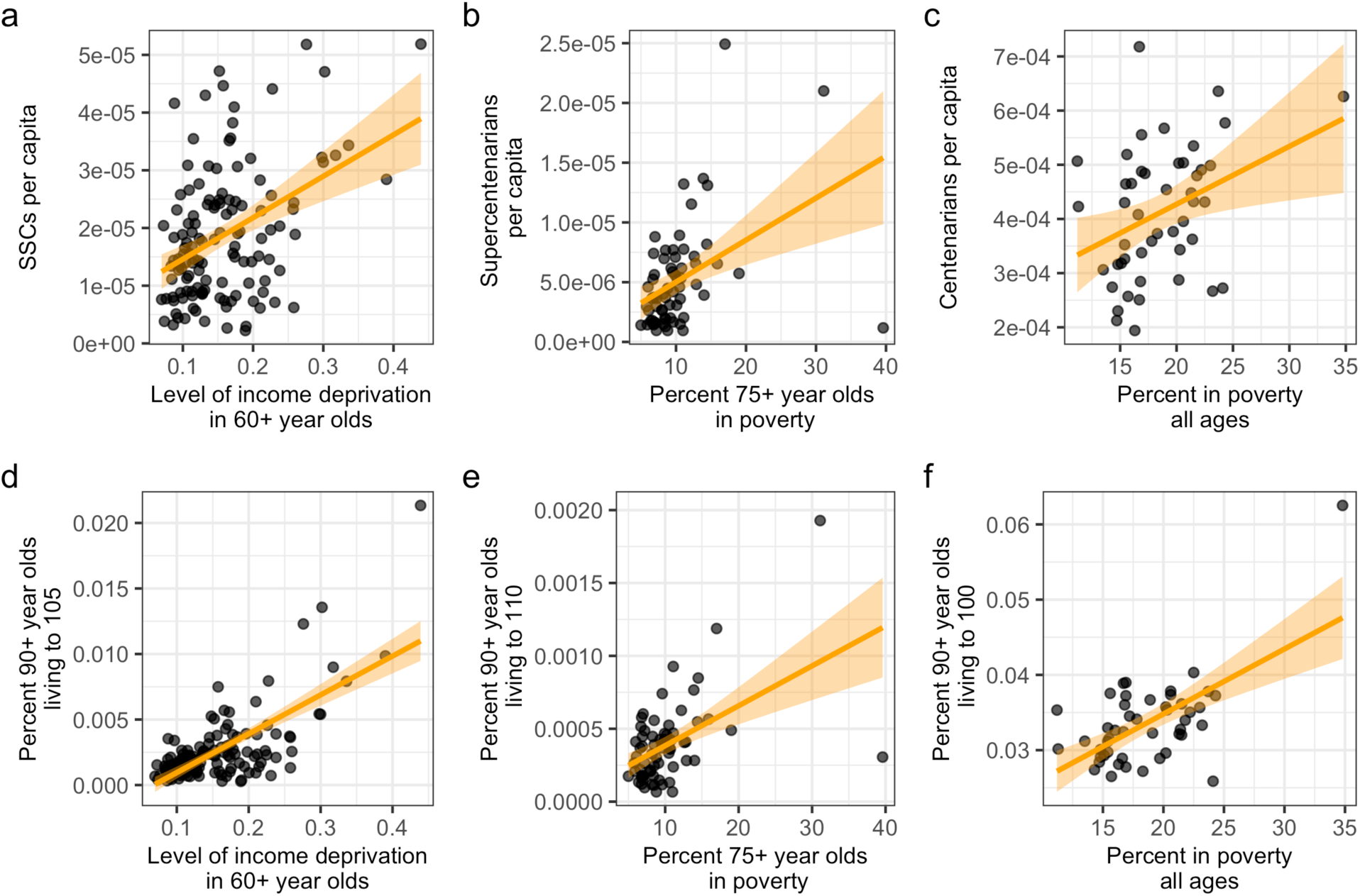
Old-age poverty and the density of the oldest-old. A higher percentage of people are centenarians, SSCs and supercentenarians in high-poverty and income-deprived regions of rich countries. Metrics of poverty and old-age poverty are positively correlated with the density of the oldest-old per capita across subnational data in England (a; r = 0.42; p < 0.000001; 127 regions; 2019 poverty data), France (b; r = 0.42; p = 0.0004; 66 departments; r = 0.58 in the 63 mainland departments; 2016 poverty data), and Japan (c; r = 0.36; p = 0.01; 47 prefectures; 2012 data). These relationships strengthen if density of the oldest-old is measured, not per capita, but by the fraction of 90+ year old people who are over age 105 (d, England; r = 0.71, p < 2e-16), age 110 (e, France; r = 0.51; p = 0.00001), or age 100 (f, Japan; r = 0.62, p = 0.000004; all data were the latest available national poverty statistics).

The IDOP ranking is an accurate indicator of the distribution of income deprivation in older people across England. When aggregated into NUTS-3 regions containing at least one SSC, IDOP scores are positively correlated with SSC abundance per capita (r =0.42; p = 0.0000009; Fig 4a). That is, higher income deprivation for residents over 60 predicts *higher* numbers of people surviving past age 105 (Fig 3b). This predictive accuracy improves markedly when predicting the number of SSCs per 90+ year old resident, rather than the number of SSCs per capita (r =0.70; p < 1×10^-15^; Fig 4d).

Like English data, the percentage of French supercentenarians in a population was predicted by higher rates of poverty in older people (r = 0.42; p= 0.0004; Fig 4b). Again, these associations were stronger when predicting the fraction of 90+ year-olds who lived over 110: across the 66 regions of France with sufficient data, the poverty rate over age 75 predicted half of the variation in supercentenarian abundance (r=0.51, p= 0.00001; Fig 4e).

Japan displayed similar anomalies in the regional distribution of extreme-age records. The first and second ranked Japanese prefectures for centenarians per capita, Shimane and Kochi, had the worst and second-worst regional economic rankings (Table S5), while extensive anomalies in third-ranked ‘Blue Zone’ of Okinawa are detailed in the supplementary materials (Supplementary Materials; Table S6). Japanese poverty rates in the general population were again positively correlated with attaining remarkable lifespans (r = 0.36; p = 0.01; Fig 4c), an interaction that strengthens when poverty rates are used to predict the fraction of 90+ year old people living past age 100 (r = 0.62; p = 4e-06; Fig 4f). The number of centenarians in Japan is also negatively correlated with income per capita (r = −0.44, p=0.001), the minimum wage (r = −0.64; p = 1e-09), and the Japanese financial strength index (r = −0.70; p = 3e-08) across all 47 Japanese prefectures. Prefectures that spend more money on old-age welfare per capita, a disincentive for welfare fraud, also produce fewer centenarians per capita (r = −0.49; p= 0.0004). These factors share latent drivers and are highly autocorrelated: prediction models for centenarians per capita based solely on poverty (R^2^ = 0.37; p= 0.0001; S2 Code) approach the accuracy of linear mixed models containing all available socioeconomic variables (R^2^ = 0.43; p = 3e-06) yet have a lower Akaike’s information Criteria (S2 Code).

Linking French historical^32^ e0 to some 143 French supercentenarians with sufficient data revealed that the cohort e0 values of each supercentenarian was not significantly different to the contemporary national average (Fig S4). That is, mainland French supercentenarians were not, on average, born into regions with either significantly longer or shorter life expectancies. Given it is only known where these individuals where born and died, but not necessarily where they lived most of their lives, it seems notable that the modern economic conditions and poverty rate of a birth location should be predictive of becoming the ‘oldest-old’ (Fig 4) while life expectancy at birth is not (Fig S4).

There is some ongoing debate as to whether urbanity is linked to higher historical mortality rates^41^, yet observed regional differences were not explained by any clear urban-rural divides. High rates of centenarian, SSC, and supercentenarian attainment was mixed across both rural (e.g. southern Italy, the overseas departments of France) and urban regions (*e.g.* Tower Hamlets, Manchester), with the primary determinant being poverty rather than urbanity. In addition, both rich rural (*e.g.* Hampshire, Savoie) and rich urban regions (*e.g.* Hauts-de-Seine, Kensington) routinely produce no supercentenarians and the extreme old.

Enrichment for specific days of the month in birth dates is seen as indicative of nonrandom age selection due to fraud, error, and clerical uncertainty. This check, however, is limited in that it cannot detect diverse sources of error, such as identity fraud or failed death registrations, which retain a representative distribution of birth dates.

Days of the month (e.g. the 1^st^ or 2^nd^ of each month) are nearly uniformly distributed throughout the year. As a result, US births had a near-uniform distribution of births, with minimal deviation from random sampling. Even after the widespread uptake of induced births, which avoid weekends and public holidays, birthdays generally varying by less than two per cent across different days of the month (Figure S5a), such that the distribution of births across days of the month is nearly invariant in the modern USA. In contrast, supercentenarians in the GRG database are 1.4-fold more likely to be born on the first day of the month (Figure S5b) and 1.2-fold as likely to be born on a day that is divisible by five. The number of supercentenarians born on the first day of the month is 150% higher than the previous day. Given the near-complete absence of caesarean sections in these population, this pattern may be explained if a large percentage of people have non-randomly chosen or manufactured birthdates.

These patterns are broadly reflected in the IDL birthdates, with days divisible by five, excepting the 25^th^, being over-represented amongst supercentenarians and SSCs (Fig S5b). In contrast the first day of the month was not over-represented amongst the IDL data (Fig S5c): this was initially difficult to reconcile, given these databases overlap considerably and largely represent the same individuals.

However, this differential appears to be a result of dates of birth being missing from 48% (N=797) of the IDL supercentenarians: despite extensive records being available, every US supercentenarian in the IDL has their day and month of birth removed. In addition, all Japanese records are absent. In the GRG data, most of the signal for over-enrichment arises in Japanese and US birthdates (Code S1-S2), with Japanese supercentenarians 2.77 times more likely and US supercentenarians 1.57 times more likely to be born on the first day of the month. In the primarily European countries in the remaining GRG database, individuals instead have a 0.67 odds ratio of being born on the first day: oddly, the inverse of the US and Japanese pattern. As a result, the omission of certain dates and records from the IDL data, for reasons that are unclear, masks the age heaping apparent in the USA and Japan.

## Discussion

Basic economic and social indicators in the modern economy, such as GDP per capita and poverty rates, predict the distribution of extreme age records. Despite constraints on model construction and accuracy, such as unavoidable differences in per capita adjustments, these basic models approached reasonable accuracy. However, the direction of these interactions is the opposite of rational expectations.

Diverse social and economic indicators that normally predict worse health outcomes, such as income deprivation^42,43^, poverty, and high unemployment, are all positively associated with a higher probability of reaching an extreme age. These factors are linked to a lower probability of survival and worse health outcomes at every age below 90, for every population included in this study^42–45.46^ For example in the USA, individual differences in income between the top and bottom 1% of individuals predict a substantial 14-year difference in e0 values: comparable to the 15 year e0 gap between, for example, the longest-lived country in 2024 (Hong Kong) and mid-conflict Yemen. The richest and poorest states likewise have a 8-year gap in life expectancy. In the UK, the primary factors predicting lower survival – across all ages below 90 – are poverty, unemployment, obesity, smoking, drinking, and disability rates^46^. These patterns monotonically increase with, for example, increasing deprivation leading to lower survival at all ages^46^ below 85-90.

However, these ‘anti-health’ factors exhibit a consistent positive association with extreme longevity. In England, which contains the only national data with sufficiently granular regional health measures, even poor health itself is positively associated with attaining a remarkable lifespan (Fig 3c). Across England and Italy, a larger number of people over the age of 90 is a significantly predictor of *fewer* people over age 105 (Fig 3e; Fig S3g-i): a difficult pattern to explain, given the younger population contains the older, unless errors are a primary factor.

Observed rates of vital registration in extreme-age and supercentenarian data are even more difficult to explain. For example, 96% of all 105+ year olds died since 1990, overwhelmingly in populations with over 95% rates of death certification. During validation, the intensive search for vital registration documents and the removal of unsupported cases should have enriched this death certification rates *above* the 95% baseline. It is therefore unclear why only 8% of all SSCs and 1.4% of all US supercentenarians recorded in the IDL have a death certificate unless these data are mostly errors. A similar pattern holds for birth certificates: even an intensive search could not exceed 15% coverage^47^.

Finally, claims to remarkable regional longevity are often contra-indicated by substantial, but curiously uncited, studies. For example, the Centres for Disease Control generated an independent estimate of average longevity across the USA: they found that Loma Linda, a Blue Zone supposedly characterised by a ‘remarkable’ average lifespan 10 years above the national average, instead has an unremarkable average lifespan^29^ (27^th^-75^th^ percentile; Fig S6). European Blue Zones also enjoy independent estimates. According to the collective governments of the European Union^48^, the Sardinian and Ikarian Blue Zones occupy regions that are 36-44^th^ and 56-65^th^ in the EU for old-age (85+) longevity. The Ikarian Blue Zone also falls into a shared statistical region with the much larger island of Samos, which it aggregated into in the census. Despite this, Ikaria and Samos combined only had thirteen centenarians in the 2011 census after the Greek government undertook at error-correction, and the observed rate of centenarian attainment was unremarkable.

Viewed in isolation these anomalies may, perhaps, be explained away by reference to unknown lifestyle factors. However, these findings should be considered in the context of other diverse and incongruous patterns observed in extreme old age studies. The global pattern of centenarian data, for example, reveals universal problems in the patterns of late-life survival^49^. These concerns are so extensive that discussion of even the broad outlines requires a critical review (appended as Critical Review).

Reflecting these wider problems this paper has now undergone and passed peer review, at BMC Public Health, by the majority of nine peer reviewers. These reviews have been underway since February 2024, were finalised in February 2025 with no substantial revisions required, yet somehow the paper is still not published. Contained in these nine reviews, and the extraordinary excuse-making of the editors who have failed to publish a paper that passed such an unnecessary threshold, is a curious case-study of the problematic nature of demographic science. These nine unsigned reviews, the editor’s reasons for non-publication, and my necessarily lengthy responses to all parties are appended in the Supplementary Materials as Ongoing Reviews.

The problems of extreme-age research may, however, be focused into two critical points. The first is observational. Extremely high-frequency errors are not impossible, improbable, or hypothetical^19^: instead, they have been routinely discovered after long evading the notice of demographers, often in the most intensively studied populations in the world. Any statement that such error rates could not happen^19^, for example due to the vigilance or rigour of demographic validation, are therefore belied by historical fact.

In the USA, for example, 27% (‘white male’) to 66% (‘non-white female’) of Americans had multiple official ages in 1960, with 8%-30% misreported by more than a decade^20^. At least 54% of US centenarians were revealed as errors in 1979^50^, and over half of all decedent African-Americans had multiple official ages in 1985^22^. In 2003, Stone checked 550 US supercentenarians recorded from 1980-1999 and, after an exhaustive search, just 43 of ‘supercentenarian’ records had a birth certificate (14.5%), and only 217 (40%) had a consistently-reported age ^47^.

Substantial error rates were recently uncovered in every ‘Blue Zone’. In 1997, thirty thousand Italian citizens were discovered to be claiming the pension whilst dead^51^. In 2008, 42% of Costa Rican 99+ year olds were revealed to have ‘mis-stated’ their age in the 2000 census^52^ and, after limited error-correction, the Nicoya Blue Zone shrunk by around 90%^53^ and old-age life expectancy plummeted from world-leading to ‘near the bottom of the pack’^54^. In 2010, over 230,000 Japanese centenarians were discovered to be missing, imaginary, clerical errors, or dead^55,56^ -- an error rate of 82% in data then considered among the best in the world^7,57^. Greece followed in 2012, when at least 72% of Greek centenarians reported in the census were discovered to be dead or, depending on your perspective, committing pension fraud^58^. Finally, at least 17% of centenarians in the USA were discovered to be non*-*centenarians in 2019, not through intensive validation or qualitative interviews, but by reading two plain-text files and finding the dates did not match^14^.

Only two of these cases were discovered, and all seem generally ignored, by professional demographers^14,52^. Some cases were seemingly not even noticed *post-hoc*: when most Greek centenarians were found to not exist in the aftermath of the global financial crisis, their non-existence was published in newspapers^58^, but was not mentioned or cited in the demographic literature.

If equivalent rates of fake data were discovered in any other field – for example, if 82% of people in the UK Biobank or 17% of galaxies detected by the Hubble telescope were revealed to be imaginary – a major scandal would ensue. In demography, however, such revelations seem to barely merit citation.

It seems worth asking, therefore, why the recurrent discovery of high rates of non-random age errors have been ignored by the scientific community, given the fundamental importance of accurate age data to fields like medicine, gerontology, the social sciences, and epidemiology.

A typical response to discovering most data is illusory has not been to stop trusting data built on identical methods and documents, but to instead call for better validation and a closer examination of documents^59^. This raises the second critical point. The examination of documents, the only measure of human age in virtually all demographic research, cannot detect routine errors.

Again, this is not an assertion but a reality. Consider, for example, the 1878 Taché investigation^60,61^ that pioneered testing centenarian status using validation ‘*proved by authentic documents, examined with a rigorous scrutiny’*: the unchanged state-of-the-art method. Of 421 claimed centenarians, 82 passed documentary validation, and only nine passed subsequent investigation, including Pierre Joubert: the world’s oldest man and first validated supercentenarian^60,61^. The case endured over a century of scrutiny, and was continually included in world records, before Joubert was found to have died at age 65. It was written on his death certificate the entire time^60,62^.

Extreme-age demographers seem to have misunderstood the central lesson of such events: not that errors can be detected though increased scrutiny, but that acceptance of such cases is dependent on the often-arbitrary generation, survival, and detection of documents. Without all three, completely fake cases can remain ‘validated’ indefinitely, regardless of whether they are true.

Demographers have confounded consistency with accuracy by assuming that consistent and valid documents are accurate, or at least strongly indicative of accuracy. This assumption is deeply unsound, especially given the nonlinear accumulation of error rates with age^15^ and the rate at which errors have previously passed validation. Showing that documents are consistent and valid does not mean that they are accurate, or that any connection to their supposed owner is real.

To understand this problem, consider a room containing 100 real Italian centenarians, each holding complete and validated documents of their age. One random centenarian is then exchanged for a younger sibling, who is handed their siblings’ real and validated birth documents.

How could an independent observer discriminate this type I substitution from the 99 other real cases, using only documents as evidence? More problematically, how could that observer measure the frequency of such errors, and discriminate whether one, ten, or a hundred ‘Italian Siblings’ are in the room, given they each hold validated and consistent documents?

This simple class of error cannot be excluded based on documentary evidence: every document in the room is consistent, real, and validated. Any such type-I error therefore has the potential to indefinitely escape detection. Scrutinising documents, usually the only method open to demographers, provides no guarantee of accuracy.

Compounding this issue, any younger relative is also likely to have sufficient biographic knowledge to pass an interview (which almost never occur) with an accuracy indistinguishable from a plausibly-forgetful centenarian. This is how at least three of the former ‘world’s oldest men’ remained long-undetected ‘Italian Sibling’ cases: Shigechimo Izumi and Jesus Arenas were their own little brothers, and Pierre Joubert was his own son for a century. In addition, almost no living or dead extreme-age cases have been interviewed, close to zero cases have been re-interviewed by independent scholars, most extreme-old cases are immune to follow-up interviews because they are dead, and even when conducted many interviews seem distorted by biases whereby sizable anomalies – like the forged or multiple birthdays in the Kimura case – can simply be ignored by the interviewer^63^.

The ‘Italian Sibling’ though experiment illustrates just one reason that type I age-coding errors cannot be ruled out — or even necessarily measured — using documentary evidence alone.

Instead, the experiment presents a substantial problem for remarkable-age databases, which can be embodied in a deliberately provocative, if seemingly absurd, hypothesis: Every ‘supercentenarian’ is an accidental or intentional identity thief, who owns real and validated 110+ year-old documents, and is passably good at deception.

This hypothesis cannot be invalidated by the further scrutiny of documents, or by models calibrated using document-informed ages^64,65^. Rather, invalidating this hypothesis requires a fundamental shift: it requires the measurement of biological ages from fundamental physical properties, such as amino acid chirality^66^ or isotopic decay ^67^. These metrics already exist and have been applied in a small number of cases but require improvement and broader application within human populations. Even application of these techniques to small random-sampled groups, or subsets at extreme ages, would constitute a serious step towards understanding undetected error patterns in human age documentation.

The long-hidden flaws of document-based age measurement may, however, be starting to unravel. New biometric methods of measuring ages^68,69^ are revealing huge discrepancies with document-reported ages. For example, measurements of age based on epigenetic data uniformly predict the oldest-old are 8-20 years younger than their paper records suggest^70–73^. Researchers have dismissed these estimates as the result of slower aging in centenarians, of mysterious aetiology, even when centenarians were measured to have undifferentiated or ‘non-significantly slower’^71^ aging rates. It is hoped another explanation — the epigenetic ages are accurate, and the documents are not — may eventually be considered.

Until then, we can only hope that the development of new and better biometrics of age^67,68,74^ may end our long reliance on documents, that we may finally stop ignoring the remarkable error rates observed in old-age demography.

## Statements and Declarations

### Ethics approval

This study was performed in line with the principles of the Declaration of Helsinki constituting a reanalysis of published anonymized data from publicly accessible databases. No ethical approval was required.

### Consent for Publication

Not Applicable.

### Clinical trial number

Not Applicable.

### Availability of data and materials

All code is supplied in the supplementary materials. All data is available on request from the author and, as appropriate, from the Gerontology Research Group and the International Database on Longevity. However, it should be noted that these groups did not employ version control at the time of the study and that the International Database on Longevity has, seemingly in response to the criticisms in this paper, deleted or hidden data on rates of case validation birth and death certification – without any explanation.

### Competing Interests

The author has no relevant financial or non-financial interests to disclose.

### Funding

The author declares that no funds, grants, or other support were received during the preparation of this manuscript.

### Author Contributions

SJN conceived, designed, analyzed, and wrote the study and supplementary critical review.

## Supporting information

Critical Review

Ongoing Reviews

## Acknowledgements

The author would like to acknowledge Dr Elena Racheva for advice and encouragement, Zoe Campbell and Prof Heather Booth for providing much-needed feedback and editing advice, Chris Mulligan for providing inspiration to use US birth data, Sally Morell and Dr Jim Docherty for interesting leads, Dr Gideon Meyerowitz-Katz for the final nudge toward publication, and an extraordinary cross-section of the scientific community for providing commentary and support.

## Notes

### Competing Interest Statement

The authors have declared no competing interest.

### Summary of Updates

I added almost 20,000 words of peer review, with extensive responses by myself, to nine peer reviewers and the editors. I call it the 'exhausted by a lack of answers' version. In addition, new material has been added to the critical review.

