## Supplementary material for "Supercentenarian and remarkable age records exhibit patterns indicative of clerical errors and pension fraud": Critical Review

### Supplementary Materials: Critical review

Issues raised in the analysis of extreme-age records suggest a remarkable situation: either that the anomalous pattern of extreme-age records can somehow be explained, or that extreme-age demography suffers extraordinary and systematic problems. Given the highly disruptive outcomes suggested by this analysis, the latter scenario requires some examination.

#### Indicators of systemic cultural problems in extreme-age demography

While individual experiences do not necessarily inform an entire research field, the author has independently uncovered multiple, recent, high-profile, and uncorrected issues that suggest deep-seated problems in the research culture of extreme-age demography. It is disturbing and noteworthy that that one individual can uncover broad-ranging problems, largely unaided, over such a short period.

First, a prominent Nature[1] paper on extreme ages rounded most of their old-age data down to zero[2] by accidentally using the wrong column in a life table ($l_x$ not $q_x$) and treating zero as equal to one in a log transform. The authors then concluding that the rate of change in this data, which they had accidentally rounded off to 0 they then turned into 1, was lower than the rate of change in unaffected data at younger ages[1]. Using the correct column of the same life table generates the opposite result[2] and contradicts the entire study. Despite these issues being revealed through open code and data, the paper remains unretracted and its many criticisms[2–7] ignored.

Barbi *et al.* then promoted an opposing view in Science[8] after picking the most empirically biased and worst-fit model possible from 861 alternatives[9,10]. Barbi *et al.* precipitated this remarkable coincidence by selecting "*middle age*" as the ages 65-80 in their regression[9]. Fitting exactly the same regression to any other age range eliminates the claimed plateau, contradicts every research finding in the paper, and provides a far better fit of their own data[9]. The author highlighted these problems through a peer-reviewed, reproducible, and open analysis[9,10] and confronted the lead authors who explained their choices in a way that indicated research fraud. Again, however, the paper remains unretracted and uncorrected.

Further published explanations of late-life mortality patterns were then uncovered that are similarly problematic – such as work by Alvarez *et al.* that simply hides old-age data, but not the error bars, above the y-axis (see their figure 2) before eyeballing the remainder and claiming the data "*levels off*"[11]. Other research merely stretches credulity, such as the idea promoted by leading demographers that inbreeding is somehow good for survival[12,13], but some striking and unchallenged explanations push outright scientific racism.

Recently, for example, Vallin[14] claimed that extreme age records on Guadeloupe and Martinique are a positive outcome of the African slave trade's "*selection of the strongest individuals*" through "*the tremendous health selection effect of slavery*", and that such hypothetical effects are further enriched "*due to low immigration and high fertility among black people compared to white*"[14]. Vallin then proposes, without providing sources, that gaps in supercentenarian status between these two islands and Réunion exist because mortality rates from slavery was markedly lower in the Indian Ocean: a claim directly contradicted by contemporary and historical mortality estimates[15,16].

Even ignoring the neo-eugenics, such explanations make no historical or biological sense. Over 5.3 million people, 8% of the mainland French population, are descendants of the transatlantic slave trade[17,18]. This population is over ten times larger than Guadeloupe and Martinique combined, and includes hundreds of thousands of Guadeloupe- and Martinique-born internal migrants[19]. However, despite enjoying better healthcare, higher wages, lower poverty, and more complete vital registration records of mainland France, this larger slave-descendant population has apparently not produced a single supercentenarian. Given 16% of birth certificates in supercentenarian families are missing (Vallin Table 6) it seems the claim that slavery enriches for 'robust' individuals is just racist window-dressing of poor-quality data[14].

This view is unfortunately not an anomaly, but a continuation of openly propagated racism in extreme-age demography. A general argument that late-life mortality crossovers[20,21] develop when weaker individuals are 'selected out' at a young age[22], an argument that seems unlikely given high early-life mortality predicts higher late-life mortality[20], has long been co-opted for racist ideas. In the words of Markides and Machalek "*While errors in the data are recognized as a possible explanatory factor, a consensus seems to have developed that the* [old-age mortality] *crossover is indeed real and results from higher early mortality that removes less* *hardy blacks*"[23]: a casually racist idea that has enjoyed decades of support, often because the primary alternative is acknowledging residual undetected errors in demographic data.

The director of the GRG Robert Young typifies this thinking, with a desire to resurrect scientific racism through his "biological superiority hypothesis" developed at Georgia State University[24]. Young proposes that supercentenarians are more likely to be African-American because the GRG director can (apparently) see that African-Americans have "*thicker skin than Caucasian-Americans*" which "*protects their internal organs*" because "'*Black don't crack*'"[24]. His other explanations include that "*greater selection pressures, and thus faster evolution*" caused supercentenarian status in African-Americans because of the "*pruning*" benefits of slavery (as with Vallin) and the forced breeding of slaves[24] because "*often the most-healthy African American males were selected to be the 'stud' that would then be mated with the healthy females; less-healthy slaves would be discouraged from breeding*"[24].

Young extends his racist theory to explain why the USA has supercentenarians and Africa does not He first states that racial 'pruning' must be moderate to be effective, as "*clearly, the Jewish population in Auschwitz had a very short life expectancy (akin to cutting a tree down, not pruning it)*"[24]. He then uses this argument as an explanation for the lack of African-resident supercentenarians[24] by equating living standards in modern Africa to being Jewish in Auschwitz.

It is hard to adequately capture the full scale of reprehensible ideas proposed by this key Supercentenarian researcher, who has been dramatically promoted since publication of these ideas, or by other individuals who use scientific racism as an explanation for longevity patterns. It is equally hard to explain why these ideas have been propagated for over 15 years, loudly and in public, without any comment or push-back from the demographic community.

Young is responsible for compiling one of the two major supercentenarian databases as director of the GRG, contributes to the IDL, and assembles many of the Guinness World Records for longevity using his qualitative assessment of supercentenarian data. That his qualitative assessment includes staring at African-American supercentenarians to see if they have 'thicker skin'[24] should ring alarm bells. In contrast, Young has been promoted to the top of the field *after* publishing such views.

Collectively, these issues suggest fundamental problems with the research culture of old-age demography. It is not that racist ideas and largely fake data are present in the literature, although these are extremely serious problems, but that such issues are met by a resounding absence of criticism or action. Excepting Dong *et al.*[1] none of the above dubious explanations, apparent misconduct, deeply suspicious models, or outright racism have been met with any perceptible response from the broader demographic community. Hundreds of thousands of downloads of this preprint, for example, have not led to a single public mention of the racism detailed here – not a social-media mention, let alone an outcry – and mute compliance seems to remain the default. That needs to change, rapidly, if we seek to understand the actual patterns of old-age human survival.

**Indicators of fraud and error in population-wide data and 'Blue Zones'**

Indications of a problematic research culture and a high error rate in extreme-age demography can be evidenced at all scales: broad population patterns, population studies, individual case evaluations, and even the basic logical evaluation of data and evidence.

Emblematic of these systematic issues are the 'Blue Zones'. Blue Zones (BZs) are five regions – in Okinawa, Costa Rica, California, Sardinia and Greek Ikaria – with claimed high concentrations of extreme age records[25–27]. The primary claim of BZ research is simple: that these regions have remarkable rates of achieving extreme old-age survival, or remarkable average survival in the case of Loma Linda and Costa Rica. In particular, it is routinely claimed these regions have unexpectedly high rates of reaching centenarian status[27].

The supposed enrichment for extreme-age survival in BZs is then subject to a host of secondary claims, each aimed at explaining the primary pattern of extraordinary longevity. Old-age survival in the BZs is supposed to result from diverse causes such as 'moderate' drinking at twice the NHS heavy-drinking guidelines[28], plant-based diets[25–27] and inbreeding[12,13]. Most prominent amongst these secondary claims is the idea that nine specific behaviours, the 'power nine'[29], underpin the achievement of extreme longevity (see accompanying manuscript Table S6).

The BZ longevity claims have been made, very publicly, for decades and have seen widespread support in the general demographic community. However, evaluation of the central BZ claims to extreme longevity, and secondary claims of their supposed cause, reveal the general neglect of error processes as a potential generative factor in remarkable age records, and more broadly reveal a lack of critical appraisal of extreme-age claims.

Consider the first BZ, the one most amenable to measurement: extreme longevity in Okinawa. In Okinawa the central claim of remarkable survival to extreme ages was established in 2004, before 82% of Japanese records were debunked[30,31]. Okinawa then had the highest number of centenarians per 90-99-year-old of any Japanese prefecture and was world-famous for remarkable longevity.

However, according to the statistics bureau of Japan, Okinawa also has the highest murder rate per capita, the worst over-65 dependency ratio, the second-lowest median income, the highest percentage (60%) of older people on welfare[32], and the highest unemployment rate of all 47 Japanese prefectures[33]. Despite prior claims of dietary benefits based on high vegetable[29], oily fish, and sweet potato consumption[26,27], Okinawa has the lowest per capita intake of sweet potato, fruits, vegetables, seafood, taro, shellfish, root vegetables, pickled vegetables, and oily fish such as sardines and yellowtail[34] of any prefecture.

Mortality rates in Okinawa 'cross over' after age 50, such that older individuals and cohorts have age-specific mortality rates far below the national average[35]: a pattern indicative of unreliable data and misreported ages[20,36]. Okinawa also has the second-highest per capita intake of beer and the highest per-capita intake of KFC, the highest child poverty rate at 36% (15% higher than the next-highest prefecture)[37], the most 'flophouses' and shotgun weddings[38], and according to USDA estimates Okinawan residents each consume an average 14 cans of SPAM per year[39]. Okinawa has the second-lowest minimum wage (by one yen)[33,38], the lowest household savings[33,38], the highest percentage of over-65s on income assistance[33,38], the highest poverty rate[37], and the worst average body mass index (BMI) of all 47 prefectures[38,40].

These rankings do not represent a recent sudden shift away from 'traditional lifestyles', and have not changed substantially for extended periods before, during, or after the BZ surveys[37]. For example, the body mass index of the over-75 residents of each prefecture has been continuously assessed since 1975, with the first measurement comparing residents born in 1900 or before to every other 75+ year old in Japan[40]. Every single year, over-75 Okinawans have retained the worst BMI [40]. Similar patterns hold across other nutritional indicators: in no case, at any point, have the massive population-representative surveys of Okinawa supported the claimed vegetarian dietary patterns pushed by BZ proponents[27,29,41].

These results become even more questionable given the capacity to directly assess each of the 'power nine' claims for BZ longevity[29] in Okinawa (accompanying manuscript Table S6). All nine claimed drivers of extreme longevity are assessable through data measured by the government of Japan. The 'power nine' claims are directly contradicted in every single case, usually through population-representative surveys of hundreds of thousands of people, with levels of inaccuracy that border on farce (accompanying manuscript Table S6).

If nothing else, these surveys reveal both astounding attention to detail by the Japanese government, and the scale of data that has been overlooked by BZ researchers. The older residents of Okinawa are not filled with purpose or "Ikigai" at remarkable rates: over-65 Okinawans have the 4th-highest suicide rate in Japan [42]. Older Okinawans do not "grow gardens"[29]: they self-report the lowest rate of gardening in the country, beating only the apartment-dominated Tokyo and Osaka megacities. Okinawans do not eat "Meat … only five times per month"[29] in 3-4oz. servings, which would total 5.1-6.8kg a year: they consume well over 40kg of meat a year[34,43] without including seafood. Nor do Okinawans overwhelmingly "belong to some faith-based community"[29]: they are 93.4% atheist[44], the most irreligious population in Japan, ranking third-last in the country for religious attendance[45].

What is most astounding about such claims is not that they have been used to generate profit by selling the fountain-of-youth repackaged for a modern wellness audience. Rather, it is that the demographic community, after hundreds of papers on the subject, does not seem to have read or cited the world-leading, comprehensive, population-representative surveys[37,40,44–47] that refute the BZ claims.

In Okinawa, the anomalous pattern of centenarians is even compounded by other error-generative processes. Japanese birth and marriage records are not generated by a central bureaucracy, but instead are hand-recorded by family members as 'Koseki' documents, which are then filed in local town halls and government offices. This combination of citizen self-reporting and government filing allows the propagation of errors without requiring fraud. On top of this self-report data, the large-scale bombing and invasion of Okinawa involved the destruction of entire cities and towns, obliterating around 90% of Koseki records[35] with almost universal losses outside of Miyako and the Yaeyama archipelago[48]. Post-war Okinawans subsequently requested replacement documents, described from memory[48], through a US-led military government that largely spoke no Japanese and which used a different calendar. As observed by Poulain[35] the number of replacement documents issued, a proxy measure of American bombing and shelling intensity, predicts 79% of the variation in centenarian status across Okinawa.

Despite the enrichment of regular error-generating processes through American firebombing, Okinawa is not an atypical case amongst the BZs. It is, rather, just the best-surveyed BZ region. Every proposed BZ displays patterns that suggest a dominant role of error, fraud, and (to phrase it generously) researcher degrees of freedom in explaining the distribution of extreme-age records.

As detailed in the manuscript, Sardinia, Ikaria, and Okinawa represent deprived regions of a rich high-welfare state. These BZ regions rank amongst the least educated, poorest, highest-crime and least healthy regions of their respective countries (accompanying manuscript Table S1-S2, S5). Therefore, it may be hypothesized that the relatively low incomes and high poverty rates of these populations generate age-reporting errors and pension fraud, and therefore remarkable age records.

The two remaining blue zones, Loma Linda and the Nicoya Peninsula, are considered exceptional due to their high average longevity rather than the presence of the oldest-old[25,26]. As such, these claims are not relevant to assessments of supercentenarian status, yet their analysis reinforces the evidence for broader problems in extreme longevity research.

Loma Linda is a Californian suburb containing just 23,000 people, designated as a BZ because of a supposed average lifespan of 86 years for females and 83 years for males. Even f taken at face value, this average lifespan is matched or exceeded by the 125 million citizens of Japan, the seven million citizens of Hong Kong, and the seven and a half million citizens of Singapore[49]. However, when assessed independently by the Centers for Disease Control (CDC) the five small-area survey tracts covering Loma Linda instead have an average life expectancy of 76 to 81 years[50]: the 27th to 75th percentiles of US life expectancy (Fig S6). This means, at best, the independent CDC estimates rank Loma Linda as the 16,101st most long-lived neighborhood in the USA (Fig. S6; S1 code). As such, it is again unclear why the lifespan of this community has been considered remarkable.

Like most BZs, Loma Linda is not a standard census tract or statistical area, but a custom-selected region within a larger geographic area: the largest US county of San Bernadino, which has an average lifespan of 78 years[50,51]. The Sardinian BZ was similarly delineated: by drawing circles on a map[12,25] and cut across the two standard, independently surveyed regions in Italy that have the lowest and sixth-lowest probability of survival to age 55[52]. The Nicoya peninsula in Costa Rica, where independent estimates are currently not available, is also a non-standard region cut from several independent census units of moderate life expectancy. Even worse, the goalposts have frequently moved on the BZ locations. For example, the Costa Rican BZ became two regions, and now one of the two Costa Rican BZs has now disappeared, and the other has shrunk by ~90% in population, while a third another has apparently appeared 300km away near Nicaragua[53].

Given the lower life expectancy of their encompassing regions and the uncertain basis of their ascertainment, it remains somewhat questionable that these custom-selected and frequently redefined regions should be considered valid outliers for human longevity.

**Indicators of error and fraud in health studies**

Like anomalous population-scale patterns, indicators of poverty and fraud and contra-indications of health are regularly ignored or downplayed in targeted studies of extreme age. For example, smoking rates of *e.g.* 17-50% and illiteracy rates of 50-80% are often observed in samples of the oldest-old[54,55]. Surveying the BZ of Ikaria, Chrysohoou *et al.* observed that the Ikarian oldest-old (ages 80+) have a below-median income in over 95-98% of cases, moderate to high alcohol consumption (5.1-8.0 L/ year), a 10% illiteracy rate, an average 7.4 years of education, and a 99% rate of smoking in men[56].

Seemingly more moderate rates of smoking occur in the Tokyo study of exceptional longevity, where "*few*" centenarians smoked, some 15.4%, of the mostly (78%) female centenarian sample were current smokers[57]. However, according to the national statistics bureau of Japan, only 3.9% of Japanese women and 19.3% of men over the age of 80 are smokers[58]. Tokyo centenarians therefore smoke at around twice the rate that could be expected in a younger, 80+ year old cohort with an identical sex ratio. Likewise, 80% of the 'exceptional' health-status centenarian population were daily drinkers, followed by 49% of the 'normal' and less than 40% of the 'frail or fragile' centenarians, resulting in "*a* [significant] *positive relationship between drinking habits and functional status*"[57]. In contrast with these figures, Japanese government surveys estimate only 2.8% of women and 23% of men aged 80+ drink every day[58]. Daily drinking peaks at 36.7% in men aged 60-69, the heaviest-drinking cohort in Japan[58]. As such, Tokyo centenarians drink at higher rates than any other age group, and smoke at rates equal to a 45-year younger population[58].

In the USA only 8.4% of general population over the age of 65 currently smoke[59], and in Europe 4.1% population over the age of 75 currently smoke[60] and 31% formerly smoked: figures that continue to fall with age due to two-fold higher mortality rates in smokers[61,62]. However, in the US and Europe individuals over the age of 100 often smoke and drink at higher rates than any other age group: in one US centenarian study 60% of people over age of 95 were former smokers[63], compared to just 25% of individuals over the age of 65 in the broader USA.

This latter study is notable as it compares lifestyle factors in the oldest-old to earlier surveys of the same cohort. This comparison revealed that centenarians have similar or worse body mass index, rates of physical activity, smoking and alcohol consumption than the younger baseline population from which they were drawn[63]: a striking equivalence given the baseline population was 35 years younger. Longitudinal follow-up studies of NHANES I cohort observed a >200% increase in mortality rates of smokers and overweight individuals during a 10-year follow-up[64]. As such, for these two samples of the same cohort to have a similar BMI, smoking, and drinking rates would require an implausible change in which smoking and obesity caused death until age 80-85, yet then somehow prevent death above that age to such an extraordinary degree that smoking and obesity rates increased again to match or exceeded their starting values[62]. [64]Alternately, it is possible that smoking, obesity, and drinking cause mortality at all ages, but that the decline in these factors with age is offset by rising vital statistics error rates with age[10] because smoking, drinking, and poverty-linked obesity are correlated with poor records and pension fraud. Suitable alternatives explanations may be that these centenarians had worse health, lower exercise and much higher rates of drinking and smoking than the general population at baseline, or that they constitute a population of younger individuals with lifestyle patterns and smoking rates that reflect higher rates of fraud, illiteracy, or chronic government neglect.

That is, harmful lifestyle factors like smoking tend to decrease in frequency with age because they kill people that engage in them. This happens until extreme ages are reached and harmful behaviors, anomalously, remain stable or even increase in frequency with age: as exemplified by the increased smoking rates in the oldest-old across Japan, Greece, and the USA. The only ways to explain such patterns is to either understand that errors increase nonlinearly with age and are correlated with smoking, or to imagine that smoking is protective from death at advanced ages.

While centenarian studies likely contain cleaner samples than supercentenarian populations, they often reveal broad assumptions on sample integrity. For example, the benchmark New England Centenarian study masks a widespread lack of birth certificates. Only 30% of the total ascertained sample had an official birth document discovered[65]. Of the 80 'centenarians' who were reported to be alive in 1995, roughly 8% were an entire century younger than reported, thirteen (16%) had reported a false birthday, and eleven refused to enroll but were investigated anyway[66]. Even in this post-screening sample, just 45 of the 67 investigated centenarians had a birth certificate. This rate of possessing documents is, inexplicably, substantially lower than the background population, which achieved complete coverage for birth certificates in all New England states[67] by 1897. For the remaining individuals who lacked birth certificates, age validation was instead carried out using documents including "*military certificates, an old passport, school report card, family bible, and baptismal or other church certificate*"[66].

The study is widely considered a gold-standard in aging research yet suffers both a troubling lack of documentary evidence, relative to the population baseline, and a reliance on qualitative judgement and "school report card" style evidence when birth certificates are absent. In one validation conducted by New England Centenarian Study staff and Robert Young, centenarians were ascertained by watching the internet for news stories and waiting for unsolicited emails[68]. Cases were then screened using the qualitative judgement of Young, who both promoted race science in his thesis on supercentenarians[69] and "*played key roles in* *subject recruitment, ascertainment of demographic data, and age validation*"[68]. The authors conclude that, in their sample of internet-and-phone-call ascertained cases filtered by Young, "*ascertainment bias is likely minimal*"[68]. Immediately after this screening "*age verification of centenarians began with obtaining subjects' birth certificates*" because "*stringent requirements for age verification were deemed necessary*"[68]. The lack of a birth certificates did not, however, cause centenarians to be excluded from the study: only 14 of the 35 prospective subjects, that "*had already gone through an age validity check*" by Young, had a birth certificate. Only two people who lacked birth certificates were then removed[68].

Missing birth certificates are also common in other samples of the oldest-old. For example, in the IDL database none of the 797 supercentenarians from the USA is listed as having a birth certificate[70] while 41% of US cases had the possibility of a birth certificate explicitly ruled out. Even an intensive search of 297 US supercentenarians by Stone[71] only managed, as an absolute best, 43 birth certificates (of which three were inaccurate, generating 13% coverage). This absence of birth registration data extends to the larger sample of supercentenarians and SSCs across the IDL data: only 19% of all supercentenarians, and 20% of the 'exhaustively' validated supercentenarians, have either an original or copy of a birth certificate[70] (Supplementary Code 1). Across all individuals in the IDL database, only 6.6% have an original birth certificate listed, and 74% of cases have no listed birth documents of any kind. These low rates of birth certification have not addressable in intensive follow-on searches: the most intensive search of vital registration data, conducted in US supercentenarians, discovered 43 birth certificates amongst 550 starting candidates[71].

Death certificates are also largely missing. The first follow-up survey of the population-representative NHANES I cohort, representative of the US population, found 3.8% of decedent men and 5.7% of decedent women did not have death certificates and remained alive on paper while actually dead[72]. However, for supercentenarians listed as dying in the USA under the IDL database, only seven are listed as having death certificates: some 98.7% do not have listed death certificates. In total only 15% of supercentenarians and 8% of SSCs in the total IDL database are listed as having death certificates[70].

These numbers are unachievable if these individuals came from a population with representative rates of birth and death certification, and suggest extraordinary – and likely 100% – rates of error or fraud. Exhaustive and intensive case validation should increase the fraction of the population with birth and death certification well above the background rate of certification – which in the latter case is at least a 95-99% rate of death certification. Instead, virtually none of these cases end up with a birth or death certificate after clearing validation. The only tenable explanation seems to be fraud or error, coupled with an extraordinary numerical blindness on the part of validating demographers. In response, it seems, rates of validation and birth and death certification are now hidden from the IDL database[73].

Similar failures of extreme-age health studies have a long history. In 1975 Professor Alexander Leaf proposed the existence of 'longevity zones' in Soviet Georgia, Pakistan, and Ecuador –regions where individuals apparently survived to remarkable ages[74]. Like the later Blue Zones studies, which also partnered with the National geographic society to garner press, this study proposed semi-vegetarian diets and mountain-village lifestyles as apparent secrets to longevity[74]. The study ignored or overlooked explanations that highlighted the poverty and bad record keeping and, like studies before and since, was quietly dropped from the longevity literature when it was discovered the project was based on bad data, age exaggeration, and poor records[75,76]. The longevity zone of Ecuador, for example, contained carefully examined documentary age data for 100+ year olds that was much later discovered to be fake in every instance [75,76] .

In the aftermath of these criticisms, prominent demographers Christina Chrysohoou, Christodoulos Stefanadis, Luis Rosero-Bixby, Bradley J. Willcox, Craig Willcox, and Gianni Pes have been quick to defend the existing BZs, with Gianni Pes in particular helping to platform[77] Robert Young without mentioning his systemic and directly relevant racism. This strident defense is somewhat hard to square with the admissions of the Blue Zone LLC founder Dan Buettner, who paid many of the above demographers to find BZs[78], that his discovery of Loma Linda BZ was not the product of careful scientific research. Buettner instead admitted he produced Loma Linda because of pressure "to find America's Blue Zone"[79] from his editor at National Geographic, and now implausibly claims that he then "never bothered to delist it"[80] as a BZ during 20 years of continuous and intensive lifestyle marketing campaigns.

This absence of factual grounding is disturbing, given the reach these ideas have in public health, public policy, and in the public imagination. These ideas are now being used to monetize the behavior of doctors through the "Blue Zone Certified Physician"[81] program, and capture public health funding through "Certification" sold by ShareCare Inc: a company co-founded by Dr Mehmet Oz, the three-time winner of the 'Pigasus' misinformation in science award. The BZ certificate subscriptions are so expensive that, despite receiving $25 million US dollars in co-funding, the US state of Iowa could not afford the program and opted out after three years[82,83] after which the BZ program was, ironically, replaced by a free equivalent. The basis of such expensive interventions includes one paper that: has no statistics or statistical plan, intervenes in public health without a legally-required ethics approval, and cites only four sources – two of which are cookbooks[84]. Despite this dubious background BZ concepts have risen to be seriously discussed as a guideline for policy interventions at the World Economic Forum[85]. That this forum should base their decisions on such an intellectually hollow concept is disturbing. No global public health policy should be based on a marketing pitch.

Unfortunately, no lessons seem to have been learned from these successes in marketing and failures in logic, and extreme-age health studies seemingly continue to ignore patterns where the claimed predictors of extreme longevity closely match the positive correlates of poverty, crime, and fraud.

**Indicators of error and fraud in individual cases**

The potential for bias during case validation is of marked importance, especially in the absence of such fundamental evidence as birth certification. However, individual case studies often highlight the potential for personal judgement during age validation. These problems extend beyond cases detailed in the manuscript, such as the century-long acceptance, and then rejection of the Taché investigation.

For example, the world's oldest man, Jiroemon Kimura, is now widely considered to be a valid supercentenarian case. Evidence for case validation used primary source material collected and vetted by biological relatives of Kimura, who stood to gain from his validation[86]. As such, Jiroemon Kimura has at least three wedding dates to the same wife, was conscripted to the same military three times in four years despite the mandatory conscription period being three years long[87], has three dates of graduation from the same school, and has at least three birthdays[86]. In addition, for the first 20 years of his life all of Kimura's birthdates and school records are actually those of 'Kinjiro Miyake', a name whose connection to Kimura is not attested to by an official document, but by an unofficial hand-written note from a Korean mail and telephone company[86]. Under interview, Kimura explained one of his extra birthdays in a way that was "*not feasible*"[86], and Gondo *et al.* decided the birth date had been deliberately forged[86].

Gondo *et al.* then concluded the case validation by assuming any conflicting official records were mistakes and, amongst the diverse birth, wedding, conscription, and graduation dates, selected those they felt were accurate. The paper describing the world's oldest man then lists a new birthday for Kimura as 1987, and his brother's birthday as 1985: two undetected and uncorrected errors that rather undermine the claimed capacity of demographers to spot erroneous dates[86]. The study concludes that "*no critical discordances were discovered*" [86] and the case is still considered valid[88,89].

The validity of the Kimura case has been widely accepted under the assumption that age discrepancies can be discarded through the qualitative judgement of demographers. This reliance on qualitative judgement during case validation reflects general practice. For example, concerns surrounding the validation of ages are often met with a response that biographical inconsistencies, detected during interview by a demographer, will result in cases being removed from the record.

However, this sentiment can be difficult to reconcile with observed practice. For example, former smoker and occasional drinker Adele Dunlap, who "*ate anything she wanted*" and "*never went out jogging or anything*" was validated by the GRG and IDL as the oldest woman in the USA. This was despite Dunlap consistently maintaining under interview that she was a decade younger: when "*asked how it felt to be 113, Dunlap… looked her questioner in the eye and answered*: *'I'm 104'*"[90]. Despite consistently maintaining her age was incorrect, for years, Dunlap remains validated as a supercentenarian on the basis of documents contradicted by her own testimony.

Further indications of the role of opinion, especially when ignoring contra-indications of health, are highlighted by other prominent cases. When interviewing the then-oldest man and woman in the world – Christian Mortensen and Jeanne Calment – demographers from the Max Planck Institute for Demographic Research issued the contradictory statement that Jeanne Calment smoked both one and two cigarettes a day for an entire century[91], followed by the justification that Calment "*possibly did not inhale at all*"[91]. It was then observed that, from age 20 to age 117, Christian Mortensen smoked "*mainly a pipe and later on cigars, but almost never cigarettes… he had also chewed tobacco…but never inhaled*"[91]. Why two people might choose to smoke for an accumulated 190 years, but never inhale, was not questioned.

Similar opinions aimed at explaining the questionable health habits of the oldest old are widely considered satisfactory. A notable fraction of supercentenarians smoke and drinking, yet these anomalous health patterns are routinely downplayed. Of the five oldest men ever recorded, Kimura and Mortensen are detailed above, Emiliano Del Toro (3rd) smoked for 76 years, Mathew Beard (4th) was busted for drink-driving at age 90, and Walter Breuning (5th) smoked cigars until he was 108. The oldest man in the UK stated his secret to health as "*cigarettes, whiskey and wild, wild women*"[92], while the former oldest man in the USA smoked 12-18 cigars and drank alcohol every day, which routinely started with "*a little bourbon in* [his] *coffee*"[93]. At least three of the ten oldest women drank every day, two smoked every day, and four are of unknown smoking status, while Jeanne Calment smoked daily, drank daily, and ate around a kilogram of chocolate a week.

These instances of poor lifestyle choices are not rare but constitute a substantial fraction of all supercentenarian cases. As summarized by Coles, supercentenarians lifestyle are characterized by "*heavy smoking, heavy drinking, or both, failure to exercise on a regular basis, and no conscious effort to eat nutritiously*"[94]. Instead of prompting skepticism, under the relatively safe assumption that smoking, drinking, poverty, lack of exercise, poor nutrition, and illiteracy should not enrich for remarkable longevity records, these contra-indications of survival are routinely ignored or downplayed. For example, the study by Chrysohoou *et al.* concluded that "*physical activity, dietary habits, smoking cessation, and midday naps*" predict extreme longevity in Ikaria: a conclusion that questionably re-shapes past smoking status as a positive indicator of survival[56].

Finally, a troubling aspect of case validation is not restricted to the questionable, and largely irreproducible qualitative choices made by extreme-age demographers. Nor is it the repeat capacity to overlook extraordinarily high rates of later-invalidated cases, or the overt racism of individuals responsible for case validation[24]. Instead, the rather simpler challenge is that cases are almost never subject to even cursory investigation, let alone intensive interview. For example, the IDL listed only 365 validations for 9,878 French SSCs (just 3.7%) of which none have an original birth certificate[70]. It is unclear how many of these 365 individuals were interviewed, even under the irreproducible and qualitative-driven framework of a field that does not encourage pre-registered findings or the publishing of non-results, and the IDL has now hidden or deleted these data as an apparent defense against any further investigation[73].

Interviews are virtually never reported or published, especially not in full, and subjects are seemingly never subjected to the double-blinded repeat interviews that would allow measurement of concordance in the qualitative assessment of extreme age cases. As such, the reproducibility of qualitative judgements are not measured across different contexts, populations, or interviewers. After more than two centuries of study we do not appear to know the test-retest accuracy of a blinded, independent investigator when validating an extreme longevity case. Instead, interview methods remain a rarely conducted, unregistered, non-blinded, routinely unpublished, and largely irreproducible process in which the demographer benefits personally from overlooking any errors.

More revealing than this oversight is the pattern of response by demographers to criticisms of extreme-age data. The criticisms above reveal how even careful validation, backed by interview, routinely overlooks errors at up to 100% frequency[95] and includes some of the most carefully and intensively validated cases in the world[96]. It is as if elite athletes were repeatedly passing a drug test called 'validation', when awarded an Olympic medal, only to be repeatedly discovered as cheats years or decades later. It is troubling, therefore, to see the primary response of many demographers has been to simply how carefully they 'validate' their cases, without acknowledging the past and ongoing failure of the validation method itself. The circularity of this reasoning is easily detailed but routinely repeated[97]. The author has repeatedly had to ask why careful validation repeatedly overlooks mistakes with a high or even 100% frequency– as a historical reality – to receive the illogical response that such mistakes cannot occur, because cases are carefully validated[97]. As noted above, the IDL has now deleted or hidden all data reporting which cases were 'exhaustively' validated and which individuals have birth or death certificates, without providing any answer as to why the rate of birth and death certification was routinely zero percent[73].

**Summary**

Individually these issues raised here may be, in some cases, answerable. Yet in aggregate such patterns reveal a devastating reality. The underlying data, methods, and research practices of extreme-age demography are so systemically flawed as to call the basic existence of 'supercentenarians' into question, and to raise serious doubts about the validity and integrity of the entire field. This shortfall of rigor is so complete in extreme-age demography that, not only are critical questions unanswered, they are unasked.

The list is extensive. Why have population and lifestyle surveys of Japan, which cover over 94% of households, not been used to evaluate population and lifestyle patterns in Japan after twenty years? Why are a Nature paper that rounded its data off to zero, and a Science paper using the most empirically biased model possible, unretracted? Why do demographers trust a single metric of age – documents – when documents have routinely generated observed error rates well above 50%, and exciting alternatives are being developed? How might documentary validation or interview methods ever *reproducibly* solve the 'Italian sibling' problem? Why are leading demographers allowed to state that African-American people live longer because they "*have evolved 10% more genes*"[24], or are more "*robust*" due to "*faster breeding*"[14], whilst being promoted to senior roles in major institutes, without a solitary word of criticism?

Instead of being asked, such questions are ignored, basic critical analysis is derided, and fundamental problems written in reproducible code have no effect. Such responses may be effective in removing criticism and have resulted in the author leaving the field. But the future of demography is saddening indeed if it cannot bring itself to answer these questions alone.

I may leave demography with a final question. Half of all births in India, for example, have no birth certificate: these people are excluded *de facto* from age-stratified medical and epidemiological models, because we don't know their age. How much better could our collective medical, demographic and epidemiological models perform, if we replaced document-measured ages with accurate biometric measurements of age across every population that, like India, suffers from systematically unreliable or absent documents[98,99]?

If ‘Italian sibling’ problems were taken seriously, and solved, we may find out. Given the reaction and conduct of the demographic community so far, however, hope for such an answer seems impossibly remote.
