## Supplementary material for "Supercentenarian and remarkable age records exhibit patterns indicative of clerical errors and pension fraud": Ongoing Reviews

### Initial Response to Review

**Submitted February 2025**

**Author's General comments to the reviewers**

I would like to thank all nine reviewers and the editor for the care and time they have taken in responding to the paper, and the overwhelmingly positive tone taken by so many people when assessing a disruptive idea. I have made my best attempt to answer the many good-faith reviews here, incorporate the surprisingly small number of suggested technical changes, and correcting any errors I have made. As modern science is often built on logic, analysis, and numbers, I am also relieved and gladdened to see that even the most adversarial of reviewers have found little to fault on a technical basis and largely disagree based on qualitative points of opinion or interpretation.

I apologize unreservedly for the time taken to respond, as I have been delayed by the unexpected number of reviewers, by personal matters, and by my own slow-and-steady thinking process which I cannot ever seem to forcibly hurry along. Unfortunately, given the remarkable number of reviews here, I will not always be able to accommodate all desires of all nine reviewers. This is especially the case when it comes to broad non-specific claims raised by individual reviewers. I shall, however, endeavor to do my best to make as many reviewers happy in as many cases as possible.

With warm regards,
Saul Newman

### Reviewer Reports and Replies

#### Reviewer 1

The empirical analyses in this paper are excellent. the authors did an incredible amount of work, and their analyses are very well-done.

However, there are two really major flaws with the paper:

1. It really does not do justice to the extensive and very elaborate literature on predictors of death. We know so much more about longevity than you acknowledge. To be clear, I am not saying that you can add a little text to references previous research. This is a major flaw in the paper: we already know the answers to the questions you ask!
2. Related to #1, your findings really add nothing to the literature. In fact, they simplify a broad, well-developed literature into a few simple points. I know that previous work has not studied extreme age much, but the predictors are the same....and we know so much more than you acknowledge.

I wish I had stronger feelings overall about this paper, but I just don't see how it contributes to the literature (beyond really nice figures).

**Author's Reply (SJN)**
Thank you very much. I am happy to see that nothing technical in the paper needs to be addressed, and I am happy to leave any question of relevance, novelty, or contributions to the literature to the editor and the members of the scientific community who have taken an interest in these findings. I have added a small and, admittedly, insufficient sample of the very many predictors of mortality at younger ages and regret that further discussion is outside the scope of an already lengthy paper. Thank you again for your review.

#### Reviewer 2

**(R2)**

This review will address what appear to be two different papers -- the formal submission and the supplementary note that is far more comprehensive. My general comment is that questions about the validity of claims of extreme ages being achieved in humans is legitimate -- especially given the history of embellishment, exaggeration, and lying that has permeated the field since the 20th century when work on people from Soviet Georgia revealed multiple problems with age validation or lack thereof. When the BZ concept was first proposed by the non-scientist that has since sold the brand (the term itself originated with a Belgian scientist), there was no plan at all to validate ages. This indicated that the motive for this work was to profit from selling products associated with these alleged claims of extreme longevity -- and indeed, that is what happened anyway, even after a Belgian demographer was brought on board to validate ages. In short, questioning the ages of these individuals is legitimate, but it must be done without bias -- not based on inference, as I'm detecting here. Some of the reverse bias appears in the supplement where equally dubious statements have been made -- see below. In short, this discussion should be based on empirical evidence, not bias, and I see problems on both sides.

Specific points

1. The statement that superager status in the U.S. is predicted by the absence of vital registration implies that all superagers do not have vital registration records; it implies that the records that are available are false or misleading; and it implies that superager status has not been properly validated. There are also multiple unsubstantiated declarative statements in the manuscript and supplement that would need to be addressed to consider these points more seriously.

**SJN:** This is indeed the case. Exact counts of birth and death certification does indeed reveal that ‘superagers’ over the age of 110 resemble extremely poor-quality data, as virtually none of the extreme age records in the USA have birth or death records. The numbers in this manuscript were taken from exact counts of the numbers of 110+ year olds from the largest database of supercentenarians available, the International Database on Longevity. This database contained information on what number of overall and ‘exhaustively validated’ cases have birth and death certificates and the answer is, for the USA: 7/504 have a death certificate and, as stated in the accompanying critical review, “the most intensive search of vital registration data, conducted in US supercentenarians, discovered 43 birth certificates amongst 550 starting candidates[1].” Many of these were not original documents, but later replacements. Consider, as a further example, the passage from the critical review: “cases are almost never subject to even cursory investigation, let alone intensive interview. For example, the IDL listed only 365 validations for 9,878 French SSCs, of which none have an original birth certificate[2]. It is unclear how many of these 365 individuals were interviewed, even under the irreproducible and qualitative-driven framework of a field that does not encourage pre-registered findings or the publishing of non-results." That is, across the entirety of France, not one SSC had an original birth certificate recorded in the IDL.

I have provided citations for the source of these data, and the data themselves, in the supplementary materials, and I have updated the manuscript to emphasize these surprising numbers. If any further consideration were needed, I have included the typical process for the 'gold standard' of validation for the New England Centenarian study staff[3], aided by the overtly pro-racism[4] Robert Young: who was heavily involved throughout validation of this study, and whose qualitative evaluation was used to ascertain and assess the cases.

Inclusion of these numbers in the paper is now rendered more important as, in what bears a striking resemblance to a display of academic cowardice, the response of the IDL to these critiques has been to remove the number of individuals with birth and death certificates from their database – by hiding or deleting the number of exhaustive validations and both the BCERTIF and DCERTIF columns – rather than redress or even discuss this problem[5]. You may, therefore, now see a difference between the original copies I have supplied, and the version now provided by the IDL.

As such, I am afraid super-ager status is routinely not properly validated, and indeed cannot be properly validated because our current methods of 'validation' are inadequate to detect consistent errors in paperwork. These sentiments have recently been validated in discussion with leading authors in the field, including Professor Horvath, who are unable to articulate how a residual difference between a paperwork age and an epigenetically estimated age may be reliably assigned to either a clerical error or super-aging status.

**R2:** 2. Just because places that have high alleged supercentenarian populations also have poverty and overall shorter lifespans, does not preclude the overrepresentation of supercentenarians -- both can happen at the same time.

**SJN:** I agree this *may* be true, but I have proposed error patterns as an explanation, and that explanation seems to fit. I do not see how I might rule out all other possible (and here unspecified) hypotheses. I am very curious to know any testable hypothesis as to how it could reasonably occur, as heterogeneity seems to fail for several reasons described below, but unless you tell me what your explanatory hypothesis is, I am afraid cannot test it. Given the size of the paper already, I also feel that any new hypotheses may also be better left to later discussions, of which I am sure there will be many.

**R2:** 3. Beginning on line 365. I understand why the conceptual linkage between poverty rates and supercentenarian numbers is being raised. After all, one would ordinarily expect fewer people to make it to extreme ages when poverty is high, but perhaps the linkages are not as clear cut as indicated. I read the discussion of this in the supplement, and I saw reference to selective survival, which is a real biological phenomenon without reference to race. For example, in the immediate aftermath of the 1917-1919 influenza pandemic, life expectancy rebounded rapidly because susceptible older populations were removed from the risk pool by a virus that differentially removed that segment of the population. The implication from the supplement is that these are all race related arguments -- they are not. Viruses evolve along the same biological principles.

**SJN:** I have not implied these arguments are all race related, merely pointed out the gateway to unchallenged racism that persists in the field. In addition, it is not clear to me how you would establish that selective late-life survival is indeed a 'real' phenomenon, if the basis of the evidence is all based on the incredibly flawed data presented here and elsewhere. It is now clear that centenarian data are globally flawed to the point of being unusable, as are any and all conclusions derived from such data: please see analysis to this effect in Newman 2024 here[6]. As such, I have no intention of clouding a long-overdue critique of racism in demography by making unsupported arguments about mortality crossovers.

**R2:** 4. I don't see how the life expectancy of the regions into which supercentenarians were allegedly born, is that relevant. The number of people that fall into this age range are extremely small to begin with -- even when you add them up across all humans. Looking for correlations to life expectancy seems superfluous as you're making the inference that these supercentenarians should not be present in populations with low life expectancies (which can and often are influenced by high early age mortality -- which has nothing at all to do with extreme survival except shortening the survival pool available).

**SJN:** I cannot really understand the rationale behind the reviewer's argument. It is not at all clear why life expectancy at birth should not form a relevant predictor of mortality in a healthy population. The point also ignores the very clear negative correlations with mid and late-life mortality rates presented in the paper, such as the negative correlation between late-life survival and mid-life (age 50) survival in Italy, which occur longitudinally within the same cohort, or the negative correlation between the number of 90+ year olds and 105 year olds in the UK. I have therefore tried to make these analyses more prominent, but I am going to have to simply ignore the strange idea raised that life expectancy should not predict health. Finally, I would expect that ‘shortening the survival pool available’ would be a substantial contribution to rates of attaining extreme old age.

**R2:** 5. The reported birth dates is interesting to be sure (divisible by 5), and this is indeed an anomaly and should not happen, and it tells us that some of the reported dates are from memory and not from records. This is a major concern and I agree with the author here.

I think the proper approach here is to reassess a large sample of the supercentenarian cases, or as many of them as possible, to revalidate their ages. Then link the invalid cases to your correlation analysis. This paper is devoted to looking for correlations that don't make sense where supercentenarians have arisen, but I see the real issue is whether we can believe the validation exercises conducted by those responsible for validating ages. If you can convince me that a substantial number of the alleged supercentarian cases are false or highly questionable (past or present), then suddenly the correlation exercise would make more sense. To focus only on the correlation exercise seems to me to be more a position of advocacy. You may very well be right that there could be gross exaggeration and embellishment going on here; after all, this profession is filled with both entrepreneurs and scientists that profit from keeping this narrative of exceptional longevity going. The best way to address this is by examining the actual cases rather than a series of correlation arguments that may seem on the surface to make sense, but which have their own questions.

**SJN:** I think the reviewer may unfortunately be missing the point made at the end of the paper. This seems to be my fault, and I have sought to address it in the text by clarifying in the paper: ‘validation’ is not a useful method, and further ‘validation’ as currently described by demographers cannot ensure the accuracy of these data. Evidence for this is plentiful, including the 100-year-long false positive of the ‘validated’ Pierre Joubert, the existence of over 230,000 dead Japanese centenarians with generally accurate and validation-amenable paperwork, and the still-validated but obviously flawed Kimura case. That said, I have added a couple more cases where high-profile validations have failed to detect errors, regardless of scrutiny, for decades, to reinforce the unreliability of current ‘validation’ methods. In short, no, there is no point in attempting to ‘revalidate their ages’ as the validation method does not work.

**R2:** A couple of points on the supplement that I found problematic, and which make me question the validity of this particular exercise. I see multiple instances of declarative statements without proper citation (e.g., "82% of Japanese records are debunked" -- citation needed). What is the reason why the demographic community has not read or cited the refutation of BZ claims? You haven't presented their side of the case with any degree of rigor.

**SJN:** I provide citations for this figure elsewhere throughout the paper and have added an additional reference here. As for the response of the demographic community, I cannot present their counter-arguments if they have made none. This was, tragically, the point of the statement: that 82% of Japanese centenarians indeed disappeared, as well as 72% of Greek centenarians, but that the demographic community took no notice and did no soul-searching whatsoever in the aftermath. These cases are almost completely uncited in the literature, besides my own citations and the solitary paper I cited on Japan that seems focused on arguments to the effect that the shockingly high error rates observed did not affect average lifespan very much: a straw-man, which steadfastly avoids the widespread use of these data in extreme-age research for other purposes. That is the very shocking intellectual omission I am highlighting. Of course, if you could point me to any papers that deal with these substantial issues in any meaningful way, I would gladly cite them.

**R2:** On the suggestion of replacing document-measured ages with "accurate" biometric measurements of age, all I can say is that there are none, and you provide no citation here. I know this because I've personally been involved in this research; I'm familiar with the validation exercises (the problem is variability in rate of biological aging -- that is, people senescing more rapidly or slowly will show chronological ages that are higher or lower than their actual chronological age, but the science is not there yet to generate valid chronological age -- which is all you want in this case). This statement alone in this document gives me pause about this paper.

**SJN:** I don't quite understand this objection, and I think there is just a misunderstanding here that I have tried to fix. I had made a call is for the active development of a more accurate metric, not for using a metric that already exists, and I cite two robust starting-points that already achieve a decent level of accuracy[7,8] including in other species. These cited examples are two successful, if noisy, instances where ages are measurable by physical means: amino acid racemization of teeth[7] and carbon dating of corneas[8]. I apologize for the confusion, and I have attempted to make this distinction clearer in the paper. In addition, it seems that, as in the last comment, the reviewer is claiming I have not provided citations, even though they are present throughout the text, and I regret that they have overlooked them.

Second, the variability of biological age is largely derived from comparing cross-sectional variation in aging clocks, of various types, to documents. If there is a discrepancy between the clocks and the documents, it can simply indicate that the documents are incorrect, not that the individual senesces more slowly. I have been in contact with Prof Horvath via a journalist, and he has not been able to clearly answer how, for example, he would discriminate whether a 10-year gap between an epigenetic clock age and a paperwork record would be caused by paperwork errors, or by differential aging rates. He seems to have no answer.

This is unsurprising given the background. Repeated ‘intensive validation’ has repeatedly and spectacularly failed to spot obvious errors [9–14]. I have given many historical and current examples of such failures in the text, and there is no reason to suspect the recurrent failure of paperwork validation[9–14] has magically ceased creating errors after the arrival of an epigenetic clock. As such, we cannot currently tell if differences between paperwork ages and epigenetic clocks are real or artefactual. This unfortunately suggests that much of the work on ‘superagers’ may be driven by artefacts introduced through bad paperwork, not ‘super-aging’, and that intensive efforts are needed to test this possibility. However, such a discussion is largely outside the scope of this work.

#### Reviewer 3

**(R3)**
This study addresses an important topic in longevity research, presenting a well-structured argument. However, there are several areas for improvement and expansion:

1. Terminology and Metrics:
   - Recommend using "e(0)" to denote life expectancy at age 0, rather than the broader term "life expectancy."
   - The graphs appear to show period variation in life expectancy. Consider estimating synthetic cohort life expectancy using successive census data, if possible.

**SJN:** Thank you. I have included the use of e0 to denote life expectancy at birth, where suitable, retaining only a couple of instances where the use of the full 'life expectancy at birth' helps to clarify the text without remembering the definition. This cleans up a fair section of the methods. I have always used cohort life expectancy where possible, such as the Italian data, and have reinforced and clarified where longitudinal data are used in the text.

**R3:** 2. Comparative Analysis:
   - While the critique of four "blue zones" seems thorough, the treatment of Loma Linda warrants reconsideration. This community, with its significant Adventist population, may indeed exhibit longevity factors linked to lifestyle. However, the small population size (approximately 25,000) may limit reliable e(0) estimates, especially when considering subgroups. Can adjustments for race, sex, religious practice, etc. be made?

**SJN:** Since these comments have gone to press, the founder of the Loma Linda Blue Zone – Dan Buettner – has openly admitted Loma Linda was discovered, not using scientific evidence, but because his editor pressured him to "find America's Blue Zone"[15]. He then implausibly claimed that, during almost 20 years of intensive marketing and product-pushing, he "never bothered to delist it"[16] as a Blue Zone. These comments expose the striking intellectual vacuum behind the Loma Linda Blue zone, a sentiment reinforced when seeking to find the origin of the claim that Loma Linda residents live "ten years longer"[17]. The basis of these claims is a single study, the 'Adventist Health Study I', that did not measure people in Loma Linda but surveyed residents of California in general[18]. The findings were run by a Seventh-Day Adventist organization in a Seventh-Day Adventist university, only applied to a subset of Californian Seventh-Day Adventists and (despite the obvious conflict of a religious organization finding that their own religion was better at living than everyone else) were not independently replicated in either the Adventist Health Study II follow-up, or any other studies I have ever seen.

As such, I do not feel the Loma Linda Blue Zone warrants any consideration as a scientific concept. It appears to be a marketing scheme, arrived at under editorial pressure on a suspiciously short timeline, and situated in a community on the turn-off to Las Vegas from Buettner's house in LA. That such an idea was ever seriously considered in public health, let alone at a high level of public health for almost two decades, is a deeply disturbing outcome. I feel deeply sorry for everyone convinced by these arguments, and I have included this new evidence in the manuscript.

**R3:** - Include a discussion on the historical relationship between wealth and longevity. Historically, the wealthy often had shorter lifespans due to urban living conditions, potentially harmful medical treatments, and rich diets. Suggest referencing:
 * Woods, R. (2003). Urban-rural mortality differentials: An unresolved debate. Population and Development Review, 29(1), 29-46.
 * Fogel, R. W. (2004). The Escape from Hunger and Premature Death, 1700-2100: Europe, America, and the Third World. Cambridge University Press.

**SJN:** It is a good point that urbanity may be an important predictor here, and I suspect it plays a role in (for example) China, where rural and urban areas even implemented different birth registration systems at wildly different rates until quite recently. I have added a paragraph discussing this idea, but the results are mixed. I am reluctant to wade too far into an already complex debate surrounding these historical patterns, but I have done my best, as I am afraid there is not sufficient space for a lengthier discussion.

**R3:** 3. Data Presentation:
 - All figures should clearly state the year of data collection.
 - Ensure figures are self-explanatory with complete references where applicable.

**SJN:** Figures should now be complete and figure legends now state the year of data collection.

**R3:** 4. Broader Context:
 - Expand the discussion on the blue zones paradigm's influence on policy making, global health prioritization, and lifestyle recommendations. Consider referencing recent developments, such as the Blue Zone Certified Clinician program. If the blue zones are false, this should be noted.
 - Address the feasibility of suggested improvements, particularly regarding widespread biomarker collection.
 - Include and reference other research that challenges the blue zones concept to

provide a balanced perspective.

**SJN:** These are very good suggestions, and despite the space constraints I have done my best to include them in the manuscript and critical review. I also find it extraordinary that, for example, these ideas have been seriously discussed at the World Economic forum and beyond, when they have so little basis in reality or fact. I am afraid that I have found no push-back to the Blue Zones concept that predates my own criticism, and the only other published challenges to the idea seem to come from rival nutritionists. I am reluctant to pass judgement on these ideas and include them, as the field is both outside my expertise, and often seems dominated by rivalries and commercial concerns more than scientific accuracy.

**R3:** 5. Future Directions:
 - Elaborate on the "best way forward" section, considering both short-term actionable steps to actually find good data on longevity by region.

Overall, while the stduy provides valuable insights, broadening the scope of analysis and discussion would strengthen its impact and relevance in the field of longevity research.

**SJN:** Thank you very much for your comments. I see very limited prospects for a ‘best way forward’ but I have included my most optimistic assessment.

#### Reviewer 4

**(R4)**
The idea of super longevity, that by living the right lifestyle in an accommodating environment, we can beat the odds and live well past 100, has clearly caught the imagination of demographers and not only that of the public at large. So much so, that the former's critical faculties may have been dulled by the lure of the so-called Blue Zones (places with concentrations of super-centenarians). The present paper makes a forceful argument for a critical reconsideration. Blue Zones, it argues, are a figment of statistical imagination, and poor statistics at that! Identified Blue Zones are scarce, with no apparent geographic logic to their location; they reflect unreliable or non-existing birth registration; and they are to be found in those places which are least likely to exhibit reduced mortality: poor, disadvantaged populations, with high crime rates and low probabilities of survival to age 90 and above. Consequently, it is concluded, zones of super-longevity are probably the result of faulty data or fraud, or both. The paper concludes by calling for a greater emphasis on biomarkers of old age and less on administrative records and individual memory, both of which are subject to error and manipulation.

I find the paper compelling, at least to the point of presenting a case that proponents of Blue Zones need to answer, but there are, I feel, a number of issues that need addressing.

**SJN:** I am extremely glad to hear this said, and I am, as I have been from the start, happy to address any issues.

**R4:** 1. The paper is mainly structured by location, covering the USA, Italy, France, UK and Japan, in each case discussing the reasons why the existence of the identified Blue Zones is unlikely. While this may be a good format for a provocative conference paper, in written form there is a sense of repetition combined with lack of focus on the main issue, namely that the Blue Zones, as identified, fly in the face of everything we know about good data and the conditions for survivorship and longevity. I would therefore recommend restructuring the paper in order to present the main counter-arguments to Blue Zones, namely the geographic distribution; data quality; and social conditions, using the identified Blue Zones as examples.

**SJN:** I would gladly consider restructuring the paper but, given the very large number of reviewers here, feel it may be unfair to return to them with an entirely new layout.

**R4**: 2. The logic of the paper suggests that Blue Zones are an unlikely phenomenon.

Unlikely, but not impossible. There are, I believe, certain arguments that do need to be discussed concerning the possibility of Blue Zones

**SJN:** While regions of better or worse extreme late-life survival will no doubt exist, their location is clouded by poor data at a substantial scale[6], I feel a more important first step is to discuss ways to overcome systematic age errors in human data. Otherwise, we run an enormous risk of repeating past mistakes with the same methods. Indeed, the discussions further down and in the manuscript note the incredible similarity of the current 'Blue Zones' to the previously-promoted-then-debunked 'longevity zones' proposed by Prof. Leaf of Harvard medical school in the 1970's. In addition, it has since emerged that at least one Blue Zone appears to have been invented under editorial pressure. I have elsewhere tried to measure extreme old-age survival using data untainted by commercial concerns (see Newman 2024 here[6]) but find that the question is so clouded by bad data as to be presently unsolvable.

**R4**: a. Many, if not all, of the identified Blue Zones, are in isolated mountain communities, even though the effect may be breaking down as younger generations take on more standard, 21st century lifestyles, including diet and locomotion. Nonetheless, might there not be something in these areas which might explain low mortality at very advanced ages?

**SJN:** I have added a (hopefully interesting) discussion on the history of this idea, especially by the inclusion of Professor N. Leaf's former work. Professor Leaf's work was taken very seriously and promoted the idea of long-lived, largely vegetarian mountain communities[19]. After error correction, however, it was discovered that these regions were long-lived because they were poorly documented and replete with fraud[13,14]. I would further suggest, although the idea is admittedly tentative, that remote mountainous regions may often have worse documentation because they are harder to reach, and therefore harder to accurately and routinely census, which may obscure the question of whether these regions have better historical longevity.

**R4:** b. The macro-determinants (or predictors) of mortality at very old ages may not necessarily be the same as those at younger ages. It is thus possible, even if improbable, that under certain circumstances a small group of nonagenarians survive to be centenarians, and semi- or supercentenarians.

**SJN:** This would indeed be improbable and – to abuse and rewrite a well-worn cliché – improbable theories require slightly remarkable evidence. I think any discussion of such evidence is beyond the scope of the paper and is likely best left to later discussions: I obviously cannot rule out, nor can anyone really rule out, the possibility that a different hypothesis may fit their data better. All I can do is highlight the explanatory power of error rates for explaining the observed distribution. I have added one brief discussion on a debate in rural-urban divides, where urban dwellers may have had lower historical survival. However, the debate is not settled and our observed patterns here cut across rural and urban areas in a way that does not fit this idea at all.

**R4:** 3. Some of the graphs (Figures 3 and 4) show a heavy concentration of rates and proportions at the low or high end of the distribution, thus allowing the regression lines to be led (or misled) by outliers. Transforming to logs or logits may symmetrise these distributions, giving a clearer picture of the relation between the variables.

**SJN:** Unfortunately, some patterns are somewhat visually 'squashed' by transformations. Log-transforming the data (which worked in e.g. figure S3) has not really eliminated this issue and has the unfortunate effect of distorting the shape of a linear regression in a way that often obscures the visual information of the linear model fit. In the very few cases where a data point approached or (for three data points in France, which changed the regression from $r = 0.509$ to $r = 0.501$) exceeded the threshold Cook's distance for significance, the regions to be removed were regions long proposed as exceptional. For example, in Japan the only data point with a Cook's distance of over 1 was Okinawa, which would hardly make sense to be removed completely as it is the focus of the investigation and widely promoted as the most well-supported data point: outlier tests are only justified for removing or identifying data errors. Removing it would therefore exclude many hypotheses that were being evaluated, and given the primary reason for removing outliers is the assumption that they are measurement or data errors, would leave me open to allegations of bias. For example, removing the French regions with poverty rates above 30% (see Fig 4b) would dramatically increase the strength of the positive correlation between poverty and supercentenarians per capita (from $r = 0.42$ to $r = 0.58$; respectively $p = 0.0004$ and $p = 3.3e\text{-}07$) and nominally strengthen my arguments. This would also, however, remove Martinique and La Réunion from the regression and the plot (two of the most interesting cases) on the dubious assumption that the poverty rates for these places were measurement or data errors. I believe these places really do have such poverty rates, and so removing these data seems difficult to support. I do not want to start throwing out the very exceptions we want to examine.

At the risk of some repetition, because this has come up informally, I have performed outlier tests for these regressions with precisely this concern in mind, to little or no effect. Primarily, the correlations become stronger in the 'wrong' direction – *e.g.* more poverty is even more strongly predictive of more extreme-age cases if you remove outliers.

In addition, log-transforming many of these variables obscures the absolute values. For example, it is an important qualitative as well as quantitative argument to see that poverty rates are so high, not just in relative terms but in absolute terms, for many of these regions. I think supplying the code and, due to personal threats from the GRG, privately the data is the best remedy.

**R4**: In sum, this is a valuable contribution to the Blue Zone literature, and even if unreviewed copies of it have been available for a number of years, it is important that it be published in the formal literature. I trust that the suggestions above will help strengthen the argument and thus give a better focus to the debate.

**SJN:** Thank you very much, and I hope that publication may finally generate some explanation or response, given the now thoroughly well-read preprint has been unanswered for so many years. I appreciate your help and apologize that I have been unclear in places.

#### Reviewer 5

**(R5)**
The issues addressed in the article are very important, but the main problem lies in the scientific nature of the argumentation. First, the number of centenarians with documented records is very small. According to the concept of percentiles, these people are among the top one in ten thousand. Establishing statistical connections between such a small proportion of the population and other meso-level variables requires great caution because these environmental variables are averages and may not necessarily apply to such a small proportion of people as centenarians. For example, how can it be proven that the relative poverty level of a region in a developed country will affect a person's basic survival needs? How can the suicide rate in other people's regions reflect the psychological state of centenarians in this group? How will the mean differences in smoking and drinking proportions between regions affect the centenarian group? Therefore, I think that using statistical regression methods to explain special groups does not necessarily have an inevitable scientific connection between the two, although this explanation seems reasonable.

**SJN:** I agree that establishing a causal connection here is not possible, and I doubt it would ever become possible. To fill this gap as much as I can, I have included as much detail as possible on the personal patterns of health in supercentenarians in the critical review and to a lesser degree in the manuscript, showing that these individuals do indeed have surprisingly high rates of (for example) smoking and drinking, and in addition have discussed direct measures of smoking in Japanese centenarians – who smoke at higher rates than prison populations.

**R5:** Second, when using environmental variables, the selection of these variables and the selection of their periods are relatively arbitrary, and no theoretical basis for selecting variables is given. For example, why is the survival rate of middle-aged people aged 55 selected to predict the proportion of centenarians?

**SJN:** This value was selected as it is the mid-life survival rate (halfway between 0 and 110) provided for these cohorts by the Italian statistical bureau ISTAT. That said, the result was largely indifferent to this choice of mid-life values as mortality rates are stable and highly concordant across all selected ages until roughly age 80. I have clarified this choice in the text.

**R5:** To what extent does birthplace information affect the current residence of centenarians? Is there any relationship between the selection period of relevant environmental variables and the current period of centenarians? We do not know key information such as whether centenarians have migrated and whether they have lived in the current area all their lives.

**SJN:** Environmental variables were selected to be the most recent available. Spatial patterns of crime and inequality display surprisingly stable long-term patterns: for example, it is unlikely that the poorest region of (say) the USA will suddenly enter the ranks of the most wealthy regions on a short timescale, or even over the course of many decades. Instead, environmental-social patterns are largely stable across long time periods, and I have therefore favored the most current (and therefore accurate) environmental variables available. I have clarified this rationale in the text. In addition, roughly 70% of supercentenarians were born and died in the same region across the sample. While this does not necessarily mean that they never moved – such data is unavailable – it does suggest that their residence patterns are stable long-term and that the use of environmental data is reasonable[20,21]. I have again sought to explain this possibility, and limitations of the environmental data, in the text.

#### Reviewer 6

**(R6)**
The author makes sweeping generalizations that almost all supercentenarians represent false cases. To support his views the author conducts multiple correlations of the number of supercentenarians with different socioeconomic indicators. The choice of these indicators is not justified and looks odd taking into account that most of these indicators are contemporary indicators reflecting conditions of much younger ages.

**SJN:** I disagree that these are sweeping generalizations: the cases here are backed up by a focused examination of many supposedly 'gold-standard' cases including the world's oldest man, for example, and it seems simply untenable to claim that the broad indicators of health chosen here (such as poverty) are somehow unusual or unjustifiable. That is, I disagree with the reviewer's suggestion that average life expectancy, income, poverty, and GDP constitute irregular choices for predictors of broad-scale health. These variables are the core predictors of health globally and are standard fodder for analyses of population health patterns. They are, in short, vanilla choices that are easily justified, and to suggest otherwise seems a bad-faith argument. I have justified why such choices were contemporary indicators: because pension fraud is contemporary. Furthermore, patterns of regional inequality remain stable across more than a century in the countries being analysed[20,21].

**R6:** I have the following comments about the paper:

1. It is not clear what is the rationale for selecting certain variables for correlations. Most of these correlations are weak. These correlations probably may need an adjustment for multiple comparisons.

**SJN:** I have reinforced the rationale for selecting predictors such as poverty, GDP, and life expectancy at earlier ages for use as predictors of survival at advanced ages throughout the text. These are, as stated in the text, arguably the most vanilla or unimaginative possible choices for predictors of population health. Furthermore, I have reinforced that it is not the strength of the correlations that is at issue, but their mere existence and their consistent inversion from what should be a perfectly inverted relationship, that is at stake. For example, it is not necessary to show that a negative correlation between the number of 90+ year olds and 105+ year olds is strong, merely that it exists, to call into question the quality of old-age data. Such a relationship should be impossible, much like the failure of supercentenarian researchers to detect this kind of anomaly, unless errors and fraud dominate the field.

**R6:** 2. Comparison with life expectancy at supercentenarian birth does not make any sense because this indicator strongly depends on infant mortality at time of centenarian birth.

**SJN:** If there is an indicator of adult mortality available for historical French departments, I will gladly use it as well: please provide me with a citation or link and I will include such data in my debunking. Absent such data, I will continue to focus on life expectancy as a sound and almost universally adopted indicator of health disparities, used because infant mortality and regional deprivation are so strongly positively correlated with adult mortality. This is why life expectancy at birth is used as a health indicator by virtually every government on earth. It is also a reason why supercentenarian data are strikingly at odds with other health patterns.

**R6:** 3. What is the reason to link the number of total supercentenarians to modern socioeconomic indicators if their survival was determined mostly by past conditions.

**SJN:** There are several simple reasons. Pension fraud is determined by modern conditions. Most exposure to mortality risk occurs late in life (even including the obvious exception of infant mortality, which is strongly correlated with later-life mortality risk up to about age 80), and is thus also determined by recent conditions. Furthermore, modern patterns of inequality strongly reflect past intergenerational patterns of inequality, which persists across decades or centuries within the countries analyzed[20,21]. This includes spatial inequality, which remains surprisingly stable over time across *e.g.* England and France for more than a century. I have reinforced this reasoning in the text.

**R6:** 4. It is not clear what is the accuracy of survival estimates to ages 100 and 110 years. Demographers often use models for these old ages. Before calculating correlations using data for small provinces it makes sense to study methods of life table construction.

**SJN:** These data are not determined by estimates or models, but by exact counts of 100 to 110+ year olds. The data sources are detailed in the supplementary materials and in the code. I am sorry I did not make this clearer and have added to the text to reinforce this. The only exception to the use of exact counts in cohort data are the cohort survival rates supplied by the referenced ISTAT database – the national statistical database of Italy.

**R6:** 5. Taking into account that survival to advanced ages is predominantly determined by familial longevity all correlations with contemporary indicators make little sense.

**SJN:** The analysis debunks the accuracy of such data and therefore raises serious doubts about the familial correlations derived from that data. As such, this comment appears to miss the point. It is illogical to disregard an analysis that shows survival to advanced ages is driven by bad data, by referring to findings derived from that nominally bad data. That is, if the analyses here are valid, the previous work detailing familial patterns of extreme longevity are invalid. As an aside, although it is hardly necessary, the evidence here also highlights a capacity for other simple explanations of extreme familial longevity. Crime, for example, runs in families too.

**R6:** 6. The author wrote: "Across England and Italy, a larger number of people over the age of 90 is a significantly predictor of fewer people over age 105 (Fig 3e; Fig S3g-i): a difficult pattern to explain" This pattern can be easily explained. If certain area has high population aging then health care services experience significant strain, so that very old groups receive less attention and lower survival.

**SJN:** This comment neatly illustrates the extraordinary gulf between my own logic, and the logic prevalent extreme-age research. Why would significant strain on a healthcare system cause *more* 105-year-olds per capita? Or, if you prefer, why would a higher survival to age 90 be indicative of a healthcare system suffering 'increased strain'? What evidence is there for this explanation?

To me the comment seems unattached from the normal constraints of a healthcare system, but perhaps a 'worse healthcare systems will cause more 105-year-olds' argument makes sense to others. I don't see it, and absent some remarkable or robust evidence I will not be including such an argument in my paper. If the reviewer wishes, I welcome any evidence the reviewer could muster to support such a position if they choose to make their theory public. Or perhaps I have simply misunderstood, and the reviewer could provide more detail on how this process is meant to work.

**R6:** 7. It should be taken into account that the IDL database is highly heterogeneous. Data for some countries were updated while data for the U.S. were not renewed from the previous version.

**SJN:** This is one reason for examining patterns within countries, which ignores heterogeneity between countries. It is also the reason for using two databases, the GRG and the IDL, both of which suggest astounding levels of error.

**R6:** 8. The author should use caution when comparing life expectancy calculated for small areas. Demographic methods used in this case typically model survival after age 85 years rather than calculate them from observed mortality.

**SJN:** As noted above these are exact counts, not models. The reviewer has misunderstood the data, and I have sought to make this clear in the manuscript. I have, in addition and as always, used caution in all my comparisons.

**R6:** 9. Non-documentary methods of age determination (amino acid chirality, etc.) are not very accurate either and are based on rather weak correlations.

**SJN:** For now, yes, but this is a call for developing better measurement than any that now exist. These methods also have the benefit of normally distributed random errors, which demographic methods lack due to the nonrandom processes outlined in the paper and earlier[22]. This means that even noisy age estimates can, at least, be relatively unbiased. Please see further discussion with reviewer 2.

**R6:** 10. Many URLs in the references do not work.

**SJN:** Please tell me which URLS are not working, and I will fix them or find a more stable link. Link decay or 'dead links' is an internet-wide problem that is beyond the capacity of the author to fix, but I shall do my best to find a replacement if you would highlight the links required.

**R6:** 11. The author claims in the Abstract: "Finally, the designated 'blue zones' of Sardinia, Okinawa, and Ikaria corresponded to regions with low incomes, low literacy, high crime rate and short life expectancy relative to their national average." This is not the case for at least Okinawa, which up to 2005 had the highest life expectancy among regions of Japan. For example, in 2005 life expectancy at age 65 for Okinawa women was 24.86 years compared to 23.42 for Japanese women. For men in 2005, life expectancy at age 65 was 19.16 versus 18.33 in Japan [12]. This sentence should be corrected.

**SJN:** The study placing Okinawa at the top of the table for life expectancy uses data from the Japanese Ministry of Health, Labour and Welfare (Poulain[23] Table 2): the same data that was revealed to be 82% dead people. I do not consider these data useful, reliable, or valid.

As noted in the manuscript, these figures are also drawn from data that predate the cleanup of massive error rates in Okinawa. Other contemporary estimates of life expectancy for Okinawa in 2003, a year before the Blue Zones study, place Okinawan life expectancy at birth at 26$^{th}$ of 47 prefectures[24]. Again, these estimates predate any statistical clean-up caused by discovering most centenarians were dead, and yet still show at best wholly unremarkable numbers. I have included this citation in the study for you.

Finally, even if I accepted the figures for Okinawa as accurate (I do not, and clearly detail why in the manuscript), the sentence as it stands would remain an accurate general categorization of the Blue Zones: 80% of which would still have short life expectancy relative to their national average. As for the poverty, and the other social indicators, these statistics are cited throughout the paper. Okinawa is again clearly last in Japan, with double the poverty rate of the next-lowest province.

**R6:** The author attempts to argue that most cases of supercentenarians are false by using indirect methods and referencing older studies on centenarian age validation. However, a recent direct study of age validation for semisupercentenarians and supercentenarians was conducted for three birth cohorts (1898, 1900, and 1902) using the Social Security Death Master File ("Mortality Analysis of 1898-1902 Birth Cohort" by the Society of Actuaries). Unlike earlier studies, this research benefited from computerized data for many government records. Validation was achieved by linking each centenarian record to early data sources, with records considered valid if confirmed by multiple sources from before 1950. The number of unconfirmed cases increased after age 109, accounting for 30-40% of all cases. Therefore, it is accurate to say that about half of all supercentenarian records cannot be confirmed, but the other half represents validated cases. Thus, it would be incorrect to claim that all supercentenarian data are fraudulent, and these claims should be revised.

**SJN:** I have not only debunked validated claims, including the oldest man in the world Kimura/Miyake (validated in 2017), I have also demonstrated throughout the manuscript that validation is not a method that can eliminate errors or be trusted on any level. I regret that this was not sufficiently clear to the reviewer. I have added some extra examples to reinforce the repeated systemic failures of the field.

These new cases reinforce the repeated finding that demographic 'validation' is not a robust method for detecting errors – worse, it is routinely misleading and biased. This is evidenced by the astronomical rates and duration of error observed in 'validated' cases subjected to the most intense scrutiny, including *e.g.* the 100-year-long 'validation' of Pierre Joubert as the world's oldest man and first supercentenarian, a case that ended in his real age of 65 years being written on a death certificate. That is, even when errors are extremely obvious, demographers have been almost completely incapable of detecting said errors (see Joubert, White, Kimura, etc etc etc). I am afraid that I cannot take seriously any defense that simultaneously admits making a 100-year-long 'validated' extremely high-profile mistake, but then claims that 'validation' is somehow a rigorous and defensible test of human age. It is not. That is a primary finding of the paper.

On top of this extraordinary problem, even when the documents are perfectly consistent, many types of error can remain undetectable. The Japanese case of >230,000 dead centenarians was not discovered via documentary evidence, which was in most cases completely consistent. Neither was the Greek pension fraud: their paperwork was fine, yet they were dead. The capacity for further undetected errors is unmeasurable.

As such, the limited concordance between the HMD database and other 'validated' data seen in the study cited is an indicator that independent documents data often says the same age, but does not provide (and cannot provide) any information on whether those consistently reported ages are also accurate.

Prof Ken Wachter admitted this state of affairs when he stated "Occasionally… a mistake will escape even a rigorous validation procedure" and such cases will remain undetected[25], yet exposed the flawed thinking of demographers when he then neglected to explain the part where he (somehow) divined the 'occasional' frequency of an undetectable set of errors. That is, how could he (or the reviewer) claim to know the frequency of an undetectable error process?

They cannot, and neither can anyone else. To pretend otherwise, without presenting evidence, seems absurd.

The claimed accuracy of 'validation' by documentary analysis is therefore merely an opinion of accuracy: one that deliberately ignores the long history, prominence, and huge number of 'validation' failures detailed in this paper, and the extraordinary frequency with which these errors can remain invisible to demographers. I have added several more such prominent cases to the paper, in which the supposedly oldest person in the world was exhaustively validated and subsequently turned out, years *after* validation, to be fraudulent.

The entire process of 'validating' ages put forward by Vaupel, Leaf, *et al.* is therefore deeply flawed, incapable of detecting obvious errors for decades on end, and based on assuming a 'low' frequency for several unobservable error-generating processes. That is the core finding of the paper, and I regret to see that the reviewer has either not understood or chosen not to understand this critical finding. I will make efforts to make the point more clearly in the text.

#### Reviewer 7

**(R7)**

Review of "Supercentenarian and remarkable age records exhibit patterns indicative of clerical errors and pension fraud" for BMC Public Health

This article examines the geographic level correlates of areas which contain large proportions of supercentenarians, those who are 110 years of age, and older ('Blue Zones').

In opposition to reasonable expectation the correlates are all the reverse. For example, areas with higher poverty, which is well known to increase mortality, have higher proportions of supercentenarians.

This article is of critical importance as It challenges a popular notion that these 'Blue Zones' serve as examples for humans on the behaviours that will lead to longer lives. Rather, these places are much more likely to be hot spots for pension fraud.

The data, it's handling and the interpretation of the results are all sound, in my opinion.

The article should be published subject to one relatively minor area of concern.

The methods section concentrates on data issues but should make the intellectual approach much clearer: here the author should be more explicit about the assumptions that the supercentenarian are either errors or fraud, and not likely genuine. The article implicitly assumes a monotonic effect on the proportions of supercentenarians in an area of literacy (positive), poverty and crime (both negative), lifespan  (positive) and birth registration (zero). This is reasonable.

But we need this to be justified with the appropriate citations. And alternative mechanisms where effects could be different dependant on the quantile of the distribution should be addressed (i.e. could some effects reduce the mean but increase the standard deviation, like some sort of positive survival effect). Can you cite studies which show that age follows a normal distribution?

Or in other words, the author needs to provide a better grounded, referenced and justified methodological setup for the first two paragraphs of their conclusion:

"Basic economic and social indicators in the modern economy, such as GDP per capita and poverty rates, predict the distribution of extreme age records. Despite constraints on model construction and accuracy, such as unavoidable differences in per capita adjustments, these basic models approached reasonable accuracy. However, the direction of these interactions is the opposite of rational expectations.

Diverse social and economic indicators that normally predict worse health outcomes, such as income deprivation, poverty, and high unemployment, are all positively associated with a higher probability of reaching an extreme age. These factors are linked to a lower probability of survival and worse health outcomes at every age below 90, for every population included in this study." (page 17).

This should not be hard, and should not take much time. But it is essential.
**SJN:** I would like to thank the reviewer very much, and I am happy to supply more detail. I have added some key citations and additional text to justify my logic here – such as the link between inequality, drinking, and mortality in the UK and USA for all evaluated ages – and hope this is sufficient.

**R7:** A table summarising the effects of each of these variables with a brief commentary, and a sufficiently detailed outline of how the regression approach used in the paper aligns with this, should suffice.

**SJN:** I agree and, while it is a little cumbersome, I have added the requested table as Table S4. I have also added some annotation to the code. I would supply the entire models and data themselves, much more openly and in line with the spirit of open science, but unfortunately personal threats from the GRG make more open data practices impossible and I am restricted to supplying these directly upon request. In lieu of this I have made the code and as much of the data and models available as I can.

**R7:** Small Comments

Page 4: "Here, we explore the possibility that extreme old-age data are dominated by age-coding errors" should be "…errors and/or fraud"

**SJN:** Agreed, thank you.

**R7:** There are small typos throughout, "It written…": top of page 20 is one example.

**SJN:** Thank you, I appreciate the effort in spotting these mistakes and I have corrected these errors wherever I can find them.

#### Reviewer 8

**(R8)**
This article starts from an undeniably important question, and proposes some very rich and impressive work on the data. What it ultimately establishes is however not clear to me. And I think it's entirely possible to work on clarifying it.

This article is a radical challenge to a major body of work studying the phenomenon of extreme ageing, centenarians and rather even super-centenarians (dying at over 105) and super-super-centenarians (dying at over 110). The author maintains that this phenomenon is an artefact resulting mainly either from errors or inadequacies in the statistical apparatus, or from fraud.

Given the number of published works implied by these results, the quality of the demonstration is a key issue. 2599 articles referenced on Pub Med appear when you search for the word 'centenarians', growing exponentially since the beginning of the century, with less than 10 per year before 1990 and more than 150 per year since 2020. What's more, the works that refer to the population of centenarians use these populations to test hypotheses and rarely describe the method used to establish ages at death. The author could have analyzed these practices and examined to what extent the progress made in identifying centenarians has been considered (including by withdrawing articles published using data that incorrectly identified centenarians).

The author is however not the first to question the scientific data on individuals with extreme longevity (we are not referring here to the existence of numerous cases of more or less imaginary characters whose lifespan is supposed to have exceeded several hundred years). And it does not really account for the existence of these works, which are old and some of which date from before the “centenarian cult” mentioned by P. Laslett[1]. For example, Leaf, Alexander. 1975. *Youth in Old Age*. New York, McGraw-Hill, Jeune B., Vaupel J., *Exceptional Longevity: From Prehistory to the Present*, Odense Univ. Press, Odense, Denmark, 1995, J. M. Robine and M. Allard, "The Oldest Human", *Science*, 20 Mar 1998,Vol 279, Issue 5358, Heiner Maier, Jutta Gampe, Bernard Jeune, Jean-Marie Robine and James W. Vaupel, *Supercentenarians*, Heidelberg, Springer 2010, the work of the IDL (International Database one Longevity) which can be found at https://www.supercentenarians.org/ and which contributed to the recent publication of *Exceptional Lifespan*, edited by Heiner Maier, Bernard Jeune, James W. Vaupel, Springer 2021. Etc.

[1] P. Laslett, « The Bewildering History of the History of Longevity » in Bernard Jeune, ed, *Validation of Exceptional Longevity*. Odense University Press, 1999

**SJN:** The paper has accounted for, and debunked the methods of, most of these works. I directly cite *Supercentenarians*, and Jeune and Vaupel *et al.*, and debunk their methods. I have been at pains to show these methods have little to no capacity to detect longstanding errors. For example, Pierre Joubert was validated by these methods, for a continuous century, before being debunked. Kimura remains validated by these methods, as of 2025, despite the case being deeply flawed. Other works are not relevant as they do not detail any methods or novel we have not discussed and debunked (*e.g.* as the full total of their evidence Robine and Allard[26] simply say, without providing any further evidence, that "*The age of Jeanne Calment was carefully verified in 1995*": of course, so were all the fake and later-debunked cases detailed in this manuscript). Finally, I have clearly detailed instances where perfectly consistent records can pass these methods of validation, which I cite, without detection. I regret not making these various citations, and the debunking of their methods, more prominent. I have included more examples and revised the text to reinforce these ideas.

To clarify these themes, I have also several reinforced prominent failures of the methods of Vapuel *et al.* in the text, stating again quite clearly how demographers have studiously ignored the decades- or century-long failure of their own methods in case after case of self-supposed extreme longevity. Carrie White is one very informative example of this, as she was the world's oldest (and most scrutinized) person before the equally 'robust' case of Jeanne Calment. Her case was examined for roughly two and a half decades, including by Vaupel *et al.*, without them uncovering that the case was a fake or detecting any inconsistencies in her many documents.

I personally find it very difficult to accept the by-now rote line that these methods (or people) are somehow capable of detecting errors, when they have repeatedly failed to do so for decades on end in the most heavily-scrutinized cases on earth. In addition, I find it remarkable that I am also supposed to trust the qualitative judgement of researchers that I have personally caught in acts that bear a striking, exact, unexplained, and inadmissible resemblance to research fraud[27] as I have with Professor Vaupel. That is, I am not obliged to trust the qualitative judgement and non-reproducible interview-based methods of people who make decade-long mistakes and cover up unwanted data[27] with deeply questionable 'coincidences.' As such, I have also added these sentiments to the text with extensive citations.

Finally, as for Professor Leaf, I am glad for the opportunity to bring it up. I had been deliberately ignored his work as it was long shown to be based on dubious records and bad data[13,14]. However, it does serve yet another instructive example of systemic failure in demography. To strengthen the above idea that current demographic methods are incapable of detecting major problems for decades at a time – like Professor Leaf's claim that a valley in Pakistan made famous by a fictional novel is a region of extreme longevity – I have now included references to the professor and his work.

**R8:** His approach focuses on the existence of errors in the measurement of the age of supercentenarians and on the analysis of the means of statistically validating these ages: his attention does not, therefore, dwell on works that seek to validate particular cases, that of Jean Calment being one of the most often cited[2], but there are many others.

**SJN**: I had thought that providing many pages debunking such cases, including the world's oldest man, was sufficient. Regrettably, it seems that the failure of demographic methods needs to be further reinforced. I have gladly expanded a detailed and clear list of examples, showing even more failures of demographers to uncover transparently bunk cases for decades at a time. A special focus has been allotted to the aforementioned professor Leaf, and to some of the very many validated-then-debunked world's oldest people.

**R8:** It focuses on 5 countries: USA, France, Japan, England, and Italy, although others could have been considered, notably Sweden and Canada, which have particularly reliable databases.

**SJN:** I sought to make the rationale for the selected countries clear, namely that they contain ~80% of all centenarian cases and that other countries such as Sweden and Canada do not have sufficient numbers for a meaningful within-country analysis. I have reinforced this rationale to make it more clear in the text. In addition, the lack of sufficient centenarian numbers in countries with 'particularly reliable' datasets is itself damning, and the qualitative assertion that records in France and England are somehow on a sufficiently lower level than Sweden and Canada as to make these cases non-comparable is very questionable. For example, French birth registration begins in 1792, and English birth registration begins in 1837: a history of vital registration that far predates the central registration of births in Sweden (1860), the beginning of Canadian birth registration (early 1890's-1900's), and even the existence of Canada as a country (1867).

**R8:** For these 5 countries, the author uses a large set of data in a controlled and systematic way, a set of data which corresponds to those used by authors working on cases of extreme longevity. It would be a considerable task to repeat all the programs used by the author to check their accuracy: we can see that the tables provided give

[2] Robine, Jean-Marie, Michel Allard, François R. Herrmann, and Bernard Jeune. 2019. "The Real Facts Supporting Jeanne Calment as the Oldest Ever Human." *The Journals of Gerontology: Series A* 74(Supplement_1):S13–20.

coherent results in line with what we know in their descriptive aspects. For example, it is a fact that birth certificates appeared in the United States at the very end of the 19th century, and the effect on the frequency of SC is striking. Similarly, it is well known that the quality of the data produced by the Civil Registry in France is lower in Corsica than in the mainland departments.

However, it seems to me that the relevant method should be to take the latest state of the art and only consider the more recent data on centenarians whose validation has been considered to be well established by the authors. For example, everyone agrees nowadays that without a birth certificate, the data is unreliable.

**SJN:** We are pleased to see the reviewer agrees with us that the vast majority of the 'validated' data listed in the IDL at the start of the study in 2019, which either have no birth certificate (over 80%) or a later replacement, should be thrown out. Also included in the list of cases with no birth certificate are the world's oldest man (Kimura), the oldest-ever man in the USA (Richard Overton), and between 59% to 95%+ of the supercentenarians in the USA. Although it would not eliminate errors, by any means, this process would reduce the global number of extreme-age records to much more manageable sizes near zero.

**R8:** The author could also have taken as a starting point the way in which validation criteria are constructed and applied today. For example, the validation procedure described in *Exceptional lifespan* quoted above takes up less than half of page 18, and even then a section is devoted to explaining that validation is carried out on a sub-sample by random selection.

The first step in the paper's statistical work is to identify in which places supercentenarians are over-represented. This is not so simple. Would it be possible to use a purely statistical method to identify the deviation of the distribution of points observed at extreme ages from the relevant statistical distribution? The idea of identifying territories where supercentenarians are over-represented corresponds to usual practice: anomalies, whether Japanese, Sardinian or Corsican, are used usually used to identify factors of exceptional ageing. This does not mean that the supercentenarians identified elsewhere are relevant. For example, are all the supercentenarians from wealthy regions really supercentenarians, or are they false supercentenarians, just individuals from poor (and fraudulent) backgrounds? In a way, the author's intention is not clear: is he trying to show that in certain populations the proportion of supercentenarians has been greatly increased, or should the very existence of supercentenarians be called into question in his eyes?

**SJN:** Yes. Clearly, the entire existence of supercentenarians is in question. As is the validity of the field, and the integrity of many of the people involved. I have reinforced this in the text of the critical review.

**R8:** It seems that what he is proposing (the author is not very firm and clear in his interpretation of the results he presents) is to refute the idea that factors of exceptional ageing can be identified by the existence of an exceptional proportion of supercentenarians in certain geographical regions, because this over-representation in fact reflects a greater propensity for measurement errors (univocal, in the sense of an overestimation of age) in these regions.

**SJN:** I have made sure to firm up my conclusions, and regret that they were insufficiently clear. I have now reinforced the text to show my opinion that demographic error rates in extreme ages are so high as to justify abandoning the field at large and, at best, starting again from scratch. I have also completed a survey of over 70 years of UN world populations prospects data, showing the same result and the same opinion[6].

**R8:** On the basis of measurements of the local rate of supercentenarians, the author identifies a link between this rate and the characteristics of the place of birth (and death) of the deceased - the question of mobility is not well addressed in the article.

**SJN:** I have addressed the problem of mobility in the article, highlighting that the majority of birth and death locations are in the same place, and reinforcing this as a limitation of the study.

**R8:** The author's main finding is that "Higher poverty and deprivation are positively correlated with all age categories of the oldest-old in every country measured" (l.323-25) and "'anti-health' factors exhibit a consistent positive association with extreme longevity" (l.424-425). Although he does not state it directly, it is on this basis that the author establishes that supercentenarians cannot be supercentenarians.

**SJN:** I have now stated directly that this means, in effect, that supercentenarians are most likely a collective statistical fantasy.

**R8:** He rejects the idea that these "anomalies" could have a relevant interpretation by reference to "unknown lifestyle factors". This observation is not in itself sufficient proof. In particular, we could analyse the hypothesis of a form of over-selection specific to this places that lead the poor to survive being more robust and hardened than the rest of the population, or the fact that they receive special attention, which protects them and increases their lifespan.

**SJN:** I can only suggest here that the author re-read the text, which in fact states “these anomalies may, perhaps, be explained away by reference to unknown lifestyle factors” in the quoted passage: *i.e.* the exact opposite to the statement here.

**R8:** In general, the author does not give sufficient consideration to what makes supercentenarians (or super-supercentenarians) special age groups in terms of mortality, where the effect of age plays out differently than it does for centenarians, nonagenarians or the elderly in general.

**SJN:** I give exhaustive consideration to why supercentenarians are a ‘special’ age group. That’s why I conclude these data are a special collection of statistical absurdities. Because to do otherwise would require I pretend that smoking and poverty is good for survival. Which they are not. I have also added further consideration of, for example, urban-rural divides.

**R8:** Furthermore, if the over-representation of supercentenarians is linked to the propensity of the poor to commit fraud, its effect is stronger in poor areas but should also be present elsewhere.

**SJN:** I don’t understand the point I am supposed to address here. I have already measured ‘elsewhere’ by capturing all the regions of each country. The ‘effect’ of poverty is not being coded as present or absent, but as a continuous variable, and rich, poor, and middle-class areas are all being measured. Rich areas have less 100+, 105+, and 110+ year olds per capita, poor areas have more. If old-age poverty leads to pension fraud, the higher old-age poverty in poor areas will be associated with more extreme age records: a pattern borne out by the evidence.

**R8:** Finally, it would be interesting to find out whether there are poor areas where such over-representation does not exist.

**SJN:** I examined every region in Italy, Japan, England, and France. The data are all available, and as you can see in the figures, and when all areas are comprehensively evaluated the answer is no.

**R8:** To reinforce his thesis, the author looks at the precise date of death to the nearest day and finds that the data is again flawed for centenarians, who do not die randomly every day of the year but on dates that have been 'manufactured'. The observation is correct, but does it constitute proof of the inaccuracy of the age of these supercentenarians in years? More generally, the author does not discuss the extent of the error: the day was lost but the year was correct or, as in some cases, the father and son or mother and daughter were confused and there was an age difference of around 30 years?

**SJN:** I will first note that the age-heaped data describe the date of birth, not death. I think this is just an honest mistake by the reviewer and I will take the comments as if they meant 'birth'.

I would thank the reviewer for acknowledging the substantial flaws evident the day of birth. However, these undetected errors do not just indicate enormously flawed distributions in the day. They indicate the inadequacy of the validation method. The relevant question here is not, therefore, 'what about the year', but rather: why were these dates promoted as near-infallible outcomes of a strictly accurate validation method, when they closely resemble an assemblage of errors, without anyone ever noticing?

That is, for decades the 'exhaustively validated' people in these databases were claimed to have completely accurate and validated ages: a 'gold standard' accurate to the exact day, month, and year. Showing clear evidence that the day of birth is inaccurate therefore completely upends long-standing claims that such a validation procedure can detect errors.

Accepting that the days are junk data, but retreating to make a claim that the year *could* be accurate, would therefore require an untenable motte-and-bailey argument that deliberately ignores the point I intended to make and the wholesale disaster of the tested data.

To ensure the point is very clear, I will repeat myself a little: the people validating supercentenarian ages have made several decades worth of assertions that their dates are 'validated' and accurate for day, month, and year alike. This is false and it does not seem possible to trust these data at all, which the reviewer appears to see and admit. So, as the birth dates are inaccurate: 'exhaustive' validation has been incapable of detecting age errors for decades on end, the validation method has not worked, and the database is unusable.

There appears to be no simple way to escape this conclusion. I have tried to make this distinction clearer in the text. It is the method that is invalid: the substantial age-heaping anomalies in days of birth are just the accessible and testable framework in which that inaccuracy and inadequacy is revealed.

The worst outcome of this failure is, however, that a retreat to an 'accurate year' defense would appear disingenuous and ignore important questions. First, if a method of validation claimed as accurate and valid cannot spot the wholesale failure of days, for decades on end, why would you trust the methods for any further analysis? Second, if demographic research is to be considered trustworthy at checking its own data, then why has arguably the oldest and most basic check of data validity in the field – age heaping – never been applied to these data before?

I have therefore made sure to raise these points in the text, clarifying why a 'motte-and-bailey' style retreat to the year of birth would be inadequate to explain this decades-long methodological and cultural failure.

**R8:** The author therefore highlights two types of source of error linked to age manufacture: poor quality of data recording at the time of birth and fraudulent manipulation of dates at the time of death. The correlation between these two types of reason and the poverty of the location is not obvious. On the other hand, it would be worth investigating the actual prevalence of either of these phenomena and their geographical distribution.

**SJN:** There is regrettably no way to measure the prevalence of either phenomenon, for reasons detailed in the paper. If there was, every government on earth would not be struggling with epidemics of identity theft.

**R8:** Finally, the article looks at the reasons for the persistence of a systematic and serious error in demographic research. In the discussion, it makes two contradictory points: the first is that there have been many proven cases of measurement error in the supercentury and that demographers have not reacted in any way. The second is that there is no internal means of detecting such errors. The imaginary example of an identity substitution between an individual and his (younger) brother, who would have borrowed the former's documents (or even his memories, should he be questioned) illustrates this possibility of errors that are undetectable using the usual historical sources.

**SJN:** I regret that identity substitution is not an imaginary example, but a source of error that has remained well known in demography for centuries[28]. I have included a concrete example of exactly this type of substitution in extreme-age records: three men that remained undetected by demographers for a considerable number of years, despite being interviewed and validated as claimants to 'world's oldest man'. That said, this error pattern is only one of the many diverse types of error that demographers remain incapable of either detecting, measuring the frequency of within a population, or correcting.

**R8:** It does not seem entirely convincing insofar as the possibility of an ever more in-depth investigation into individuals of extreme ages seems, at least in theory, always possible, the existence of a younger sibling being a possible trigger, for example.

**SJN:** Over 98% of supercentenarians are dead. Many of the rest either refuse requests for interviews or are suffering debilitating mental illnesses that render them incapable of coherent answers. I am therefore curious how the reviewer might engage in 'ever more in-depth' interviews of the deceased, or indeed merely the dementia-suffering, in theory or practice.

Added to this problem, I have reinforced in the text the strikingly low rates at which interviews are ever conducted even once. For example, of the more than 9,878 French SSCs (105+ year olds) only 365 - or 3.7% - were ever 'exhaustively validated', and an unknown (but smaller) number appear have been ever interviewed. As these interviews are virtually never reported, it is also not possible to even measure the concordance between post-hoc evaluation of records, and the subjects are seemingly never subjected to double-blinded repeat interviews that would tell us the accuracy of the qualitative assessment of demographers. In addition, while demographers often directly and personally benefit when a case is validated, there seems no oversight of the process whatsoever, and no recordings or publications of the interview required or expected by the field. As such, I have now highlighted the unreliability and general irreproducibility of this method in the text.

**R8:** Systematic data on the basis of nominative data matched to vital data, census data and military conscription data for men at least, would make it possible to check the ages of supercentenarians.

**SJN:** I find it striking that such a solution is proposed, after I have demonstrated how conclusively this statement is not true. Systemic data matching is exactly the method that is claimed to have been used to generate extreme-age data in the first place and it has systematically failed. It is not possible to 'check the ages' of these people, as cross-checking ages does not tell you if those ages are accurate. The evidence for this is the the hundreds of thousands of dead people with no detected paperwork-errors described in the manuscript, and the many instances in which cross-checking paperwork has 'validated' a case that subsequently turned out, with a delay of up to a century, to be bogus. The thousands of dead centenarians of Greece that I reveal in this paper did not have any known errors in their paperwork: they were all dead with functioning, consistent, pension-paying records. I am astounded that the reviewer has chosen to ignore this basic historical fact, and many others, to continue with the demonstrably unreliable line of reasoning.

It ignores the simple veridical truth that cases with perfect paperwork are routinely validated (Carrie White), *and* that cases with extraordinarily flawed paperwork in which 'rigorous cross-checking' results in a spurious case being validated for indeterminate periods of time. Checking the paperwork does not work. It has never worked. If you doubt that, I suggest you look for the 230,000+ dead centenarians with impeccable paperwork who were living in Japan *circa* 2008. I do not know how else to frame this, and I find it incredible that these historical realities are being ignored.

**R8:** In any case, the relatively small size of the populations concerned means that such in-depth counter-analyses can be envisaged.

**SJN:** I describe well over a quarter-million deceased centenarians in this paper, most of them found to be errors after being accepted as 'amongst the best' data in the world. Personally, I do not regard this population as small. In addition, there is no evidence that revalidation or 'counter-analysis' of this dataset will produce results any different than the first round of validation and analysis: namely, a collection of junk. Virtually none of these cases were discovered by a demographer. Most were discovered by city health officials and auditors. Placing the 'revalidation' of such populations back in the hands of demographers, who have so utterly failed at their own allotted task, seems wasteful and counterproductive. These results suggest the need for resignations and repercussions, not reanalyses.

**R8:** Of course, the alternative of reliable biological measurements is tempting, but we would need to have a biological marker that has evolved since the birth of the individual in a way that is perfectly independent of the individual history of its carrier, which does not seem obvious a priori.

**SJN:** Fortunately, the rate of radioactive decay and the racemization of chiral amino acids is not, to my knowledge, dependent on lifestyle choices.

**R8:** In any case, this point leaves unanswered a fundamental question posed by the author: "It seems worth asking, therefore, why the recurrent discovery of high rates of non-random age errors have been ignored by the scientific community, given the fundamental importance of accurate age data to fields like medicine, gerontology, the social sciences, and epidemiology". This question is tantamount to acknowledging that the subject of the article, identifying age measurement errors, has long been the focus of attention in the scientific community. In any case, the article lacks a socio-historical analysis of the formation of the international research community, which is undoubtedly quite small, bringing together researchers who have made the study of extreme ages a specialty, a separate field of research. And it would be interesting to know the scientific and non-scientific uses that are partly linked to these works.

**SJN:** I cannot agree that "identifying age measurement errors, has long been the focus of attention in the scientific community" with any degree of success, when the disappearance of thousands of Greek centenarians does not produce a citation. Despite the newspaper articles, there does not seem to be a single demographer that has discussed or even cited the wholesale disappearance of Greek centenarians. Nobody discusses the effect of 30,000 dead Italians either. There seems to be only one paper that handles the disappearance of over 230,000 Japanese centenarians occurred. This work makes excuses that are, to my mind, so detached from the reality of the underlying failure of the research community to spot an 82% error rate that the paper is hardly creditable.

I therefore cannot accept that the focus of the demographic community has been the honest detection of errors, given the disappearance of entire populations has been ignored. These instances are barely, or never, cited. They certainly do not generate the required level of soul-searching that an 82% failure rate to detect deaths would cause in, for example, a clinical trial. If 82% of the money in a till were to disappear in a café, the waiter would be sacked that afternoon. In demography – a field that proclaims rigor and accuracy right up until the subject is discovered to be dead or pretending – nobody has even noticed. As such, I stand by the statement, because it is supported by historical realities of a field riddled with systemic failures that I have cited at length.

**R8:** All in all, the article certainly deals with an important issue for population demography, but as it stands at least, more as to be done for it to be fully convincing.

#### Reviewer 9

**(R9)**
The manuscript examines the phenomenon of supercentenarians - people over 110 years old. The authors question the evidence for existing records of extreme longevity. They argue that many of these records may be due to errors and possible fraud rather than true longevity.

The following aspects are examined in detail

Geography Supercentenarian status is linked to the absence of vital registration, particularly in the USA, where the introduction of birth certificates led to a significant decline in recorded supercentenarians. In countries such as Italy, England and France, supercentenarians are often found in regions with lower incomes and higher poverty rates. This is counterintuitive and leads to the question of SES and longevity: the author found that higher levels of poverty, lower per capita income and poorer health outcomes correlate with increased numbers of supercentenarians. This is inconsistent with most previous evidence that low SES is associated with poorer health! So the claim that the data sources are not reliable seems to be true.

The author also found a strange pattern suggesting data anomalies, where supercentenarians' birth dates are concentrated around just a few days, suggesting manipulation?

Overall, the findings raise serious questions about the reliability of demographic data on extreme ages.  The author suggests better methods of age verification, such as biometrics.

The author also stresses the need for better methods of age verification, such as biometric measurements.

The author used linear mixed model regressions to fit probabilities of survival and life expectancy estimates as dependent variables, with various socio-economic indicators (such as poverty rates and employment rates) as independent variables.

Overall, this is indeed a very interesting and necessary manuscript, because if the data on very old people are inaccurate, this may change outcomes and conclusions! This is particularly important as the number of cases is very small. Inaccuracies in such a small number of cases can have a big impact on the results!

I have no further criticisms other than to ask for a more detailed specification of the models used in the main text of the MS.

**SJN:** I would like to thank the reviewer very much, and I am happy to see they are satisfied with the problems raised by the manuscript. It is perfectly reasonable to ask for greater model specification and I have included an extra layer of detail in the supplementary materials as Table S4. I regret not simply being able to supply all data and models alongside the code, to adhere to the principles of open science, but I am afraid a series of personal threats from the GRG have rendered this inadvisable – I am, however, happy to continue supplying such scientific material directly in private.

### Subsequent Responses to the Editor

#### Response 1

**Sent August 2025**

Dear Natalie,

I would like to thank you for the extraordinary care it took you, over the course of six months, to come up with all four of these dot-points. In other circumstances, it would look like you did nothing for an extraordinary period – despite this being one of the most talked-about papers in demography for the past ten years.

However, given the extraordinary time it took to generate this short reply and the repeated polite requests I have made for any update, I can only assume you made extreme efforts to come up with the insightful response that you do not like my tone, and while you and the nine peer reviewers cannot raise any clear scientific objections, that I must rewrite the paper. After seven years of waiting for a logical or scientific response to my critiques, raised in the most public way imaginable, it is a warm balm to see you engage with such depth and precision, and then condense your response into something so short that it resembles a last-minute text message.

What is unclear, however, is the following problem.

You see, for an outsider it may seem that clear instances of long-term unchallenged racism – scientific racism that affect the content of a scientific debate – and the apparent fabrication of evidence is being met with a demand to remove my critical tone. As science is built on criticism, and as both the tone of this paper and the criticism contained in it have been met with such extensive press and a comic award, this presents a problem.

At least six of the nine reviewers strongly support publication with little or no further revisions, and a seventh supported publication with slightly larger workload, all of which were carried out in February. These nine peer reviewers have not articulated any meaningful problem with the content of the analysis or raised any meaningful alternative explanation for these ‘anomalous’ patterns, and neither have thousands of members of the scientific community who have offered public views on the paper.

What, then, can you hope to say to anyone I must send your way? What reason shall I give them for your requests that I remove criticism, change the title, and thereby provide cover for racists (in the most clear and direct sense of people who use race as a scientific argument) and people who have all but admitted to fabricating their evidence?

I would appreciate it if you made it clear. I would not like to disappoint anyone with the news that this paper has been held up for so long with no answer or justification, due to its politically embarrassing nature, without any scientific, logical, or moral rationale.

To really help you anticipate the questions from the community, let me suggest to you some of the inevitable questions that may follow, and you can try and provide me with some answers.

1 - Why did you spend six months sitting on a paper that was strongly endorsed by at least six of the nine reviewers, and which in your assessment did not require further review? As a reminder, in your own words back in June, you said "Given the nature of the initial reviews and the quality of your revision, the editor is currently assessing whether an additional round of peer review is necessary." Clearly it was not, as you did not send the paper back to review, and at least six reviewers were strongly in favour of publishing the paper.

2 - What would you say that you have been doing for the past 190 days or so? Why have you not answered the repeated requests I have made for an explanation of the lack of apparent action or effort to date? How can you explain your response as adequate or meaningful, to other people?

3 - Why did the paper require nine reviewers and nine published reviews in the first place and, since the paper has passed a super-majority these reviewers' critiques, what justification can you offer for ignoring the efforts of these professionals, and adding to the extensive delay in publishing a paper of extraordinary interest to the community after it has passed review?

4 - Why have you not articulated the passages I am required to rewrite, and why? Stating, as you do, that I should change my tone seems rather inadequate. Which part, exactly, am I supposed to change here?

Perhaps it is the part where an entire scientific field overlooks the fact that 82% of Japanese centenarians, which were their subject of study, were dead? Perhaps the part where a major scientist spent 20 years stating, unchallenged, that black people survived longer because they were 'more robust'? The part where the world's oldest man is clearly a fake? Or the part where an entire Blue Zone was 'discovered' to please a magazine editor, without a single solitary demographer calling out, or seemingly even recognising this level of fraud for two straight decades?

Perhaps it is the part where a French demographer states the distribution of supercentenarians is due to the biological 'benefits of slavery', and nobody says a thing?

So, please be explicit in which parts you want to defend, privately and publicly, and I will inform interested parties of the passages that mean this paper cannot see the light of day, unless I agree to your request to delete my critiques, after passing nine peer reviews.

It will require some extraordinary bravery on your part to ignore six or seven reviewers and side with such views. I look forward to your excellent justification for doing so.

In the meantime, allow me to respond to the four dot-points you have raised here, and address the more general issues that remain unresolved. Your bullet-points are marked in bold.

---

**(1) Please reframe speculative claims as hypotheses. This request also includes updating the manuscript title**

Perhaps you could help me define, as I am unclear on the distinction, what separates a 'speculative claim' from a hypothesis?

It does not seem to be an absence of evidence – I have presented over a hundred pages of qualitative and quantitative evidence here, including open and reproducible code, and (for example) the discovery that many of the world's oldest people were dead at the time of study.

Is a speculative claim separated from a hypothesis, if it is merely the type of idea you do not like? Is it an idea you find uncomfortable? Or perhaps the 'speculative claim' is something that you wish wasn't supported, despite abundant evidence that has not been successfully challenged here by any of the nine reviewers (or indeed yourself), but which you feel better about if you hide it under an adjective?

I could try and rely on the assessment of other scientists as to what is 'speculative', of course, but given recent events this approach seems unreliable. Many problems I raised were initially dismissed as wild speculation but have since found their footing – in the teeth of much harrumphing by esteemed demographers.

Take Loma Linda – a gold-standard case that it would have been wild speculation to dismiss for almost 20 years - until it was admitted that Loma Linda was generated to please an editor and the discoverer 'never bothered' to remove it. Or if you prefer, take the response to the near-complete absence of birth and death certificates in the Max Planck's supercentenarian records – which prompted, not an answer, but a rapid elimination of the underlying data from public view by the Max Planck, when they deleted the virtually-empty 'BCERTIF' and 'DCERTIF' columns from their database without explanation.

Is this the 'speculative claim' stuff? Because it looks embarrassingly well supported.

So, assist me please.

**(2) Please clearly distinguish validated evidence from conceptual illustrations**

I have been careful to make the distinction throughout the manuscript by including only validated evidence in the figures, code, and text. There are no conceptual illustrations. It's all data, supported by open code.

**(3) Please temper any provocative language used within your manuscript**

What? Why?

To shield the powerful and answerless from my pointy critiques?

If the poor dears want it put more gently, they should answer the questions before six and a half years go by. Their silence is utterly shameful. So, no. I won't take my criticisms elsewhere. I won't tone it down. I won't take out any provocative language.

I would rather enlighten some of the bloviating lipid-bilayers I've encountered as to the specific nature of their intellectual failings, but life is full of regrets I suppose, and I'll have to settle for leaving the paper unchanged.

**(4) Please explore alternative explanations for anomalous data patterns**

Maybe I can suggest it was a simple lifestyle in touch with nature, vegetables, close-knit communities, and moderate exercise. Of course, that would mean that the

600 million subsistence farmers and 3.5 billion rural residents of the world would also be supercentenarians at remarkable rates, and would invert the relationship between poverty and survival observed in every country on earth. So maybe it's something else.

Unfortunately, except for the explanation that this data is fantasy-enriched junk, no workable explanation has been raised by the many thousands of people who have seen the study so far. I'm open to it.

For example, if anyone has an explanation for why extremely-old people would around 20-70 times be less likely to have a birth or death certificate than the general population, I'm all ears. Perhaps it is their remarkable health – or *e.g.* their obviously plant-based lifestyle of raising, butchering, and eating sheep in Sardinia – makes their paperwork evaporate.

I can't suggest anything besides this being garbage data myself, and therefore have nothing more to add.

---

Please send me a prompt reply to my earlier questions, as I will not be revising this paper further and ask that it be accepted or rejected as it is. As you wish, I leave it with you.

With long-suffering regards,
Saul Newman

#### Response 2

**Sent October 2025**

Dear Natalie,

I am somewhat disappointed to discover that you seem to have discovered several new and interesting ways, by which the same and worse-supported ways, to raise objections to this paper after more than 18 months of consideration. Like your previous thoughts, I find these staggering in their novel approach to thought-processes. As for the time taken for these ideas to gestate, I can only imagine that perhaps, like vendor of a mouldy cheese, you find the length of time I took them to develop a selling-point of their quality.

Even though I am having to repeat myself, I detail a brief response to your recycled points here:

No, I am not changing the 'tone' because you do not like it.

At the risk of opening the gate to a new round of response, in which I must regard your thoughts on this subject yet again, I supply a brief reply to your further remarks below.

**Editor:**

Many claims in the manuscript, particularly those implying widespread fraud or systematic identity misattribution among supercentenarians, are presented as fact, despite being largely speculative. Examples include statements suggesting that "every supercentenarian is an accidental or intentional identity thief." Such assertions exceed what the data actually demonstrate and are likely to be perceived as sensational rather than scientifically validated.

**SJN:**

It is a shame you only quoted the second half of this sentence to make your point, and did not read that first half of the sentence (what a shame!) when stating I was passing of hypotheses as facts. You see, I was craftily hiding something there. The first half of the sentence states I was proposing "a deliberately provocative, if seemingly absurd, hypothesis: every supercentenarian is an accidental or intentional identity thief". A hypothesis, as most undergraduates know, is not a fact.

If you require assistance with reading comprehension, there are many online services and tutors available to you.

In addition, I have not implied widespread fraud. I have conclusively demonstrated the existence of widespread fraud, or cited striking instances of it happening. This is not speculation, but historical fact. These include, but are not limited to, the Greek case where 72% of the centenarian population were dead, and Loma Linda, which was generated to please an editor. What is it that escapes you in understanding that?

**Editor:** The paper relies almost exclusively on documentary records to evaluate extreme-age claims, yet it acknowledges throughout that these records are prone to errors, misreporting, and gaps in documentation. While you have correctly identified important limitations, the manuscript often conflates observed anomalies with proof of fraud, which is not justified.

**SJN:** I have provided repeat proof of fraud, including admissions by leading researchers that they came up with and 'never bothered to delist' Blue Zones.

Furthermore, it is a major theme of the paper that these data are unreliable because they have no other means of validation than paperwork itself, which is deeply and routinely flawed. Again, I must wonder what paper you have been reading, or having an AI summarise for you, to arrive at these conclusions.

**Editor:** Some of the paradoxical associations (e.g., higher extreme-age frequency in regions with worse health outcomes) may reflect reporting biases or errors rather than biological or social mechanisms. The manuscript does not provide sufficient analysis to distinguish these alternatives.

**SJN:** There has been no rational explanation for these patterns presented in seven years of scientific discussion, including by the nine reviewers and now two editors at this journal. There is no other realistic explanation – these data are garbage. Present a rational explanation or stop demanding that I have left something out.

**Editor:** The discussion contains emotive language and provocative statements that undermine scientific neutrality. While the subject is inherently controversial, maintaining objective and measured phrasing is essential for credibility.

We believe the manuscript could make a valuable contribution provided you are willing to revise it substantially.

**SJN:** I can say whatever I want in a discussion if it is supported by the data. Can you print findings that you don't like, or will you continue with these objections by finding yet another 'mouldy cheese' aged opinion?

Personally, I have no more to say than what I've already told you, at length and ad nauseum. The only real question left is whether you have the miniscule amount of courage or backbone required to publish the paper, because I will not revise it.

Your remaining suggestions are rather repetitive of those above.

**Editor:** Suggested steps include:

Reframe speculative claims as hypotheses and clearly separate proven evidence from conceptual illustrations (e.g., treat the "Italian sibling" case as illustrative, not evidentiary).

**SJN:** Again, it says "seemingly absurd hypothesis" in the description of this problem. I can underline it if you like. Perhaps a nice highlighter?

**Editor:** Temper language throughout to reflect uncertainty and avoid sensationalism.

**SJN:** I am Australia, so I must inform you this is the most polite and least colourful language my people possess. I didn't even swear once.

**Editor:** Discuss plausible alternative explanations for anomalous data patterns, such as census or record-keeping errors, demographic shifts, migration, or sampling biases.

**SJN:** There are none, or at least nobody has presented any – not yourself, not the reviewers, nobody.

**Editor:** Highlight the potential role of biological age verification methods as a complement to documentary records, without implying universal error.

**SJN:** Biological age verification is calibrated off paperwork ages. It's like using one ruler to mark out another using a pencil. If the first one is wrong, so is the other. Measuring the two rulers against each other does nothing to resolve this issue.

**Editor:** Specific points for addressal include the below:

Controversial statement (quote or paraphrase)
"Every 'supercentenarian' is an accidental or intentional identity thief, who owns real and validated 110+ year-old documents, and is passably good at deception."

Why it's an issue
This is a sweeping, unsubstantiated generalization implying universal fraud. It's speculative and framed as fact without empirical evidence. It risks being perceived as sensationalist and damages the manuscript's scientific credibility.

**SJN:** This is a logical argument. A clearly delineated thought experiment, demonstrating how universal fraud is possible to remain undetected. Nobody, anywhere, has presented a counterargument, and I do not believe you have one either.

**Editor:** Controversial statement (quote or paraphrase)
"It is therefore unclear why only 8% of all SSCs and 1.4% of all US supercentenarians recorded in the IDL have a death certificate unless these data are mostly errors."

Why it's an issue
The low certificate rate is a red flag, but concluding "these data are mostly errors" overreaches the evidence. Missing documentation can have non-fraud explanations (data privacy, record loss).

**SJN:** And why would these rates of "data privacy, record loss" be 70-fold higher in exhaustively validated cases, compared to the population from which they were sampled? Additionally, data privacy is not at all relevant when using comprehensive register data. These people are dead, and they have no death certificates. So what would be the argument? It's not like they said "nah I opt out of death certificates for privacy reasons".

As I said, for several previous 'issues' you raise, nothing in this makes sense, and you and your nine peer reviewers have not presented any alternative that makes sense.

**Editor:** Controversial statement (quote or paraphrase)
"Observed anomalies... support a primary role of fraud and error in generating remarkable human age records."

Why it's an issue
Suggests causation where only correlation exists. Without case-level proof of fraud/error, this claim overstates what the analysis can establish.

**SJN:** Everything about this suggests fraud and the general scientific community and (I grow exhausted) so do your peer reviewers.

**Editor:** Controversial statement (quote or paraphrase)
"Extreme-age demographers seem to have misunderstood... acceptance of such cases is dependent on... arbitrary generation, survival, and detection of documents."

Why it's an issue
This criticizes an entire scientific subfield in general terms without acknowledging improvements in demographic validation or alternative interpretations, which weakens objectivity.

**SJN:** There are no methodological improvements, none, that can detect these errors. That is the point. In addition, the specific terms by which I have criticised this entire subfield are contained throughout the paper, and remain collectively and individually unresolved and unanswered.

**Editor:** Controversial statement (quote or paraphrase)
"Every document in the room is consistent, real, and validated. Any such type-I error therefore has the potential to indefinitely escape detection."

Why it's an issue

While conceptually valid, this implies that any validation can be false, which undermines all existing demographic data without direct supporting evidence.

**SJN:** It does not *imply* any validation can be false. It states it categorically: because it is true. Every validation has the capacity to be false, and there is no guarantee of human ages if you only use paperwork. Again, where is your counterargument to this basic logic from yourself or anyone else? I have not seen it, and nobody has raised it with me or the general scientific community after seven years.

We have assumed that errors were low frequency or randomly distributed, and so could be ignored, yet this is not true – as my theoretical work has demonstrated.

**Editor:** Controversial statement (quote or paraphrase)
"Claims to remarkable regional longevity... are contradicted by independent studies."

Why it's an issue
Reasonable observation, but phrased too conclusively. This needs careful attribution and discussion of differing methodologies between datasets.

**SJN:** Can reasonable conclusions not be stated? Because I have seen no studies, anywhere, that employ a method that can detect, or did detect (in the Greek /US/Japanese etc cases), high frequency obvious frauds that were discovered later.

**Editor:** Controversial statement (quote or paraphrase)
"Document scrutiny cannot detect routine errors... completely fake cases can remain 'validated' indefinitely."

Why it's an issue
Overgeneralizes a genuine issue. Some errors may pass validation but implying that all document validation is futile is too absolute.

**SJN:** I love this circular logic. If errors can remain indefinitely undetected, how did you conclude only "some" of them exist? Magic? Wishes?

These errors are, by definition and as you admit, undetectable. Yet you remain sure they are not there at a high frequency - even when I have illustrated cases where they were virtually the entire population. How did you claim to know the frequency - the 'some' in 'some errors' – of errors that are undetectable?

What is your escape-route from that circle of reasoning?

**Editor:** Controversial statement (quote or paraphrase)
“The global pattern of centenarian data reveals universal problems… under such models, the majority if not all remarkable age records may be errors.”

Why it's an issue
“Majority if not all” is a strong and unproven claim. It extrapolates local, verified issues to a global population without systematic evidence.

**SJN:** I made the counterpoint to this above – that I have pointed out >80% undetected error rates in some of the ‘best’ data in the world, in a diverse set of globally distributed countries. In addition, I have cited a global study measuring every centenarian on earth. It is all strikingly problematic: these issues are systemic, globally and without exception, and do not evaporate when they cross a border.

**Editor:** Controversial statement (quote or paraphrase)
“Epigenetic data uniformly predict the oldest-old are 8–20 years younger than their paper records suggest.”

Why it's an issue
This treats emerging biomarker results as settled fact. Epigenetic clocks are not yet validated for extreme ages, so the discrepancy could reflect limits of the method, not fraud.

**SJN:** It does not, and the logical statement is valid as it stands. There is, in addition, no paperwork-free metric to validate these measurements against.

**Editor:** Controversial statement (quote or paraphrase)
“We can only hope that… new biometrics… may end our long reliance on documents.”

Why it's an issue
Emotive, editorial tone (“we can only hope”) breaks scientific neutrality. This should be reframed in neutral language emphasizing complementarity, not replacement.

**SJN:** Why compliment methods of age measurement that do not work, at advanced ages, when you can replace them with something that does?

In summary, all the answers above rely on excluding logic or reasoning or evidence because it is ‘controversial’. Why controversy should exclude logic or evidence from a scientific discussion is not made clear. Equally unclear is why your personal, qualitative perception of what is and is not controversial should have a bearing on what is true.  You have not presented any logical counterarguments and rely on “we think this should not be said” as a response. That is not science. It is opinion.

So, I must say again: no. I will not revise a word, and you shall either accept this paper as-is because it has passed peer review, or you shall show that your scientific selection process depends, entirely and without shame, on your personal bias.

With even longer-suffering regards,

Saul Newman
