## Supplementary Materials for "Supercentenarian and remarkable age records exhibit patterns indicative of clerical errors and pension fraud"

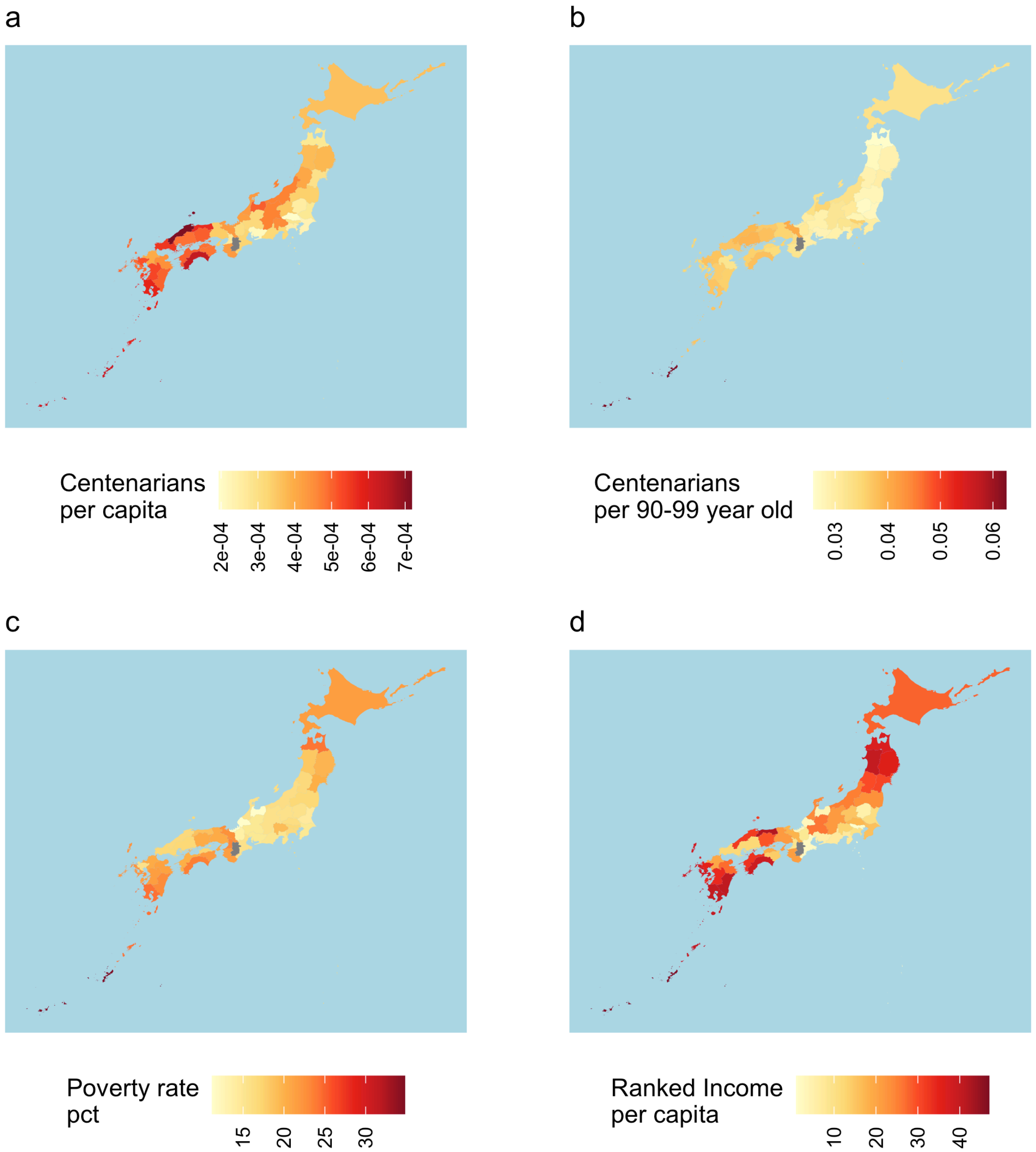


**Figure S1. Poverty and the distribution of Japanese Centenarians.** The 47 prefectures of Japan putatively contain over 48,000 centenarians, with generally higher concentration of centenarians per capita (a) and per 90-99 year-old (b) in prefectures with high poverty rates (c) and lower-ranked prefectural incomes (d).

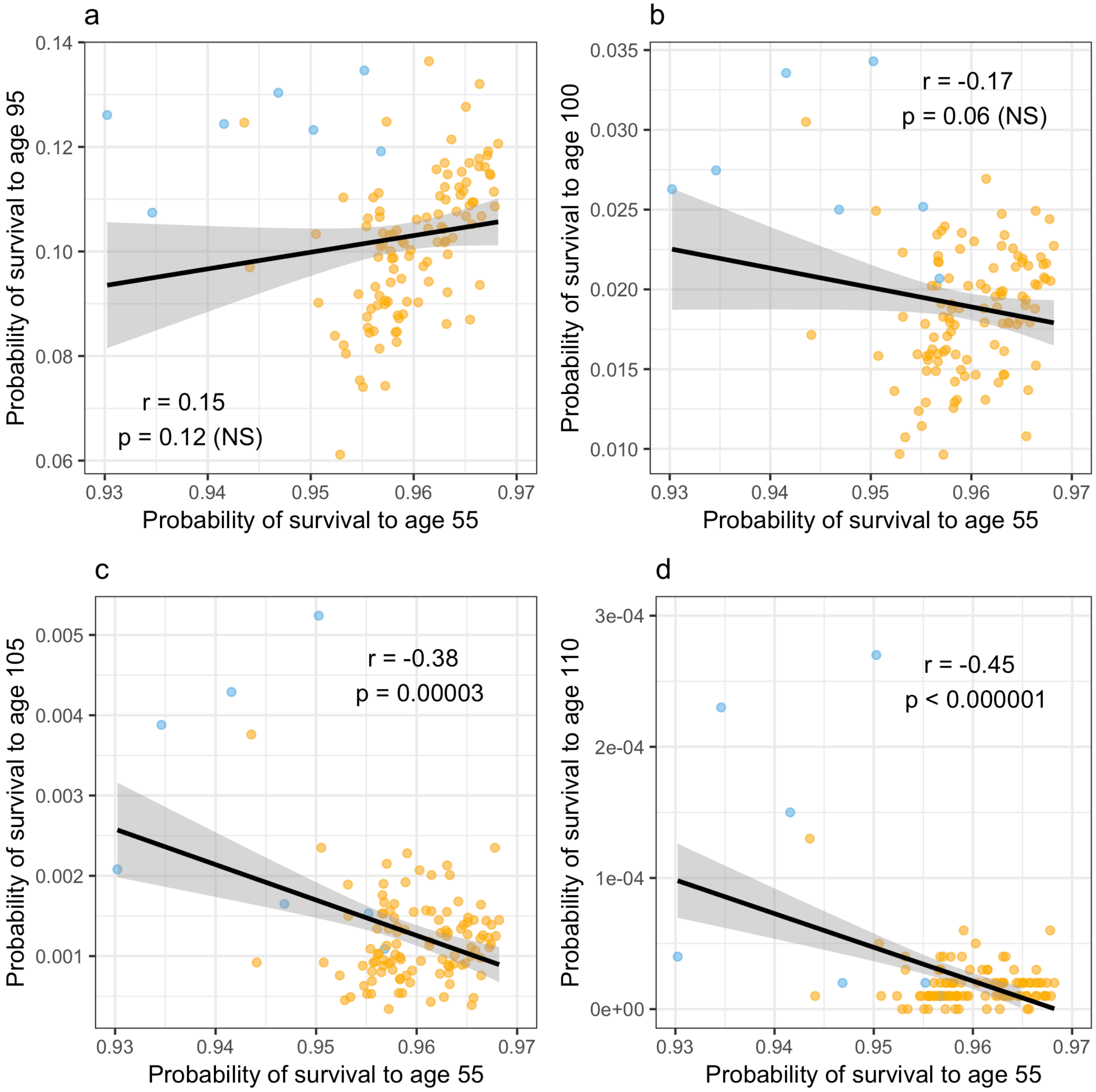


**Figure S2. Relationship between mid-life and late-life survival across Italian provinces.** Across Italian provinces (points), probabilities of survival in mid-life are positively correlated with the probability of survival at older ages until around age 95 (a; r = 0.15; p=0.1; N=116). However, this relationship inverts at advances ages: better mid-life and early-life probabilities of survival, and higher average longevity, are linked to significantly lower probabilities of survival at 100 years (b), 105 years (c), or 110 years (d) of age. Sardinian provinces shown in blue.

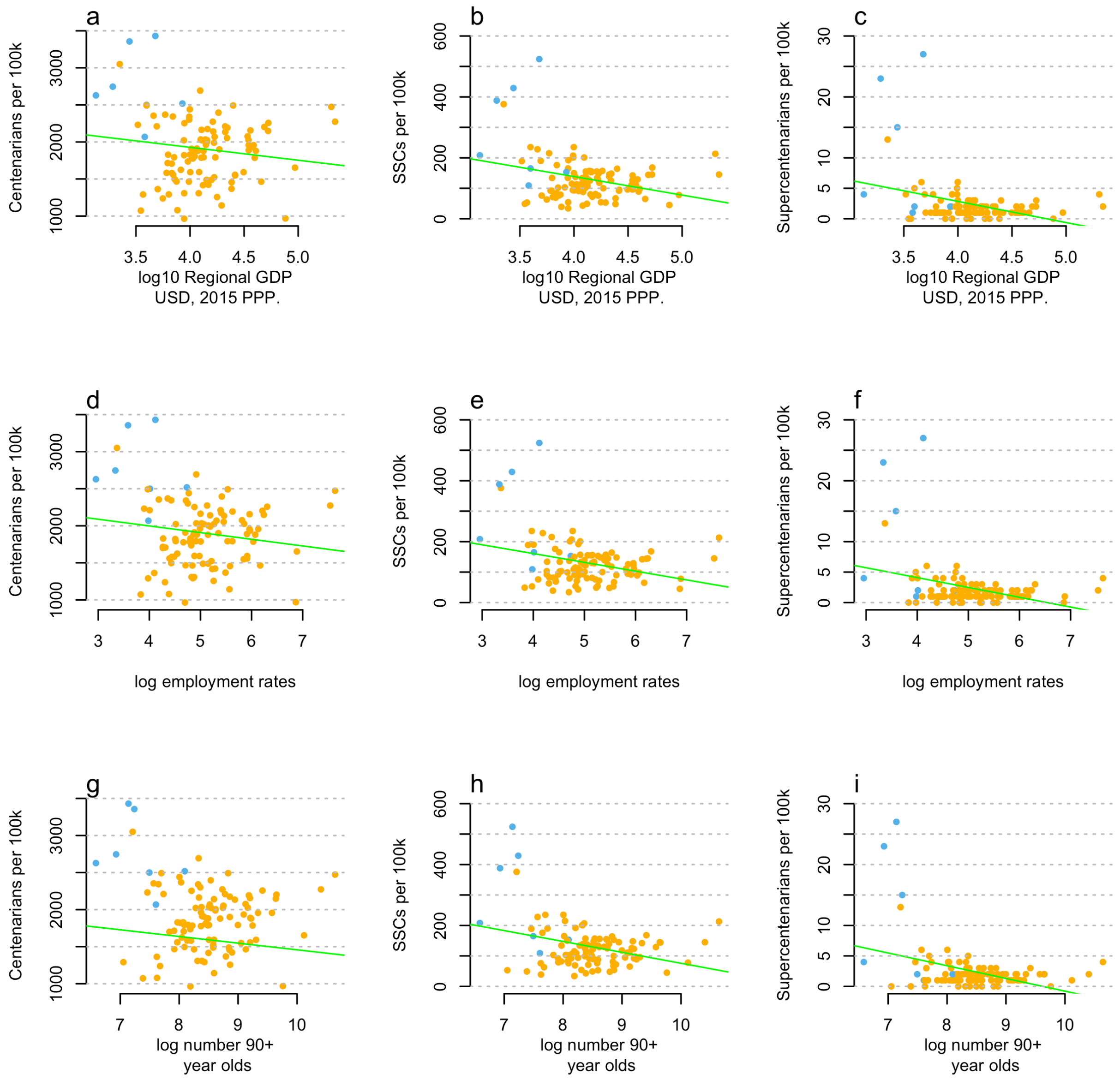


**Figure S3. Italian provinces by GDP per capita and rates of the oldest-old.** According to figures from the Italian national statistics office and regional GDP data from the OECD, purchase-power parity adjusted GDP is negatively correlated with the frequency of (a) centenarians, (b) SSCs and (c) supercentenarians per capita across Italy: a pattern repeated in employment rates (d-f). Furthermore, the total number of 90+ year olds, shown here in log scale, is also negatively associated with the per capita number of centenarians (g), SSCs (h) and supercentenarians (i) across Italy. Linear mixed model regressions shown in green, Sardinian provinces shown in blue.

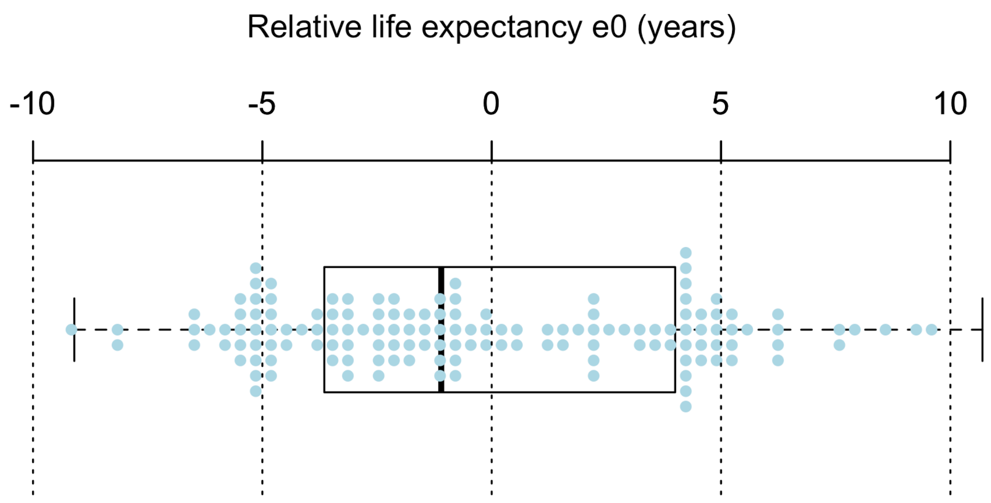


**Figure S4. Historical life expectancy in the home region of birth for French supercentenarians.** Supercentenarians are born in regions with an 84-day shorter life expectancy at birth, a non-significant reduction relative to the national average (one sample t-test NS; p = 0.52; N=143). Comparisons for supercentenarians born in overseas provinces and imperial holdings are unavailable, data show metropolitan France only.

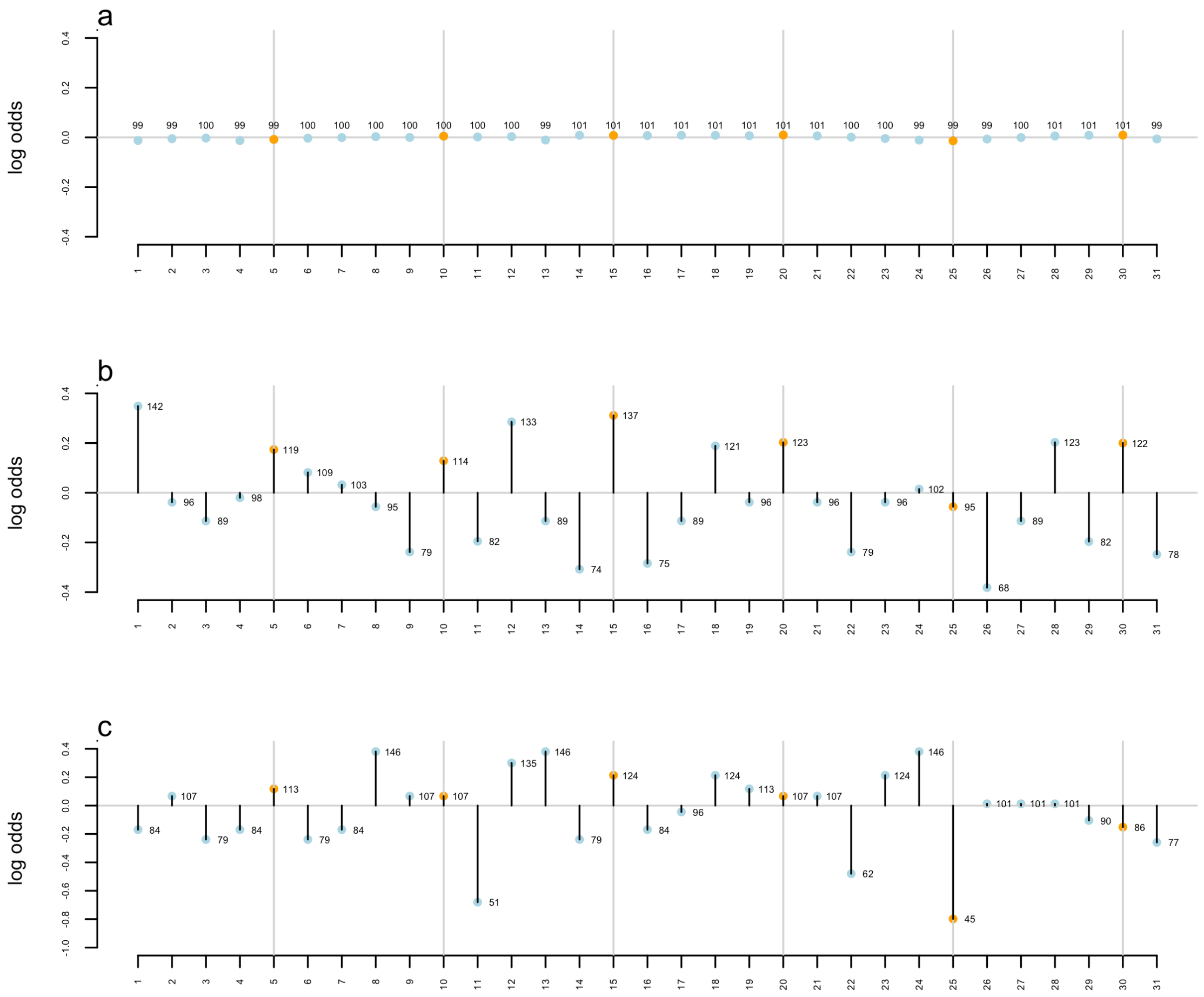


**Figure S5. Age heaping of supercentenarian births.** The distribution of modern birthdates (a), shown here by 70 million US birthdates observed from 1969-1988, display limited variation across days of the month. However, supercentenarian birth dates (N = 1739) are 1.42-fold more likely to be born on the first day of the month and 1.18-fold more likely to be born on days that are multiples of five (orange points) compared with randomly distributed births (b). Age heaping on the first day of the month or in multiples of five is not as clear in the IDL data (c), possibly because of the removal of US birth days and months, or differences in cultural patterns (the 25$^{th}$ is heavily under-represented) and data quality. Points are labeled by the percentage of births over- or under-represented, relative to random sampling.

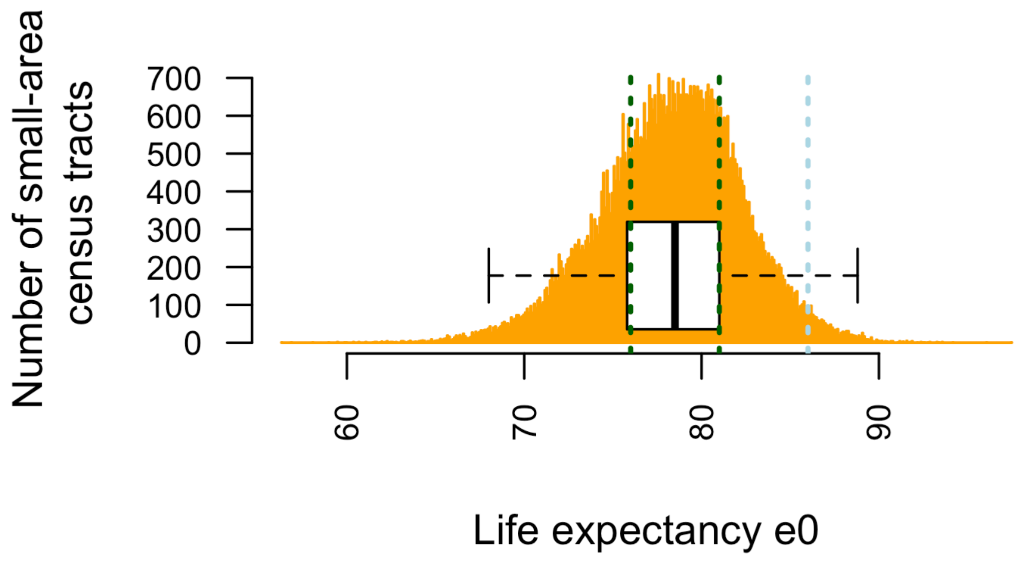


**Figure S6. Distribution of life expectancy estimates in US small-area census tracts, including the Loma Linda 'blue zone'.** When calculated independently by the Centers for Disease Control, the suburbs of Loma Linda range from the 27$^{th}$ to 75$^{th}$ percentiles of life expectancy at birth in the USA (green lines), relative to all other census tracts (orange). The absolute upper estimate, the female-only life expectancy calculated by 'blue zone' proponents (blue line), falls in the 98$^{th}$ percentile of life expectancy: high, but still behind 1401 longer-lived US census tracts and the nations of Japan, Singapore, Monaco, Spain, and South Korea.

**Table S1. Top 20 Italian provinces and regions for supercentenarians per capita.**

| Province | Region | Super-centenarians per 100k ($l_{110}$) | Rank $l_{110}$ | National ranking, 1 = Worst, 116 = Best, ND = No Data: Survival to age 55 | Unemployment | GDP per capita, PPS adjusted | 90+ year-olds per capita (1=fewest) |
|---|---|---|---|---|---|---|---|
| Olbia-Tempio | Sardinia | 27 | 1 | 7 | 14 | 41 | 8 |
| Medio Campidano | Sardinia | 23 | 2 | 2 | 2 | 1 | 33 |
| Carbonia-Iglesias | Sardinia | 15 | 3 | 3 | 4 | 13 | 43 |
| Isernia | Molise | 13 | 4 | 4 | 3 | 34 | 102 |
| Matera | Basilicata | 6 | 5 | 61 | 15 | 17 | 23 |
| Prato | Tuscany | 6 | 6 | 114 | 40 | 80 | 59 |
| L'Aquila | Abruzzo | 5 | 7 | 65 | 39 | 49 | 82 |
| Rieti | Lazio | 5 | 8 | 8 | 7 | 31 | 80 |
| Belluno | Veneto | 4 | 9 | 45 | 29 | 96 | 91 |
| Campobasso | Molise | 4 | 10 | 38 | 22 | 35 | 76 |
| Catanzaro | Calabria | 4 | 11 | 59 | 52 | 26 | 31 |
| Gorizia | Friuli-Venezia Giulia | 4 | 12 | 34 | 10 | 64 | 107 |
| Ogliastra | Sardinia | 4 | 13 | 1 | 1 | 23 | 66 |
| Pistoia | Tuscany | 4 | 14 | 87 | 37 | 59 | 84 |
| Metropolitan City of Rome | Lazio | 4 | 15 | 81 | 111 | 104 | 24 |
| Vibo Valentia | Calabria | 4 | 16 | 12 | 6 | 4 | 38 |
| Brescia | Lombardy | 3 | 17 | 90 | 106 | 94 | 25 |
| Chieti | Abruzzo | 3 | 18 | 64 | 64 | 53 | 71 |
| Cremona | Lombardy | 3 | 19 | 67 | 56 | 75 | 60 |
| Fermo | Marche | 3 | 20 | 35 | ND | ND | ND |

**Table S2. Top 20 French regions for the oldest-old per capita (metropolitan and ranked overseas territories only).**

| Region | Super-centenarians (IDL) | Super-cent. per capita (rank) | Poverty rate ages 75+ (%) | National Rankings 1 = Worst, 101 = Best, ND = No Data | | |
|---|---|---|---|---|---|---|
| | | | | Poverty rate ages 75+ (rank) | Poverty rate | GDP per capita, PPS adjusted |
| Creuse | 3 | 1 | 17.0 | 4 | 16 | 4 |
| Martinique | 8 | 2 | 31.1 | 2 | 2 | 46 |
| Guadeloupe | 8 | 3 | ND | ND | ND | 19 |
| Cantal | 2 | 4 | 13.9 | 10 | 60 | 21 |
| Haute-Loire | 3 | 5 | 11.1 | 26 | 77 | 10 |
| Lozère | 1 | 6 | 14.5 | 7 | 52 | 27 |
| Lot | 2 | 7 | 12.2 | 17 | 46 | 36 |
| Indre | 2 | 8 | 10.5 | 32 | 43 | 22 |
| Yonne | 3 | 9 | 7.0 | 75 | 44 | 39 |
| Aude | 3 | 10 | 14.4 | 8 | 5 | 11 |
| Tarn | 3 | 11 | 11.1 | 25 | 33 | 13 |
| Paris | 17 | 12 | 9.6 | 39 | 27 | 100 |
| Jura | 2 | 13 | 8.5 | 56 | 84 | 34 |
| Maine-et-Loire | 6 | 14 | 6.8 | 83 | 85 | 59 |
| Aveyron | 2 | 15 | 12.6 | 16 | 57 | 42 |
| Hautes-Alpes | 1 | 16 | 9.9 | 38 | 53 | 48 |
| Orne | 2 | 17 | 8.3 | 61 | 26 | 25 |
| Ariège | 1 | 18 | 12.8 | 15 | 14 | 6 |
| Corse-du-Sud | 1 | 19 | 15.9 | 5 | 19 | 75 |
| Alpes-de-Haute-Provence | 16 | 20 | 10.8 | 69 | 123 | 40 |

**Table S3. Top 20 regions in England for the oldest-old per capita.**

| **Region** | **Total SSCs** | **SSCs per capita (rank)** | **National Rankings 1 = Worst, 131 = Best** Income Deprivation Affecting Older People | 90+ year-olds per capita (1=fewest) | Index of Multiple Deprivation | Crime rate | Health index |
|---|---|---|---|---|---|---|---|
| Tower Hamlets | 15 | 1 | 1 | 1 | 26 | 23 | 52 |
| Lewisham & Southwark | 31 | 2 | 8 | 5 | 41 | 26 | 59 |
| Darlington | 5 | 3 | 62 | 85 | 46 | 14 | 32 |
| Lambeth | 15 | 4 | 5 | 3 | 49 | 18 | 56 |
| Camden & City of London | 11 | 5 | 55 | 18 | 90 | 119 | 110 |
| Kensington & Chelsea and Hammersmith & Fulham | 15 | 6 | 16 | 10 | 67 | 37 | 112 |
| Isle of Wight | 6 | 7 | 72 | 128 | 58 | 92 | 61 |
| East Cumbria | 11 | 8 | 120 | 121 | 94 | 130 | 82 |
| Southampton | 10 | 9 | 42 | 42 | 34 | 3 | 30 |
| Portsmouth | 8 | 10 | 44 | 65 | 33 | 10 | 48 |
| Derby | 9 | 11 | 48 | 64 | 40 | 75 | 33 |
| North and West Norfolk | 9 | 12 | 90 | 126 | 64 | 129 | 51 |
| Telford and Wrekin | 6 | 13 | 49 | 21 | 51 | 53 | 38 |
| Manchester | 18 | 14 | 3 | 6 | 4 | 1 | 3 |
| Haringey & Islington | 16 | 15 | 4 | 4 | 28 | 8 | 65 |
| Leicester | 11 | 16 | 7 | 17 | 17 | 25 | 27 |
| Tyneside | 27 | 17 | 29 | 62 | 25 | 39 | 16 |
| Liverpool | 15 | 18 | 6 | 16 | 2 | 12 | 2 |
| Suffolk | 23 | 19 | 101 | 120 | 86 | 91 | 97 |
| East Kent | 16 | 20 | 69 | 123 | 57 | 49 | 64 |

**Table S4. Analysis of variance table and model coefficients for England SSCs.**

| | ANOVA | | | Coefficients | | | |
|---|---|---|---|---|---|---|---|
| Variable | SumSq | F val. | Pr(>F) | Estimate | Std. Error | t val. | Pr(>\|t\|) |
| Intercept | | | | 2.12E-04 | 1.131E-04 | 1.88 | 0.06 |
| IMD*** | 1.3E-09 | 16.54 | 9.9E-05 | -5.76E-06 | 5.114E-06 | -1.13 | 0.26 |
| AIMD*** | 1.6E-09 | 20.34 | 1.9E-05 | -1.58E-03 | 6.741E-04 | -2.35 | 0.02 |
| Health | 1.7E-10 | 2.16 | 0.145 | 1.91E-04 | 1.4E-04 | 1.37 | 0.18 |
| Crime | 5.2E-11 | 0.66 | 0.419 | 1.26E-04 | 2.067E-04 | 0.61 | 0.54 |
| GDP** | 8.1E-10 | 10.27 | 0.002 | -8.09E-09 | 3.674E-09 | -2.20 | 0.03 |
| Employed | 7.9E-12 | 0.10 | 0.753 | -9.76E-10 | 9.974E-09 | -0.10 | 0.92 |
| Nonagenarians_per_capita** | 6.8E-10 | 8.60 | 0.004 | 8.85E-04 | 6.049E-04 | 1.46 | 0.15 |
| IMD:AIMD | 1.7E-10 | 2.19 | 0.142 | 4.78E-05 | 2.777E-05 | 1.72 | 0.09 |
| IMD:Health* | 3.8E-10 | 4.80 | 0.031 | -3.65E-06 | 5.907E-06 | -0.62 | 0.54 |
| AIMD:Health* | 3.7E-10 | 4.63 | 0.034 | -1.30E-03 | 9.172E-04 | -1.42 | 0.16 |
| IMD:Crime* | 3.4E-10 | 4.26 | 0.042 | -1.07E-05 | 8.496E-06 | -1.26 | 0.21 |
| AIMD:Crime** | 6.1E-10 | 7.74 | 0.007 | 5.80E-04 | 1.376E-03 | 0.42 | 0.67 |
| Health:Crime | 7.6E-12 | 0.10 | 0.757 | 4.30E-06 | 1.97E-04 | 0.02 | 0.98 |
| IMD:GDP | 2.0E-12 | 0.03 | 0.873 | 2.57E-10 | 1.716E-10 | 1.50 | 0.14 |
| AIMD:GDP | 8.8E-11 | 1.11 | 0.294 | 5.96E-08 | 2.09E-08 | 2.85 | 0.01 |
| Health:GDP | 1.2E-10 | 1.46 | 0.230 | -6.54E-09 | 4.69E-09 | -1.40 | 0.17 |
| Crime:GDP | 6.8E-12 | 0.09 | 0.769 | -7.07E-09 | 7.241E-09 | -0.98 | 0.33 |
| IMD:AIMD:Health | 7.8E-12 | 0.10 | 0.754 | 2.77E-05 | 3.87E-05 | 0.72 | 0.48 |
| IMD:AIMD:Crime* | 5.4E-10 | 6.84 | 0.010 | 8.56E-06 | 4.919E-05 | 0.17 | 0.86 |
| IMD:Health:Crime | 1.1E-12 | 0.01 | 0.906 | -4.48E-06 | 9.941E-06 | -0.45 | 0.65 |
| AIMD:Health:Crime | 1.4E-11 | 0.18 | 0.674 | 1.16E-03 | 9.983E-04 | 1.17 | 0.25 |
| IMD:AIMD:GDP | 6.5E-11 | 0.82 | 0.367 | -1.93E-09 | 8.711E-10 | -2.22 | 0.03 |
| IMD:Health:GDP | 2.1E-10 | 2.60 | 0.110 | 9.32E-11 | 2.096E-10 | 0.45 | 0.66 |
| AIMD:Health:GDP** | 7.3E-10 | 9.28 | 0.003 | 4.93E-08 | 3.03E-08 | 1.63 | 0.11 |
| IMD:Crime:GDP | 2.6E-11 | 0.33 | 0.569 | 4.18E-10 | 3.142E-10 | 1.33 | 0.19 |
| AIMD:Crime:GDP | 1.0E-12 | 0.01 | 0.911 | 5.52E-09 | 4.53E-08 | 0.12 | 0.90 |
| Health:Crime:GDP | 1.9E-10 | 2.42 | 0.123 | -1.96E-09 | 6.696E-09 | -0.29 | 0.77 |
| IMD:AIMD:Health:Crime | 9.0E-12 | 0.11 | 0.737 | -1.38E-05 | 4.047E-05 | -0.34 | 0.73 |
| IMD:AIMD:Health:GDP | 1.4E-10 | 1.81 | 0.182 | -9.46E-10 | 1.324E-09 | -0.72 | 0.48 |
| IMD:AIMD:Crime:GDP | 1.8E-10 | 2.32 | 0.131 | -8.82E-10 | 1.621E-09 | -0.54 | 0.59 |
| IMD:Health:Crime:GDP | 1.1E-10 | 1.34 | 0.249 | 2.42E-10 | 3.288E-10 | 0.74 | 0.46 |
| AIMD:Health:Crime:GDP | 9.6E-11 | 1.22 | 0.272 | -3.76E-08 | 3.425E-08 | -1.10 | 0.28 |
| IMD:AIMD:Health:Crime:GDP | 9.3E-12 | 0.12 | 0.732 | 3.98E-10 | 1.161E-09 | 0.34 | 0.73 |

IMD = Indices of Multiple Deprivation; AIMD = Old-Age Indices of Multiple Deprivation

**Table S5. Top 20 Japanese prefectures for centenarians per capita in 2015.**

| Prefecture | Centenarians (total) | Rank Centenarians per capita | Poverty rate (%) | National ranking, 1 = Worst, 47 = Best | | |
|---|---|---|---|---|---|---|
| | | | | Poverty rate (rank) | Financial strength | Income per capita 2011 |
| Shimane | 515 | 1 | 16.7 | 30 | 1 | 13 |
| Kochi | 486 | 2 | 23.7 | 4 | 2 | 6 |
| Okinawa | 872 | 3 | 34.8 | 1 | 5 | 1 |
| Kagoshima | 985 | 4 | 24.3 | 2 | 6 | 5 |
| Tottori | 334 | 5 | 18.9 | 21 | 3 | 2 |
| Yamaguchi | 806 | 6 | 16.9 | 27 | 23 | 36 |
| Kumamoto | 972 | 7 | 21.5 | 10 | 16 | 11 |
| Saga | 441 | 8 | 15.6 | 35 | 13 | 10 |
| Toyama | 554 | 9 | 11.2 | 47 | 25 | 44 |
| Okayama | 980 | 10 | 20.6 | 15 | 29 | 17 |
| Ehime | 720 | 11 | 20.2 | 17 | 20 | 12 |
| Miyazaki | 566 | 12 | 23.0 | 6 | 9 | 3 |
| Nagasaki | 700 | 13 | 22.2 | 8 | 7 | 7 |
| Hiroshima | 1395 | 14 | 16.9 | 26 | 35 | 35 |
| Kagawa | 482 | 15 | 17.2 | 24 | 27 | 28 |
| Tokushima | 377 | 16 | 21.8 | 9 | 8 | 33 |
| Niigata | 1105 | 17 | 16.0 | 33 | 19 | 21 |
| Nagano | 1000 | 18 | 15.5 | 36 | 26 | 25 |
| Yamanashi | 392 | 19 | 19.1 | 20 | 18 | 34 |
| Oita | 536 | 20 | 21.3 | 13 | 15 | 19 |

**Table S6. Claims and evidence from the Okinawan Blue Zone.**

| ***Claim*** | **Observed Evidence** | **Rank in 47 prefectures** |
|---|---|---|
| ***"Move Naturally"*** [1] | Body Mass Index[2] | 1st |
| *"The world's longest-lived people don't pump iron, run marathons or join gyms. Instead, they live in environments that **constantly** nudge them into **moving without thinking** about it. They **grow gardens** and don't have mechanical conveniences for house and yard work."* [1] | Percent population that engages in gardening[3] | 45th |
| | Rest and Relaxation Time[3] | 46th |
| ***"Purpose"*** [1] | Suicide rate[4] | 9th |
| *"The Okinawans call it "Ikigai" and the Nicoyans call it "plan de vida;" for both it translates to "why I wake up in the morning." Knowing your sense of purpose is worth up to seven years of extra life expectancy."* [1] | Suicide rate (2019) in over-65s[5] | 4th |
| ***"Plant slant"*** [1] | Consumption of … root vegetables[6,7] | 47th (last) |
| *"**Beans**, including fava, black, soy and lentils, are the cornerstone of most centenarian diets. **Meat—mostly pork**—is eaten on average only **five times per month**. Serving sizes are **3-4 oz.**, about the size of a deck of cards."*[1] | leafy vegetables[6,7] | 47th (last) |
| | pickled vegetables[6,7] | 47th (last) |
| | Taro[6,7] | 47th (last) |
| Note: 3-4oz of meat a serve, five times a month, is **5.1-6.8kg of meat a year**- without fish, shellfish, or other meats, actual consumption is **40kg a year**. | Sweet potato[6,7] | 47th (last) |
| | String beans[6,7] | 44th |
| | Soybean paste[6,7] | 44th |
| | Bean curd[6,7] | 46th |
| *19.1kg/year* | Pork[6,7] | 20th |
| *6.2kg/year* | Beef[6,7] | 23rd |
| *12.0kg/year* | Chicken[6,7] | 46th |
| *2.6kg/year* | 'Other raw meat'[6,7] | 3rd |
| ***"Wine at 5"*** [1] | All (pure) alcohol[8] | 4th (8.8 L/year) |
| *"People in all blue zones (except Adventists) drink alcohol **moderately*** and regularly. **Moderate drinkers outlive non-drinkers.** The trick is to drink 1-2 glasses per day* (preferably Sardinian Cannonau wine), with friends and/or with food. And no, you can't save up all week and have 14 drinks on Saturday."* [1] | Wine[8] | 8th (3.6 L/year) |
| | Beer[8] | 2nd (61.0L/year) |
| ***"Belong"*** [1] | Percent Atheist / No religion[9] | 1st (93.4% atheist) |
| *"All but five of the 263 centenarians we interviewed **belonged to some faith-based community**. Denomination doesn't seem to matter. Research shows that attending faith-based services four times per month will add 4-14 years of life expectancy."* [1] | Religious attendances (top 3 religions)[10] | 45th (third last) |

*Substantially exceeds CDC & NHS guidelines for 'heavy' drinking.

**Table S6 cont.**

| *Claim* | **Observed Evidence** | **Rank in 47 prefectures** |
|---|---|---|
| ***"Loved Ones First"*** [1] | Percent population married[11] | 46th |
| *"Successful centenarians in the blue zones put their families first. This means **keeping aging parents** and grandparents **nearby or in the home** (It lowers disease and mortality rates of children in the home too.). They **commit to a life partner** (which can add up to 3 years of life expectancy) and **invest in their children** with time and love (They'll be more likely to care for you when the time comes)."* [1] | Percentage of households with members 65 and over[4,11] | 46th |
| | Single parent households[11] | 5th |
| | Percent Divorced** – male 50-59[11] | 1st |
| | Percent Divorced** – female 50-59[11] | 14th |
| | Percent never married – women 45-49[11] | 22nd |
| | Percent never married – men 45-49[11] | 14th |
| ***"Right Tribe"*** [1] | While the idea is essentially untestable.. | |
| *"The world's longest lived people chose – or were born into – social circles that supported healthy behaviors, Okinawans created "moais"–groups of five friends that committed to each other for life. Research from the Framingham Studies shows that **smoking, obesity, happiness**, and even **loneliness** are contagious. So the social networks of long-lived people have favorably shaped their health behaviors."* [1] | Body Mass Index[2] | 1st |
| | Suicide rate[4] | 9th |
| | Suicide rate (2019) in over-65s[5] | 4th |
| | Smoking - male[7,11] | 43rd (37.4%) |
| | Smoking - female[7,11] | 23rd (9.7%) |

** The overall divorce rate in Okinawa has remained well above the national average in every measured[4] year since 1911.

**Table S6 Bibliography**
